# TransLeish: Identification of membrane transporters essential for survival of intracellular *Leishmania* parasites in a systematic gene deletion screen

**DOI:** 10.1101/2024.06.21.600025

**Authors:** Andreia Albuquerque-Wendt, Ciaran McCoy, Rachel Neish, Ulrich Dobramysl, Tom Beneke, Sally A. Cowley, Kathryn Crouch, Richard J. Wheeler, Jeremy C. Mottram, Eva Gluenz

**Affiliations:** Institute of Cell Biology, University of Bern, Baltzerstrasse 4, 3012 Bern, Switzerland; Global Health and Tropical Medicine, Instituto de Higiene e Medicina Tropical, Universidade Nova de Lisboa, Rua da Junqueira 100, 1349-008 Lisbon, Portugal; University of Oxford, Sir William Dunn School of Pathology, South Parks Road, Oxford, OX1 3RE, UK; York Biomedical Research Institute, Department of Biology, University of York, York, YO10 5DD, UK; Medawar Building for Pathogen Research, Nuffield Department of Medicine, University of Oxford, Oxford, UK; School of Infection and Immunity, University of Glasgow, Sir Graeme Davies Building, 120 University Place, Glasgow, G12 8TA; James and Lillian Martin Centre for Stem Cell Research, Sir William Dunn School of Pathology, University of Oxford, Oxford, OX1 3RE, UK; Present address: Microbes & Pathogen Biology, The Institute for Global Food Security, School of Biological Sciences, Queen’s University Belfast, Belfast, United Kingdom; Present address: Department of Cell and Developmental Biology, Biocentre, University of Würzburg, Am Hubland, 97074 Würzburg, Germany; Lead Contact

**Keywords:** *Leishmania*, membrane transporters, V-ATPase, bar-seq, CRISPR screen, parasite

## Abstract

For the protozoan parasite *Leishmania*, completion of its life cycle requires sequential adaptation of cellular physiology and nutrient scavenging mechanisms to the different environments of a sand fly alimentary tract and the acidic mammalian host cell phagolysosome. Transmembrane transporters are the gatekeepers of intracellular environments, controlling the flux of solutes and ions across membranes. To discover which transporters are vital for survival as intracellular amastigote forms, we carried out a systematic loss-of-function screen of the *L. mexicana* transportome. A total of 312 protein components of small molecule carriers, ion channels and pumps were identified and targeted in a CRISPR-Cas9 gene deletion screen in the promastigote form, yielding 188 viable null mutants. Forty transporter deletions caused significant loss of fitness in macrophage and mouse infections. A striking example is the Vacuolar H^+^ ATPase (V-ATPase), which, unexpectedly, was dispensable for promastigote growth *in vitro* but essential for survival of the disease-causing amastigotes.

## Introduction

Transmembrane transporters, pumps and channels facilitate the passage of solutes that are otherwise impermeant to lipid bilayers, including sugars, phospholipids, amino acids, ions, drugs and toxins, which need to be shuttled into and out of the cell and distributed across cellular organelles. These transport processes are fundamental to cellular physiology and the maintenance of homeostasis. For intracellular pathogens, they are at the interface between host and microbe, enabling parasitic scavenging of nutrients. Protists of the genus *Leishmania* (Order Kinetoplastida) are parasites with a dixenous life cycle, shuttling between an insect vector and a vertebrate host. During its life cycle, the single-celled parasite assumes morphologically and physiologically distinct forms that enable it to colonise the alimentary tract of blood-feeding female phlebotomine sand flies when taken up in a blood meal and persist in the midgut. The parasites then replicate as extracellular flagellated forms, which migrate to the stomodeal valve and differentiate to metacyclic promastigotes pre- adapted to initiate the infection of a vertebrate host. On deposition in the dermis of a mammal during the blood meal, the metacyclic promastigotes are engulfed by resident phagocytic cells. Inside the maturing phagolysosome, the parasites differentiate to the acid tolerant, which persist in the resulting parasitophorous vacuole (PV) where they slowly replicate (Kloehn et al., 2015) until ingested by another sand fly.

Inside the host cell, principally macrophages, parasite membrane transporters are crucial for meeting two main challenges: Firstly, the parasite must maintain its cellular integrity in an acidic milieu and antimicrobial activities from the macrophage which exposes the parasite to hydrolases, reactive oxygen and nitrogen species. Whilst residing in an environment of ∼pH 4-5, amastigotes need a set of surface membrane transporters adapted to functioning at a low external pH. At the same time, they must manage the flow of protons across their membranes to maintain a cytosolic pH of ∼pH 6.5 (Glaser et al., 1988; Zilberstein et al., 1989) and a membrane potential of −100 mV (Bakker-Grunwald, 1992) involving the activity of proton pumps that use energy from ATP to build up proton gradients across membranes. A cell-surface localised P-type H^+^ ATPase is thought to be the principal regulator of amastigote intracellular pH (Marchesini and Docampo, 2002).

The second challenge for the intracellular amastigote is to scavenge from this environment all its essential micro- and macronutrients to fuel growth and replication. *Leishmania* parasites are auxotrophic for purines, some amino acids, biotin, pterins, folic acid, pantothenate, pyridoxine, riboflavin, nicotinate and heme (McConville et al., 2015; Nayak et al., 2018) and other trace elements, all of which need to be salvaged. Some of the involved transporters have been identified, and some extensively characterised, including multiple transporters for folate/biopterin (Vickers and Beverley, 2011), nucleosides (Alzahrani et al., 2017), iron (Huynh et al., 2006; Mittra et al., 2016), heme (Huynh et al., 2012), magnesium (Zhu et al., 2009) and glucose (Burchmore et al., 2003). The PV contains a varying supply of nutrients derived from host cell digestive processes (peptides, amino acids, ribose, nucleosides, phosphate, sulphate, lipids (Burchmore and Barrett, 2001; McConville et al., 2015). The slow growing amastigote forms in the PV reduce the uptake of carbon sources compared to the rapidly proliferating promastigote forms of *Leishmania* that live freely in a sand fly’s alimentary tract exposed to the blood and sugar meals of the fly, or when grown in a rich laboratory culture medium. Following a metabolic switch, termed stringent response, amastigotes utilise glucose more effectively, compared to promastigotes, without secreting partially oxidized glucose products acetate, alanine or succinate (Saunders et al., 2014). Amastigotes also require an active TCA cycle for *de novo* synthesis of glutamate and glutamine, as transport of these amino acids is down-regulated (Saunders et al., 2014) and they scavenge fatty acids from the PV, which are actively catabolised in the parasite mitochondrion by β-oxidation. Stage-specific metabolic regulation is thought to be achieved at least in part by ubiquitination-dependent degradation of transporter proteins (Vince et al., 2011). To supply the parasite, amastigote transporters must compete with host cell transporters for substrates in the PV lumen before they are exported to the host cytoplasm. For instance, the parasite upregulates its arginine transporter AAP3 to counteract the host cell’s attempt to deprive the pathogen of arginine (Goldman-Pinkovich et al., 2016). Iron is another key micronutrient at the interface between pathogen and host cell. Surface and intracellular iron transporters have been implicated in *Leishmania* parasite differentiation and amastigote survival (Huynh et al., 2006; Mittra et al., 2016; Zhu et al., 2009). Organellar transporters also contribute to *Leishmania’s* virulence, one example being the synthesis of lipophosphoglycan, which depends on nucleotide sugar transporters located in the Golgi (Ma et al., 1997). Finally, membrane transporters are also conduits for the uptake and extrusion of anti-parasitic compounds, with well-characterised examples including uptake of the drug miltefosine via the miltefosine transporter, a phospholipid-transporting P-type ATPase, (Perez-Victoria et al., 2003) and trivalent antimony via the aquaglyceroporin AQP1 (Marquis et al., 2005).

Proteins that mediate transport of substances across lipid bilayers are a structurally and functionally diverse cohort, with dissimilar architectures, ranging from single transmembrane monomers to large multimeric protein complexes. The Transporter Classification Database TCDB (Saier; Saier et al., 2014; Saier et al., 2021) and TransportDB 2.0 (Elbourne et al., 2017) provide a classification system for these proteins based on structure and function. Comparing uncharacterised transporters against TCDB allows for classifications into families but in most cases no definitive prediction of transporter function is possible.

There are just over 30 transporters for which gene deletion attempts have been reported in the literature for any *Leishmania* sp. (see (Jones et al., 2018) and Supplementary Table 1). Many of these studies focused on transporters for sugars, nucleobases, folate, iron and heme and transporters implicated in drug resistance. For the majority of *Leishmania* membrane transporters their contribution to promastigote and amastigote fitness remains untested. Here, we conduct a systematic loss-of-function screen across the whole *Leishmania* “transportome” of 312 putative transporter encoding genes. Using a CRISPR- Cas9 mediated gene deletion and barcoding strategy (Beneke and Gluenz, 2020; Beneke et al., 2017) we successfully generated 188 null mutants and another 81 mutant cell lines that still retained a copy of the target gene. The mutants’ fitness was tested in promastigotes by assessing their growth *in vitro*, and in amastigotes in human macrophages *in vitro* and in a mouse model of infection *in vivo*. This study identified numerous loss-of-fitness and some gain-of-fitness phenotypes in transporter deletion mutants in both parasite forms. It also identified a cohort of mutants that showed conditional essentiality only in amastigotes, with 17 transporter gene deletions exhibiting a significant loss of fitness both in cultured macrophages and mice and another 20 only in mice. These data show that the amastigotes are particularly vulnerable to the loss of proton pumps and several other transporters that merit further study.

## Results

### Identification and composition of the L. mexicana transportome

A combination of database queries and literature searches were performed to collate an inventory of the *L. mexicana* “transportome” comprising channels, carriers and primary active transporters for small molecules, ions and solutes. The *L. mexicana* genome annotation (Shanmugasundram et al., 2023) was searched with relevant keywords and the resulting list was filtered for proteins that had relevant protein domains, sequence similarity to transporter proteins listed in TCDB or experimental evidence of transporter function. Components of transport systems involved in protein translocation, intraflagellar transport, nuclear pores and components of intraorganellar membrane contact sites were excluded from this study. This resulted in a set of 312 putative membrane transporter proteins, constituting approximately 3.8% of the *L. mexicana* proteome (Figure 1, Supplementary Table 1), a value on par with the TransportDB 2.0 predictions for the membrane transporter repertoire in other single- and multi-cellular eukaryotic organisms (e.g. *Homo sapiens* 1.8%; *Caenorhabditis elegans* 2.5%; *Chlamydomonas reinhardtii* 2.8%; *Trypanosoma brucei* 3.9%; *Plasmodium falciparum* 2.3%) (Elbourne et al., 2017). Blast searches of TCDB were conducted and the results used to assign the *L. mexicana* proteins to superfamilies according to the Transport Classification (TC) system (Saier et al., 2014). This identified 49 different superfamilies for which *L. mexicana* has between one and 53 protein members. Examining different modes of transport, we found 34 alpha-type channels (TCDB 1.A; 12 different families) and two beta-barrel porins (TCDB 1.B; mitochondrial porins, also known as voltage-dependent anion-selective channel [VDAC]), 184 proteins of secondary carriers (TCDB 2.A; 30 different families) and 91 proteins of primary active transporters (TCDB 3.A; including 53 ATP-binding Cassette [ABC] proteins, 19 V-type and A-type ATPase [F-ATPase] proteins, and 17 P-type ATPase [P-ATPase] proteins) and one auxiliary transport protein (TCDB 8.A) (Figure 2A, Supplementary Table 1). Note that these numbers count individual proteins, some of which are subunits of multi-protein complexes and include some near-identical copies of gene arrays. For 73% of these gene products, we found published evidence of experimental characterisation in a trypanosomatid species or bioinformatics analysis, giving varying degrees of certainty about their specific functions (Supplementary Table 1). This leaves over one hundred *Leishmania* transporter proteins for which there has been little analysis beyond the assignment to TCDB families. Overall, the specific contribution to the fitness of *Leishmania* parasites has only been tested for a very small number of transporter proteins.

**Figure 1.**
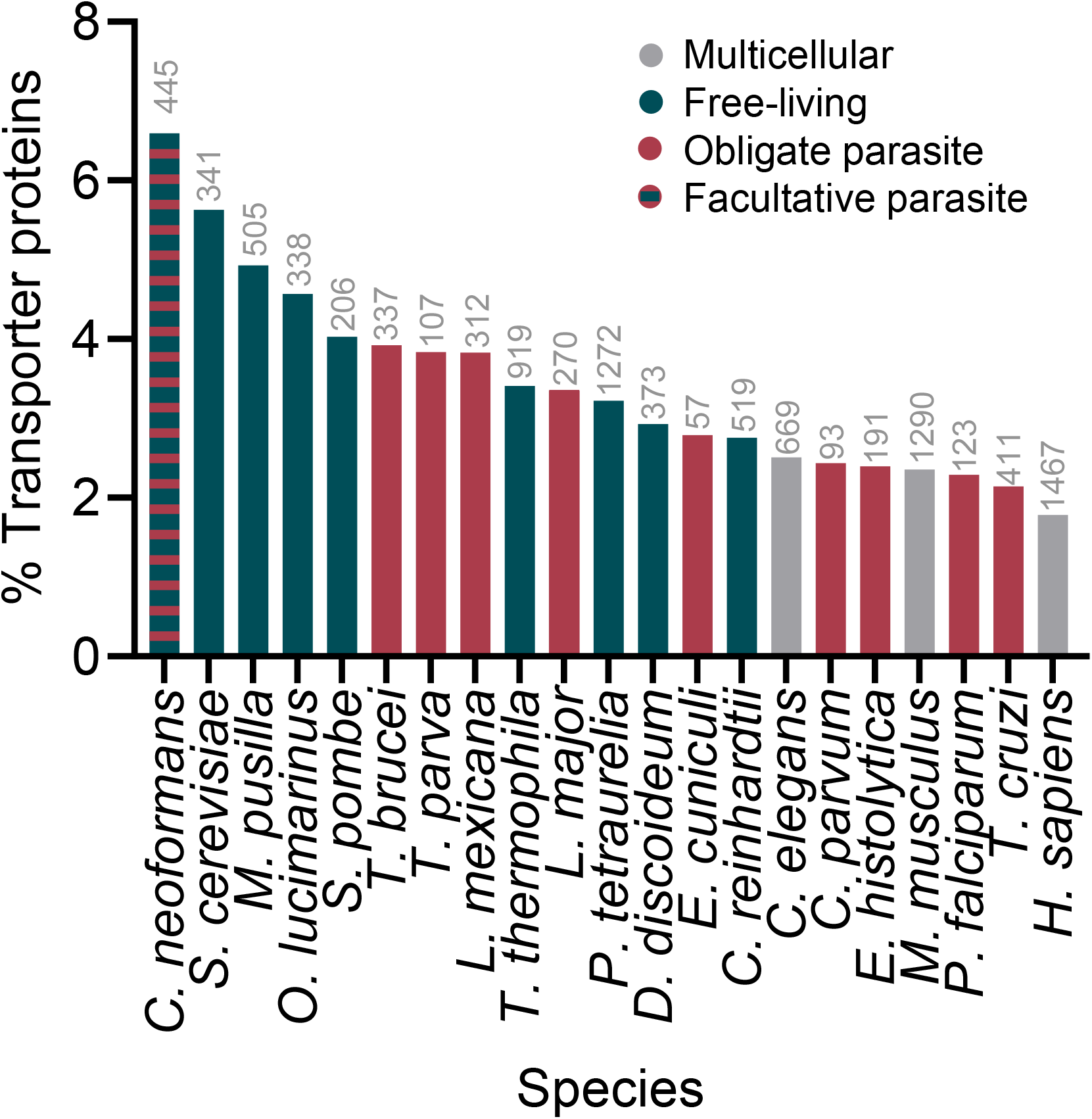
Proportion of membrane transporter proteins in selected eukaryotes. The bars indicate the percentage of membrane transporter proteins in the proteomes of selected parasitic and non-parasitic eukaryotes. The numbers in grey represent the total number of transporter proteins in the respective species (source TransportDB 2.0 (Elbourne et al., 2017)), with the exception of *L. mexicana*, which were determined in this study. The number of proteins in each proteome were extracted from Uniprot (UniProt, 2021), except for *L. mexicana,* where TritrypDB was used as reference(Shanmugasundram et al., 2023).

**Figure 2.**
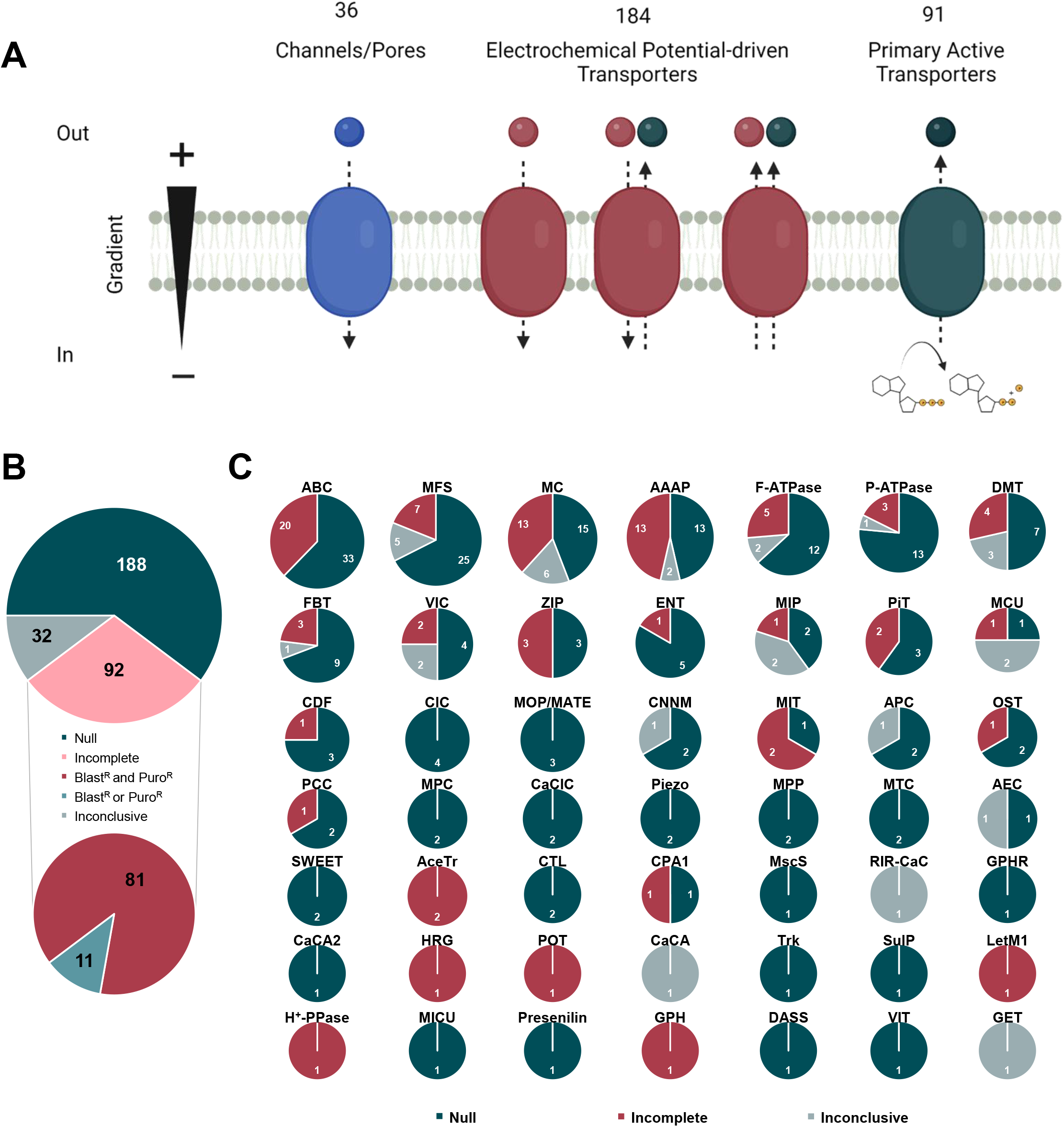
Generation of the TransLeish knock out library. (A) Overview of the number of predicted protein components of channels, carriers and pumps targeted for gene deletion in *L. mexicana*. (B) Summary of gene deletion results. The pie-charts show the numbers of successful deletions (‘null’, dark green), incomplete deletions (light pink), and inconclusive results (light grey). The category ‘incomplete deletions’ contained cell populations that were resistant to both selection drugs (Blasticidin and Puromycin, magenta) and populations that were only recovered from transfections with a single drug marker (Blasticidin or Puromycin, dark grey). (C) Summary of gene deletion results separated into TCDB families.

### Gene deletion yielded 188 null mutants

To study loss-of-function phenotypes, each of these 312 putative transporter genes was targeted for deletion in promastigotes, using a CRSIPR-Cas9-based method for gene replacement by drug-selectable donor DNA cassettes (Beneke et al., 2017), yielding 301 viable mutant populations resistant to both selection drugs (Supplementary Table 1). Diagnostic PCR tests (Supplementary Figure 1) confirmed the loss of the target gene for 188 of these (null mutants) while 81 still retained a copy of the target gene (referred to as ‘incomplete knockouts’) (Figure 2B; Supplementary Table 2). The 11 genes, for which no viable mutants were recovered after at least two attempts, were then targeted using only a single drug resistance cassette (Puromycin). With this approach, 11 mutant cell populations were successfully recovered, providing technical validation for the targeting constructs. As expected, the recovered cell populations had integrated the puromycin cassette, but all had retained copies of the target gene (‘incomplete knockouts’). For 32 cell lines, one or more of the controls in the diagnostic PCRs did not show the expected band and therefore the mutant genotype remained uncertain. For seven cell lines designated as null mutants, whole genome sequencing (WGS) was performed additionally to the diagnostic PCR, to verify the loss of the target gene (including two that had previously been reported to be refractory to deletion) and analyse chromosome ploidy. In all seven genomes, alignment of the Illumina reads from the mutant genome against the reference genome showed precise excision of the target gene in the mutants (Supplementary Figure 2). Chromosome ploidy remained mostly constant in the mutants compared to the parental genome, with a few exceptions (Supplementary Figure 3). None of the ploidy changes affected the chromosome on which the target gene was located. Whether these changes in chromosome ploidy were stochastic events or causally linked to the specific gene deletions cannot be inferred from these data.

Overall, the vast majority of the 188 confirmed deletion mutants generated here are novel and provide a population of viable mutant promastigote forms which could be subjected to phenotype analysis under different conditions.

### Deletion of genes arranged in tandem arrays

A closer examination of the genomic organisation of transporter genes identified 113 genes in 41 arrays of tandemly arranged transporter genes; these included 62 genes in dispersed arrays with similar genes on multiple chromosomes, 19 genes in mixed arrays of transporters belonging to different families and 32 genes in 16 unique arrays of similar genes confined to a single locus (Supplementary Figure 4A; Supplementary Table 3). Since remaining genes could possibly compensate for the loss of individual members from an array, gene deletions strategies were designed to remove whole arrays (Supplementary Figure 4B-D). We selected eight arrays where the deletions of individual genes from the array had all been successful, containing respectively Voltage-gated Ion Channel (VIC), Mitochondrial Carrier (MC), Cyclin M Mg^2+^ Exporter (CNNM), Major Facilitator Superfamily (MFS) and Amino Acid/Auxin Permease (AAAP) family genes. Complete array deletion was confirmed for one MC array (*LmxM.18.1290* and *LmxM.18.1300;* only 30% protein sequence identity) and one AAAP array (*LmxM.34.5350* and *LmxM.34.5360*; 52% protein sequence identity) (Supplementary Table 3, Supplementary Figure 5).

### Transporter protein family encoding genes refractory to deletion

Redundancy could occur at the level of superfamily members beyond gene arrays. Across the 49 superfamilies of membrane transporter encoding genes (Figure 2B), many genes from larger multi- and single copy families were dispensable for *in vitro* promastigote survival. Most of their substrates are not known, and therefore we cannot distinguish in this analysis between transporters whose function is not required in cultured promastigotes and transporters where there is a high level of redundancy between transporters from the same family. For the superfamilies with single members, viable null mutants were generated for nine: a mechanosensitive ion channel (MscS; Δ*LmxM.36.5770*); Golgi pH regulator (GPHR; Δ*LmxM.07.0330*); Ca^2+^:H^+^ antiporter-2 (CaCA2; Δ*LmxM.19.0310*); a K^+^ transporter (Trk; Δ*LmxM.34.0080*); a sulfate permease (SulP; Δ*LmxM.28.1690*); mitochondrial EF hand Ca^2+^ uniporter regulator (MICU; Δ*LmxM.07.0110*); Presenilin (Δ*LmxM.15.1530*), divalent anion:Na^+^ symporter (DASS; Δ*LmxM.28.2930*) and vacuolar iron transporter (VIT; Δ*LmxM.27.0210*), suggesting their functions are dispensable for promastigote growth under standard *in vitro* culture conditions. Conversely, there were six distinct families with only one or two proteins in *L. mexicana* for which no null mutants were obtained, namely the heme-responsive gene protein (HRG; LmxM.24.2230), Proton-dependent Oligopeptide Transporter (POT; LmxM.32.0710), Mitochondrial Inner Membrane K^+^/H^+^ and Ca^2+^/H^+^ Exchanger (Letm1; LmxM.08_29.0920), H^+^-translocating Pyrophosphatase (H^+^-PPase; LmxM.30.1220) and Glycoside-Pentoside-Hexuronide:Cation Symporter (GPH; LmxM.30.0040) or the two Acetate Uptake Transporters (AceTr; LmxM.03.0380 and LmxM.03.0400). These may serve indispensable functions in promastigote physiology.

### Identification of transporter deletions that affect growth fitness of cultured promastigotes

The relative fitness of the viable transporter knock-out mutants was tested next, by combining the mutants into pools and assessing their growth as promastigote forms in standard culture medium. These pools contained 264 barcoded cell lines: five barcoded parental control lines (SBL1-5), 251 mutants from the TransLeish library, Δ*Ros3* (a subunit of the miltefosine transporter, *LmxM.31.0510*) and three non-transporter knock-out mutants with known phenotypes [Δ*dihydroorotase* (*LmxM.16.0580*, slow growth), Δ*IFT88* (*LmxM.27.1130*), very slow growth, no flagellum), Δ*MBO2* (*LmxM.33.2480*), normal growth, uncoordinated movement)], and four non-transporter knock-out mutants with unknown phenotypes (Δ*LmxM.18.0610*, ΔLmxM.*30.2740*, Δ*LmxM.28.2410* and Δ*LmxM.15.0240*). These pools were grown in M199 culture medium, in three separate flasks, for two days. DNA samples were taken at the start of the experiment (0h) and after 24h (Figure 3A) to quantify the abundance of each barcode at each timepoint (Supplementary Figure 6A).

**Figure 3.**
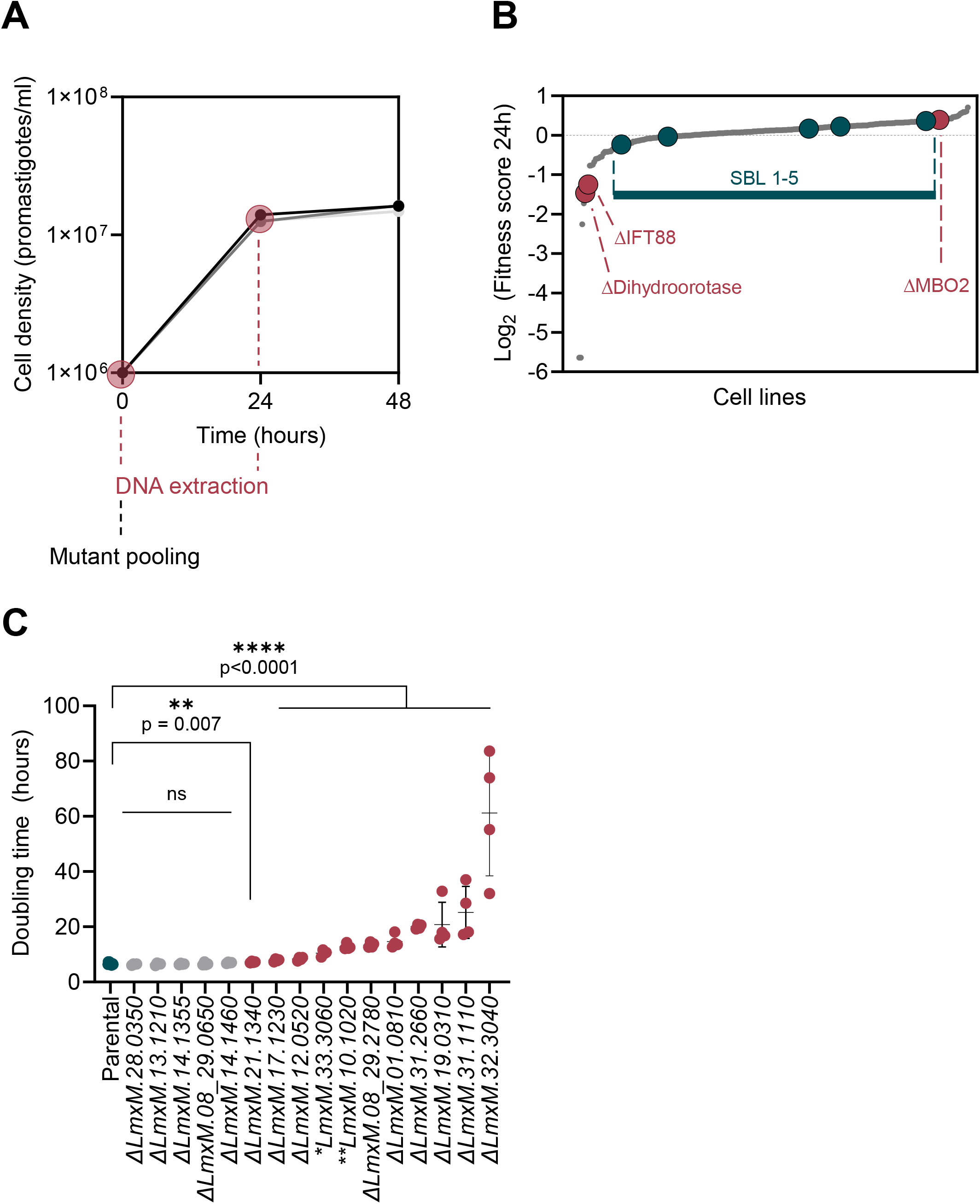
Fitness of promastigote mutants in vitro. (A) Growth curve of the mutant pool. Measurements are shown for each of the three replicate samples. DNA was sampled at the time-points highlighted by a magenta circle. (B) Mutant cell lines ranked in order of their fitness score, from lowest to highest. Green dots, parental control cell lines (SBL1-5); magenta dots, control mutants; small grey dots, TransLeish mutants. (C) Doubling times of the parental control line (green dots), the slow growing control knockout cell line Δ*IFT88* (*LmxM.17.1230*) and selected transporter knockout mutants. Mutants with a significantly increased doubling time are coloured magenta (unpaired t-test). Asterisks indicate mutants where gene deletion was inconclusive (*) or incomplete (**).

Fitness scores were calculated from barcode proportions at a given time point compared to the starting pool (Figure 3B, Supplementary Table 4). None of the mutants showed enhanced fitness in the promastigote *in vitro* pool (score above 2 and p<0.05). Seven mutants showed reduced fitness (score below 0.5 and p<0.05), including the control lines Δ*IFT88* and Δ*dihydroorotase*. Of the transporter null mutants, Δ*MIT1 (LmxM.08_29.2780)*, a mitochondrial pyruvate carrier-like protein deletion (Δ*LmxM.31.1110*) and a putative lysine transporter deletion (Δ*LmxM.31.2660*) showed significantly reduced growth fitness *in vitro*. To test the assumption that the fitness scores were linked to the growth rates of these mutants, their doubling times were measured individually, alongside selected mutants for which slow growth was noticed during the selection process. This confirmed that they grew significantly slower than the parental control lines (Figure 3C). Three null mutants (Δ*ABCI3, LmxM.32.3040;* Δ*CAKC*, *LmxM.01.0810* and Δ*LmxM.34.5150,* a putative biopterin transporter) and three incomplete knockout mutants (*LmxM.10.1020, LmxM.21.1690* and *LmxM.30.0320*) had zero read counts in some or all of the initial pool samples therefore their growth fitness could not be inferred with confidence from the pooled assay. Individual growth rate measurements that were conducted for three of these lines (Δ*ABCI3,* Δ*CAKC* and the *LmxM.10.1020* mutant) showed that their doubling times were significantly longer than for the controls (Figure 3C), which may have led to their underrepresentation in the initial pool. Overall, only a small number of viable mutants showed significant growth defects as promastigotes and most grew at a rate similar to the parental line. This reflects the selection exerted on the mutants following gene deletion, where proliferation in standard laboratory culture medium over several weeks was a necessary condition for the mutants to progress to the pooled screens.

### Forty transporter deletion mutants showed a loss-of-fitness phenotype in vivo

In their life cycle, *Leishmania* parasites experience a profound change in milieu when they enter the mammalian phagocyte and differentiate to the amastigote form. The relative fitness of transporter knock-out mutants in amastigote stages was tested in two models, in a mouse footpad infection model *in vivo*, and human macrophages *in vitro*. For these experiments, the transporter mutants were included in a larger pool of 335 distinct cell lines, comprising of 254 transporter mutants, 61 knock-outs of non-transporter genes, of which 5 served as control cell lines; Δ*IFT88* (*LmxM.27.1130*; very slow promastigote growth, avirulent), Δ*Kharon1* (*LmxM.36.5850*; normal promastigote growth, avirulent, (Tran et al., 2013)), Δ*BBS2* (*LmxM.29.0590*; normal promastigote growth, avirulent), Δ*PMM* (*LmxM.36.1960*; normal promastigote growth, avirulent (Garami and Ilg, 2001)), Δ*GDP-MP* (*LmxM.23.0110*; normal promastigote growth, avirulent (Garami and Ilg, 2001)) and 20 barcoded parental control lines (SBL1-20) mixed into the pool at defined ratios from 1:1 (equal to each mutant) to 1:32 (32 times less). These cells were grown in M199 culture medium for 4 days and then stationary phase parasites were used to infect either human induced pluripotent stem cell derived macrophages (hiPSC-Mac) or BALB/c mice. DNA samples were taken from the pool before infection (0h), from infected hiPSC-Mac at 3h, 24h, 48h and 120h post-infection and from infected mice at 72h, 3 weeks (504h) and 6 weeks (1008h) post-infection (Figure 4A) and a fitness score of the *Leishmania* mutants was calculated for each time point from their barcode proportions (Figure 4B-H, Supplementary Table 4).

**Figure 4.**
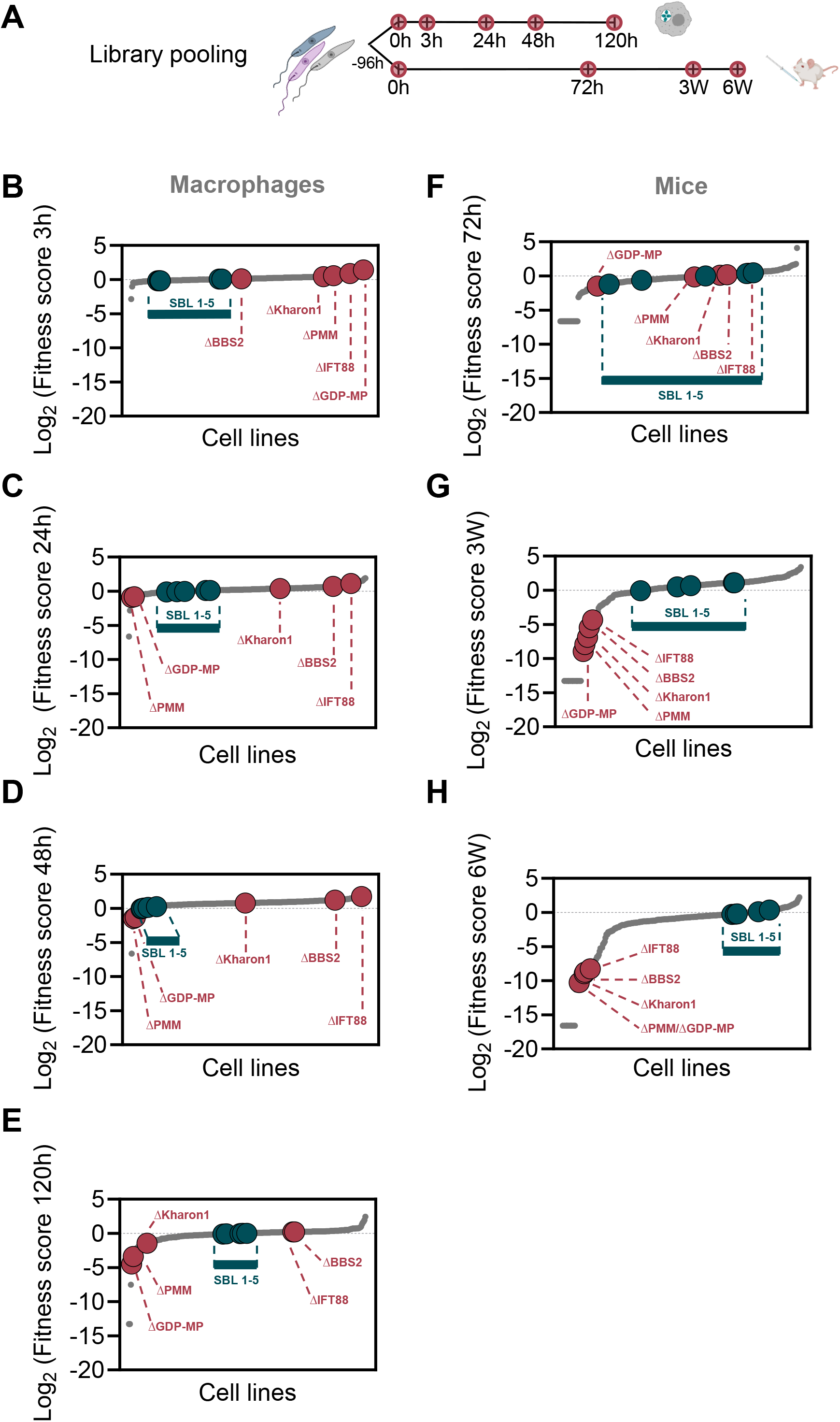
Fitness of intracellular Leishmania, in macrophages and in mice. (A) Overview of the bar-seq timeline for knockout library pooling and infection of hiPSC-Mac and mice. DNA sampling timepoints are marked by magenta circles. (B-E) TransLeish mutant cell lines ranked for each time point in hiPSC-Mac in order of their fitness score, from lowest to highest. (F-H) TransLeish mutant cell lines ranked for each time point in mice in order of fitness score. Green dots, parental controls SBL1-5; magenta dots, control mutants; small grey dots, all TransLeish mutant cell lines.

The fitness scores of the parasite lines tested in mice spanned a larger range compared to the lines tested in macrophage infections *in vitro*. The fitness scores for most of the parental cell lines remained near 1, indicating no change in barcode proportion over time, except for four parental lines seeded in the initial pool at a concentration of 1:16 or lower (Supplementary Figure 7). This indicates where the detection limit for low abundance cell lines is likely to lie.

Overall, the majority of deletions assessed in this screen had little impact on the fitness of *L. mexicana* amastigotes in hiPSC-Mac over the observed 5-day time period, with fitness scores remaining in a range between 0.5 and 2. Looking for mutants that became significantly depleted under the tested conditions, we found the biggest differences at the final time-point (120h), where 28 mutants presented with a fitness score below 0.5 (p<0.05), including three of the control mutants (Δ*GDP-MP*, Δ*PMM*, and Δ*Kharon*) and several proton pump mutants (subunits of the V-ATPase and a P-type H^+^-ATPase). The lowest fitness scores were measured for Δ*ABCI3*, Δ*CACK* and cells lacking a mitochondrial pyruvate carrier-like protein (Δ*LmxM.31.1110*). On the other end of the range, six cell lines presented a fitness score above 2 (p<0.05). Only two of these, Δ*LmxM.32.1860*, lacking the ortholog of the glycosomal transporter GAT2 (Yernaux et al., 2006) and Δ*LmxM.01.0440* (uncharacterised MFS transporter) also showed an enhanced fitness score in samples from 3- week infected mice, albeit without passing the test for statistical significance.

The largest effects were observed in the mouse model *in vivo*. As in the hiPSC-Mac infections, the majority of assessed gene deletions did not alter the fitness of the *L. mexicana* amastigotes in mice, compared to the parental line, but for a substantial minority (16) we measured a severe loss of fitness and for some a gain of fitness. The biggest spread of fitness-values was observed at the final time-point, 6 weeks post inoculation (Figure 4H), where a total of 48 cell lines had a fitness score below 0.5 (p<0.05) and one cell line presented a score above 2 (p<0.05). All five of the control mutants (Δ*GDP-MP,* Δ*PMM,* Δ*Kharon1*, Δ*BBS2* and Δ*IFT88*) showed the expected *in vivo* fitness reduction below 0.5 (p<0.05). A comparison of the data from the hiPSC-Mac and mouse infections identified 17 transporter gene deletion mutants that consistently returned low fitness scores in all samples taken after 120h in macrophages *in vitro*, and 3 weeks and 6 weeks *in vivo* in the mouse footpad and another 11 in both the 3-week and 6-week mouse samples (Figure 5A, B). The Δ*GDP-MP*, Δ*PMM* and Δ*Kharon1* mutants in this group were known to have reduced virulence in amastigotes. The mutant fitness data suggests conditional essentiality for a number of diverse transporters in amastigotes across infection models: the ABC transporters LABCG5 (LmxM.23.0380; shown to be involved in heme salvage in *L. donovani* (Campos-Salinas et al., 2011)), ABCI3 (LmxM.32.3040) and LmxM.33.0670, a calcium/potassium channel (CAKC; LmxM.01.0810), three mitochondrial carrier proteins (MCP2/MCP22 LmxM.31.1110, LmxM.08_29.2780, MCP6 LmxM.33.3060) and an organic solute transporter (OST) Family protein (LmxM.36.6690) and one MFS protein of the PAD surface transporter family (LmxM.30.3170). Strikingly, 11 of the 20 transporter mutants that showed significant loss of fitness after 120h in macrophages and ≥3 weeks in mice, carried deletions of proton pump proteins, namely ten subunits of the Vacuolar H^+^ ATPase (V-ATPase) and one P-type H^+^ ATPase.

**Figure 5.**
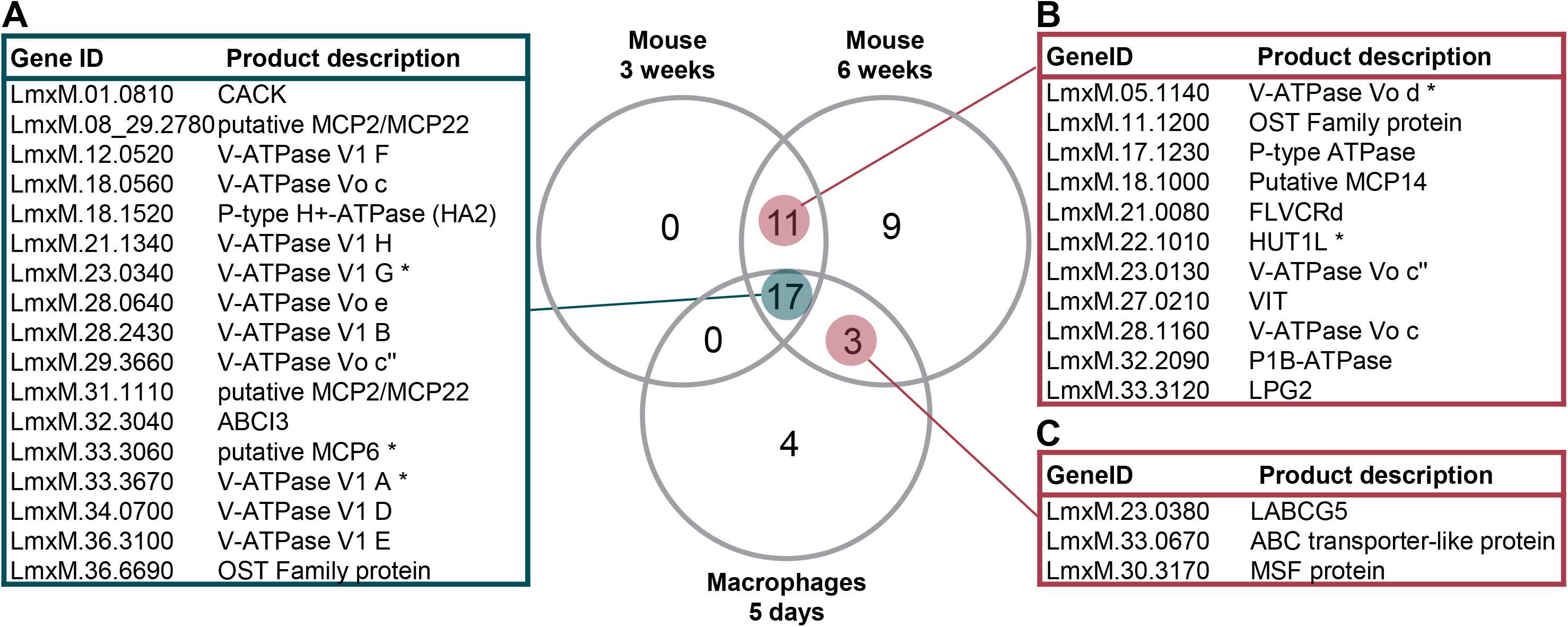
Transporter mutants with significant loss of fitness in macrophages and mice. Venn diagram of transporter mutants depleted in samples from macrophages infected for 5 days and mice infected for 3 and 6 weeks (fitness score below 0.5; p<0.05). (A) Transporter mutants that showed significant loss of fitness in all three conditions. (B) Transporter mutants that showed significant loss of fitness only in mice. (C) Mutants that showed significant loss of fitness at the final time points in macrophages and mice. Asterisks indicate mutants where gene deletion was incomplete.

### L. mexicana V-ATPase pump is essential for survival inside the mammalian host cell and plays a role in pH tolerance

V-ATPases are highly conserved multi-subunit rotary pumps responsible for the acidification of numerous organelles and contributing to cellular pH homeostasis. There are 17 homologues in the *L. mexicana* genome of the conserved yeast V-ATPase subunits, which were all targeted in the TransLeish knockout screen returning twelve confirmed null mutants, three ‘incomplete’ deletions and two inconclusive results (Figure 6A, B). Under promastigote growth conditions, these mutants remained well represented within the pool (Figure 6C). A comparison of the barcode trajectories for all V-ATPase mutants across all time-points in hiPSC-Mac and mice showed consistency: their proportions decreased over time, except Δ*LmxM.31.0920* (Figure 7A, B). Interestingly, LmxM.31.0920 is a homolog of yeast Vph1p, which is one of two isoforms of the V_o_ a subunit that is associated with distinct V-ATPase complexes with different sub-cellular localisations (Perzov et al., 2002). Deletion of the second V_o_, a subunit isoform in *L. mexicana*, Δ*LmxM.23.1510*, resulted in a significantly reduced fitness score in macrophages and mice.

**Figure 6.**
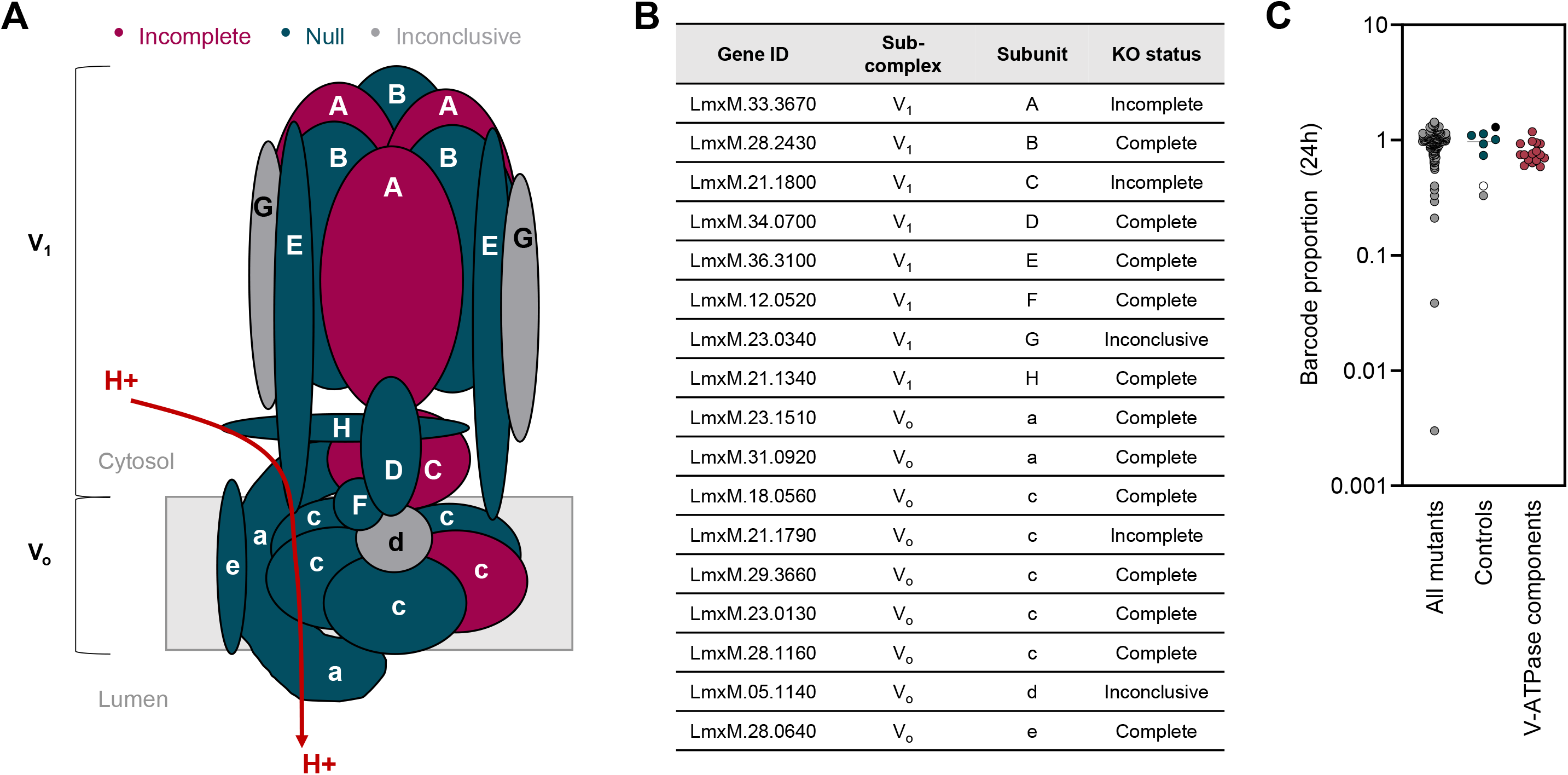
L. mexicana V-ATPase mutants are viable as promastigotes. (A) Schematic of V-ATPase components identified in *L. mexicana*, coloured by the genotype of the corresponding mutants: green, null; magenta, incomplete; grey, inconclusive. (B) Gene IDs of *L. mexicana* V-ATPase subunits. (C) Barcode proportions of mutants after 24h of exponential growth in culture, plotted separately for the following categories: all non-V- ATPase mutants in the TransLeish library (‘All mutants’), dark grey dots; barcoded parental controls (SBL1-5), green dots; Δ*dihydroorotase*, light grey dot; Δ*IFT88,* white dot; Δ*MBO2*, black dot); V-ATPase components, magenta dots.

**Figure 7.**
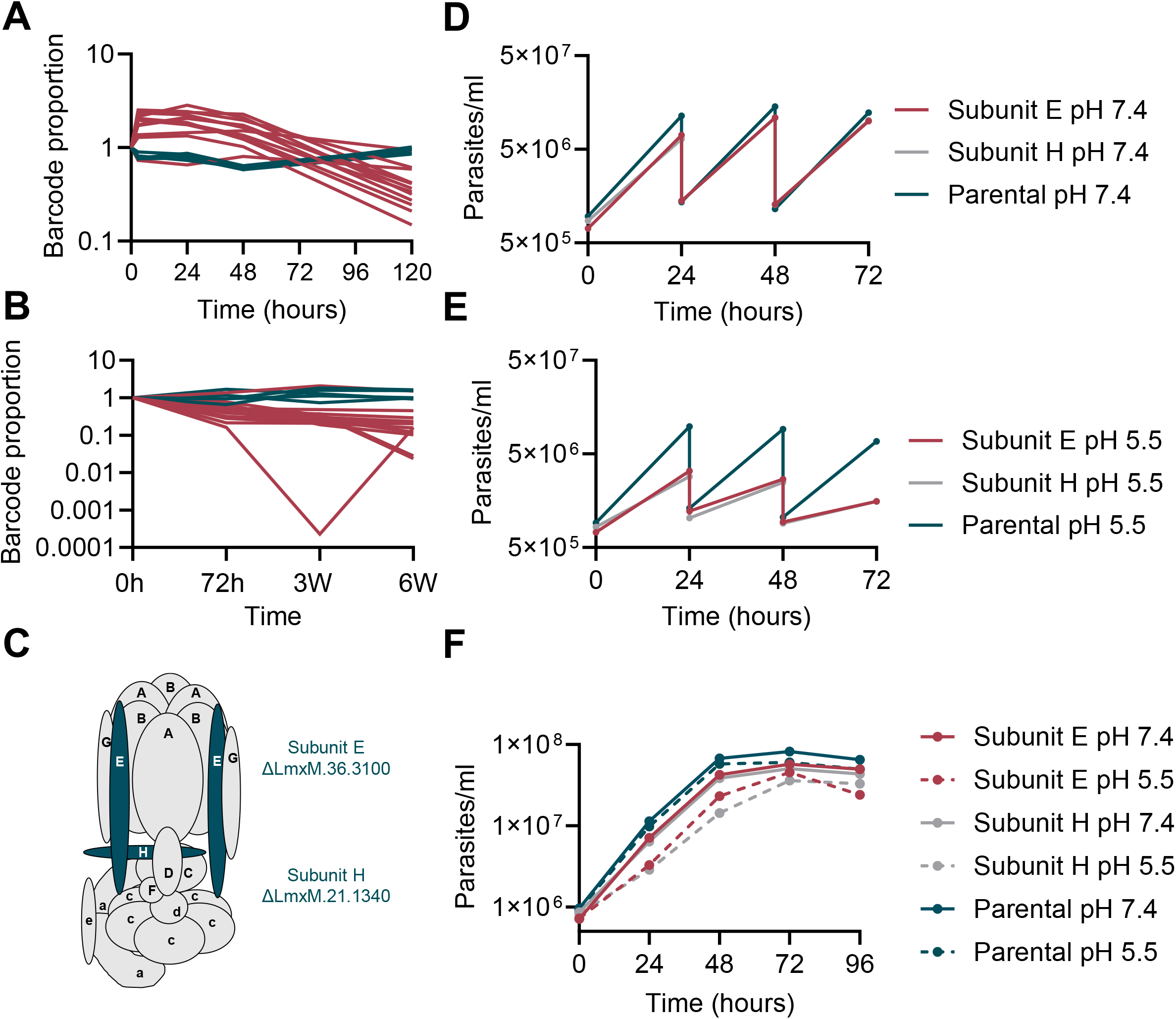
V-ATPase mutants show significant loss of fitness in macrophages and mice and reduced growth rates under low external pH conditions. (A) Barcode trajectories of barcoded parental controls (green) and V-ATPase knockout mutants (magenta) in macrophages. (B) Barcode trajectories in mice; colours as in A. (C) Cartoon of V-ATPase showing the positions of the V_1_ E and V_1_ H subunits. (D) Growth of promastigote form V-ATPase deletion mutants V_1_ E and V_1_ H and the parental cell line in standard M199 medium at pH 7.4. Cells were diluted into fresh medium to a density of 1×10^6^ cells/ml every 24h. Each point represents the average from three separate cultures. (E) Growth profiles as in D, in M199 medium adjusted to pH 5.5. (F) Growth profiles as in D and E, without dilution of the cells into fresh medium. Continuous line, growth at pH 7.4; dotted lines, pH 5.5.

A detrimental effect of V-ATPase gene deletions is compatible with their crucial role in regulating cellular pH in eukaryotic cells. What was surprising however was the discovery that loss of most V-ATPase subunits was compatible with promastigote viability and the deletions had only minor effects on promastigote growth in the pooled screen.

Since promastigotes were cultured at neutral pH while amastigotes reside in an acidic milieu of pH 5.5 or below (Antoine et al., 1990), we hypothesized that the V-ATPase was conditionally essential in low external pH conditions. To assess pH tolerance of the parental cell line and the mutants, the growth rates of the parental cell line and two separate mutants, lacking V-ATPase subunits V_1_ E or V_1_ H, respectively (Figure 7C), was compared in standard M199 medium at pH 7.4 and in M199 medium adjusted to pH 5.5. These experiments were conducted at 27°C, which prevents full differentiation to axenic amastigotes (Bates et al., 1992). In one set of experiments, cells were passaged into fresh medium every 24 hours (Figure 7D, E), in the second they were left to grow in the same flasks for 4 days continuously (Figures 7F). At pH 7.4, doubling times ranged from 6.73 – 8.30h, with little difference between the mutants and the control. At pH 5.5 the parental cell line maintained a doubling time ranging from 7.03 – 8.90h. By contrast, the doubling time for both mutant cell lines progressively increased with each dilution in the acidic medium, from 7.03 to 32.63h.

Together, these data suggest that the *L. mexicana* V-ATPase pump is required to maintain normal cellular physiology under low external pH conditions and consequently, amastigote survival inside the acidic parasitophorous vacuole is compromised in the deletion mutants.

## Discussion

Membrane transporter proteins are cellular gatekeepers for substance exchange between the cell and its environment and between membrane bound subcellular structures and the cytoplasm. They are thus fundamental for maintaining cellular homeostasis across contrasting microenvironments experienced by *Leishmania* parasites throughout their life cycle. In this study we tested for the first time the relative importance of 188 individual transporters in two distinct life cycle stages of this protozoan and discovered over forty transporter-encoding genes whose loss significantly reduced the fitness of intracellular amastigote forms in one or all of the tested conditions.

The vast majority of transporter null mutants generated in this study have never been reported before. Using the CRISPR-Cas9 method allowed for successful knockout of two genes that had been reported to be refractory to gene deletion by sequential homologous recombination in *L. amazonensis* and *L. major*, respectively, namely the mitochondrial iron transporter (MIT1, LmxM.08_29.2780; (Mittra et al., 2016)) and the mitochondrial ABC transporter ABCI3 (LmxM.32.3040; (Arcari et al., 2017)). Individual characterisation of these null mutants revealed a significant increase in promastigote doubling time, consistent with important functions for these transporters. It remains to be tested what compensatory mechanisms, if any, operate in these *L. mexicana* mutants to sustain vital functions. Our results further corroborate in *L. mexicana* previous findings from other *Leishmania* spp. that deletions of the *Leishmania* iron regulator 1 (LIR1; LmxM.21.1580) and the ferrous iron transporter (LIT1; LmxM.30.3060) are viable in promastigotes. In agreement with previous studies, we were unable to generate gene deletions for a porphyrin transporter FLVCRb (LmxM.17.1430 (Cabello-Donayre et al., 2020)) and heme transporter (LHR1; LmxM.24.2230 (Huynh et al.)), underlining the crucial importance for heme salvage in this parasite that lacks all but the last three enzymes required for heme biosynthesis (Koreny et al., 2013). It was however possible to generate viable LABCG5 (Δ*LmxM.23.0380*) null mutants, for a transporter reported to facilitate salvage of heme released from internalized haemoglobin (Campos-Salinas et al., 2011). Also refractory to deletion were the ABC transporter ABCB3 (LmxM.31.3080 (Martinez-Garcia et al., 2016)) and magnesium transporter 2 (MIT/LmxM.25.1090 (Zhu et al., 2009)) supporting previous reports that these may be essential genes.

There was a small number of genes for which knockouts had previously been reported that did not yield null mutants in this present study, encoding aquaglyceroporin 1 (*AQP1; LmxM.30.0020*), the iron/zinc permease (*LmxM.30.3070*) and the equilibrative nucleoside transporter (*BTN1; LmxM.22.0010*). Some of these failures may arise from technical limitations of this screen: For genes in tandem arrays, high sequence similarity led to inconclusive diagnostic PCRs, notably for the glucose transporter array (LmGT1-LmGT3) that was previously deleted in *L. mexicana* (Burchmore et al., 2003; Rodriguez-Contreras et al., 2007). Similarly, for the tandem array of proline/alanine transporters AAP24, shown to be dispensable in *L. donovani* (Inbar et al., 2013), deletion of one *L. mexicana* ortholog (*LmxM.10.0715*) was confirmed, but not for the second (*LmxM.10.0720*). The ABCG1 and ABCG2 array was demonstrated to be dispensable in *L. major* (Manzano et al., 2017), yet gene attempts to delete the orthologs in *L. mexicana* (*LmxM.06.0080* and *LmxM.06.0090*) returned only ‘incomplete’ knockouts. Underestimation of gene copy number could also hamper deletion attempts. Although, using our double-drug selection strategy we were able to successfully delete 11 of the 35 transporter genes present on supernumerary chromosomes 3, 16, and 30; for the remainder a deletion could not be confirmed. Additional rounds of gene replacement with additional drug selection markers might have resulted in additional successful null mutant isolations. Taken together, these results identify at least 188 transporter proteins that are dispensable in cultured *L. mexicana* promastigotes, i.e. 60% of the transportome. Since highly similar gene sequences could have led to false positive diagnostic PCR results, and gene copy number variation poses technical challenges, the true number of dispensable proteins may even be higher, and it is important not to mistake a result of ‘incomplete’ knockout as definitive evidence for gene ‘essentiality’ without further experiments.

Despite this caveat, the essential ‘transportome’ for promastigote *in vitro* must be a sub-set of the 40% of genes for which no null mutants were obtained. In the closely related kinetoplastid, *T. brucei*, growth fitness was assessed genome-wide for a library of RNAi knockdown cell lines grown as procyclics or bloodstream forms *in vitro* (Alsford et al., 2011). This included the 300 orthologs of the *L. mexicana* transporters identified in our study. For 102 (33%) of the *T. brucei* transporter orthologs, induction of RNAi resulted in cell line depletion in at least two of the tested conditions (≥1.5-fold change compared to the uninduced control; data viewed on TritrypDB.org (Shanmugasundram et al., 2023)). Further systematic comparisons of transporter repertoires and mutant fitness across different kinetoplastid species and life cycle stages could support the identification of transporters that are universally required for kinetoplastid cell functions, and those that may be vital for the disease-causing life cycle stages. A systematic analysis of the transporter repertoire and essentiality in *T. cruzi*, which inhabits a different intracellular niche compared to *Leishmania* would be particularly interesting. Amongst other protozoan parasites, a combined analysis of all available reverse genetic experiments on *Plasmodium falciparum* and *P. berghei* concluded that 78% of *Plasmodium* sp. transporters were essential at some point in its life cycle (Martin, 2020). Similar focused analyses of existing gene deletion screens of other intracellular parasites, e.g. *Toxoplasma* (Sidik et al., 2016) and expanding phenotype analyses to more *in vivo* situations and varying environmental conditions will be key to unpicking the conditionally essential transporter repertoire for all of these parasite species in each of their forms.

The phenotyping of amastigote forms showed a dramatic depletion of proton pump mutants in samples taken from macrophages and mice. Only one of the P-type H^+^ATPase genes, HA2, could be deleted in promastigotes, where no growth defect was apparent, but the mutants showed a severe fitness reduction in amastigotes. In *L. donovani*, HA1 was shown to be constitutively expressed while HA2 transcripts were expressed predominantly in amastigotes (Meade et al., 1989) supporting the notion of stage-specific functions for these genes. In both cases it is likely they function as typical single-subunit P-type H+-ATPases, which are found on the surface of plants, fungi and protists, but not mammalian cells.

The V-ATPases are highly conserved multi-subunit complexes whose proton pumping activity serves a variety of cellular functions, mostly leading to acidification of cell organelles, but occasionally also acting at the cell surface membrane of e.g. mammalian osteoclasts or tumour cells, reviewed in (Eaton et al., 2021). Protection against acid stress is a well-documented function for the V-ATPase of yeast, which emerged in screens of mutant libraries as amongst the key determinants in tolerance of strong and weak acids (Johnston et al., 2020; Mira et al., 2010). In the apicomplexan parasite *Toxoplasma*, the V-ATPase serves a dual role. In extracellular parasites, the V-ATPase at the parasite’s plant-like vacuole protects the cells against ionic and osmotic stress, while intracellular parasites were found to position it at the plasma membrane, suggesting it may serve to pump protons out of the parasite cell to protect from acid stress (Stasic et al., 2019). Individual characterisation of the two *L. mexicana* mutants lacking subunits E and H (Δ*LmxM.30.3100* and Δ*LmxM.21.1340*, respectively), showed that in neutral pH, no significant differences in growth were observed when compared to the parental control but tolerance to acidic pH was much reduced. These results point to a protective role of the *Leishmania* V-ATPase in acidic environments. Whether the V-ATPase dynamically relocates to the cell surface in *Leishmania*, either constitutively or uniquely in amastigotes, remains to be tested.

While acid sensitivity alone could explain the demise of amastigote V-ATPase mutants, there are indications that the role of the V-ATPase is more complex. Tolerance of stress conditions other than acid stress may require a functioning V-ATPase. Further detailed studies will be required to dissect the phenotype of the V-ATPase. One of the cellular processes requiring ATPase is autophagy (Collins and Forgac, 2020), which is involved in the differentiation from *Leishmania* promastigotes to amastigotes (Williams et al., 2006). Abolishing V-ATPase function may therefore already block parasite development before the cells are immersed in the acidic luminal contents of the PV, which could provide an alternative explanation for the depletion of these mutants *in vivo*. Endocytosis is yet another process for which kinetoplastid V-ATPases are important, as studied in *T. brucei* (Baker et al., 2015). Assuming *Leishmania* V-ATPase has a similar function, impeding endocytosis, could also render the amastigotes less virulent. The fact that promastigote *L. mexicana* V-ATPase knockout mutants are viable allows for follow-up experimental dissection of these phenotypes.

The exquisite sensitivity of amastigotes towards the loss of proton pump function also marks them as candidates for potential drug targets to combat PV-dwelling pathogens. Encouraging results were obtained with Bafilomycin B1, a macrolide inhibitor of V-ATPase, which had an IC_50_ of <1nm against *L. donovani* and *T. cruzi* amastigotes (Annang et al., 2015), indicating amastigote themselves rely on a working V-ATPase, consistent with our genetic evidence. Modulation of the PV environment via host-cell dependent processes could offer additional routes to parasite killing. De Muylder and co-workers (De Muylder et al., 2016) showed that administration of the µ-opioid receptor antagonist naloxonazine to *Leishmania* infected macrophages inhibited intracellular parasite growth. This was linked to an upregulation of the host V-ATPase and increased volume of acidic vacuoles in the host cell. Combined administration of naloxonazine and the V-ATPase inhibitor concanamycin A restored normal infection levels.

We hypothesize that the P-type H^+^ATPase and V-ATPase are both required for tolerance of an acidic extracellular milieu. Our working model is that the P-type H^+^ATPase is primarily responsible for maintaining cellular pH homeostasis in response to acidified environment by acting as a proton pump at the cell surface. The V-ATPase likely has more varied functions, and possibly multiple cellular locations. Besides the well-characterised activities of bafilomycins and concanamycin A, which inhibit all known eukaryotic V-ATPases, there are other inhibitors specific to certain species or isoforms (Bowman and Bowman, 2005). Further optimisation may allow for targeted perturbation of host and/or parasite functions to eliminate the parasites from their intracellular niche.

## Materials and Methods

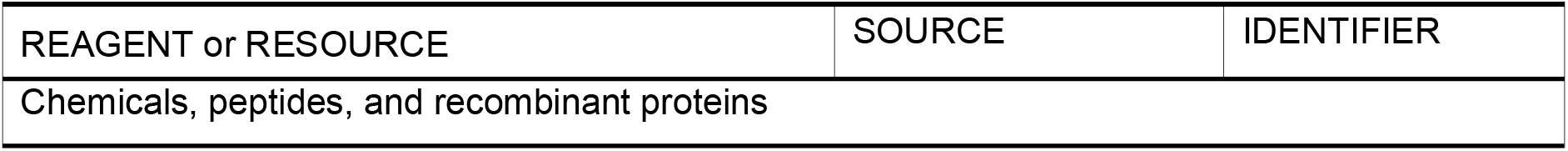
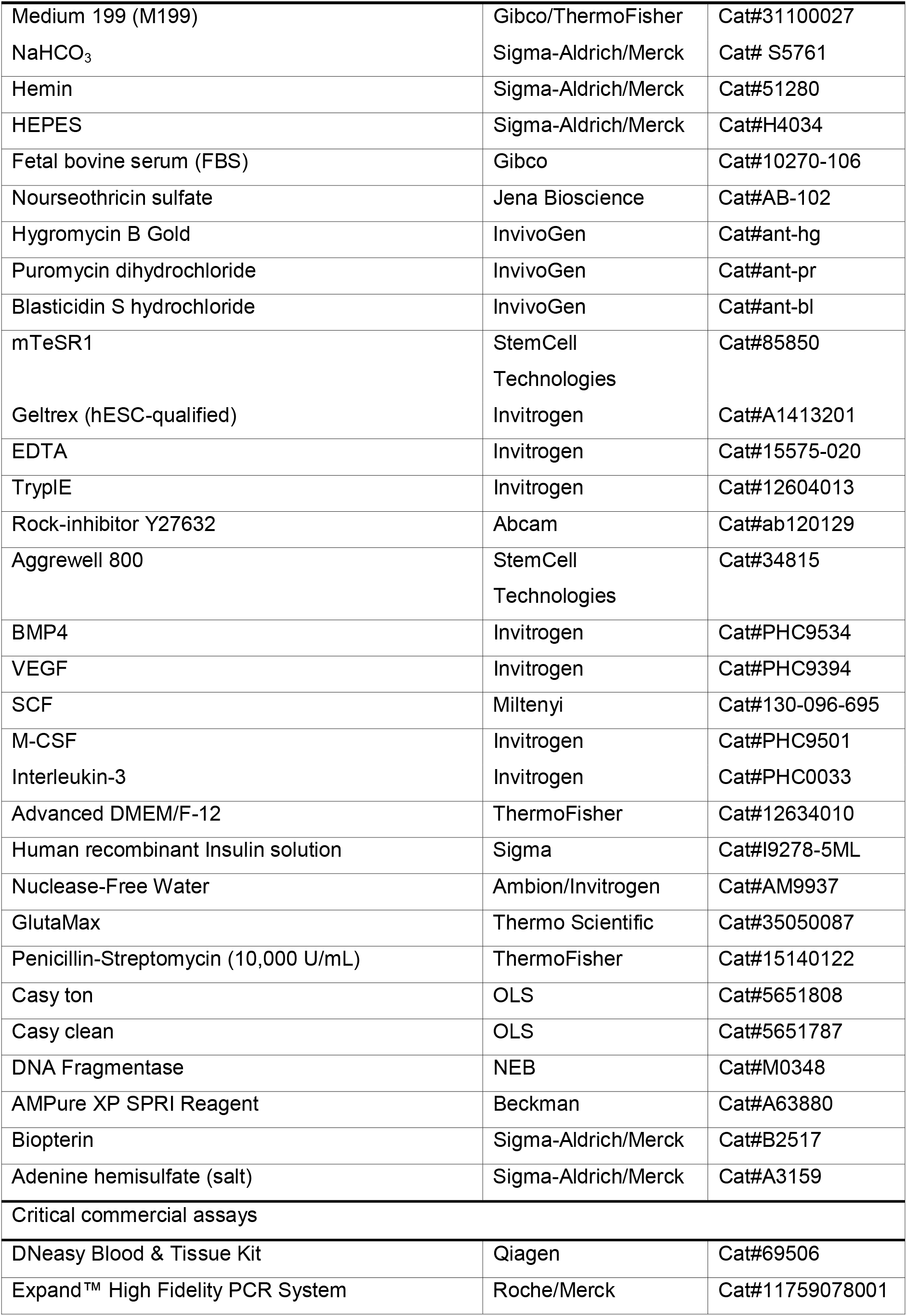
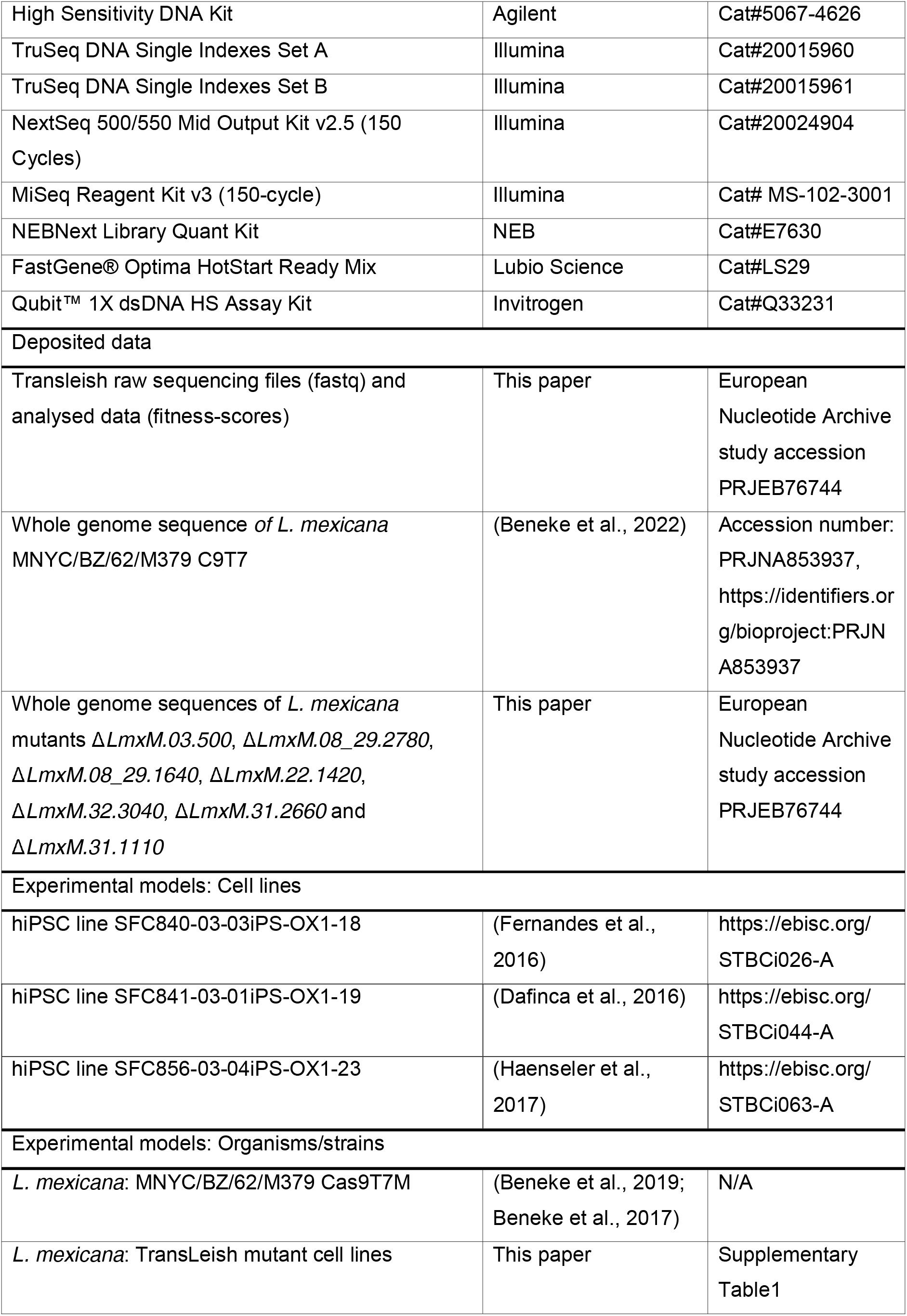
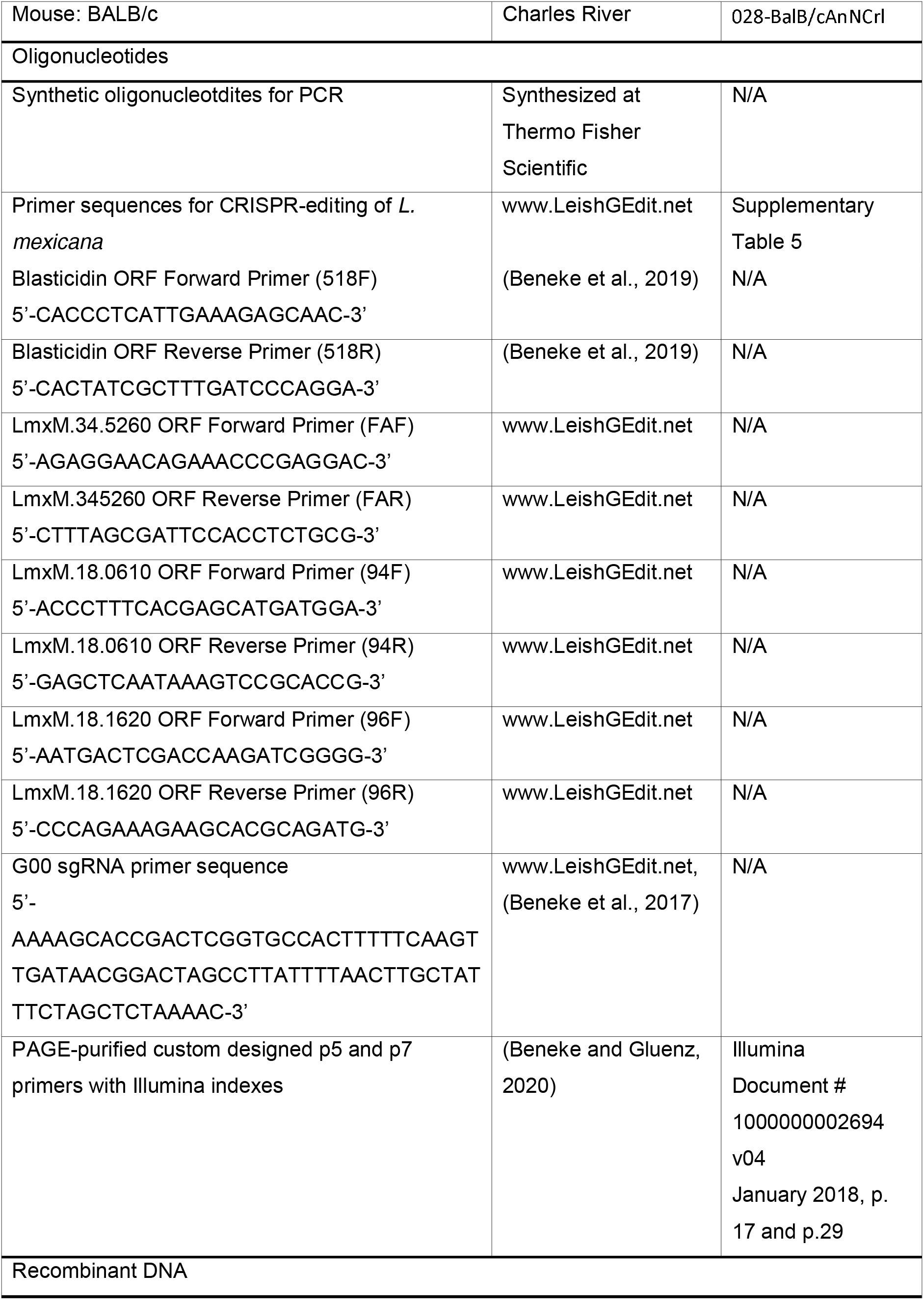
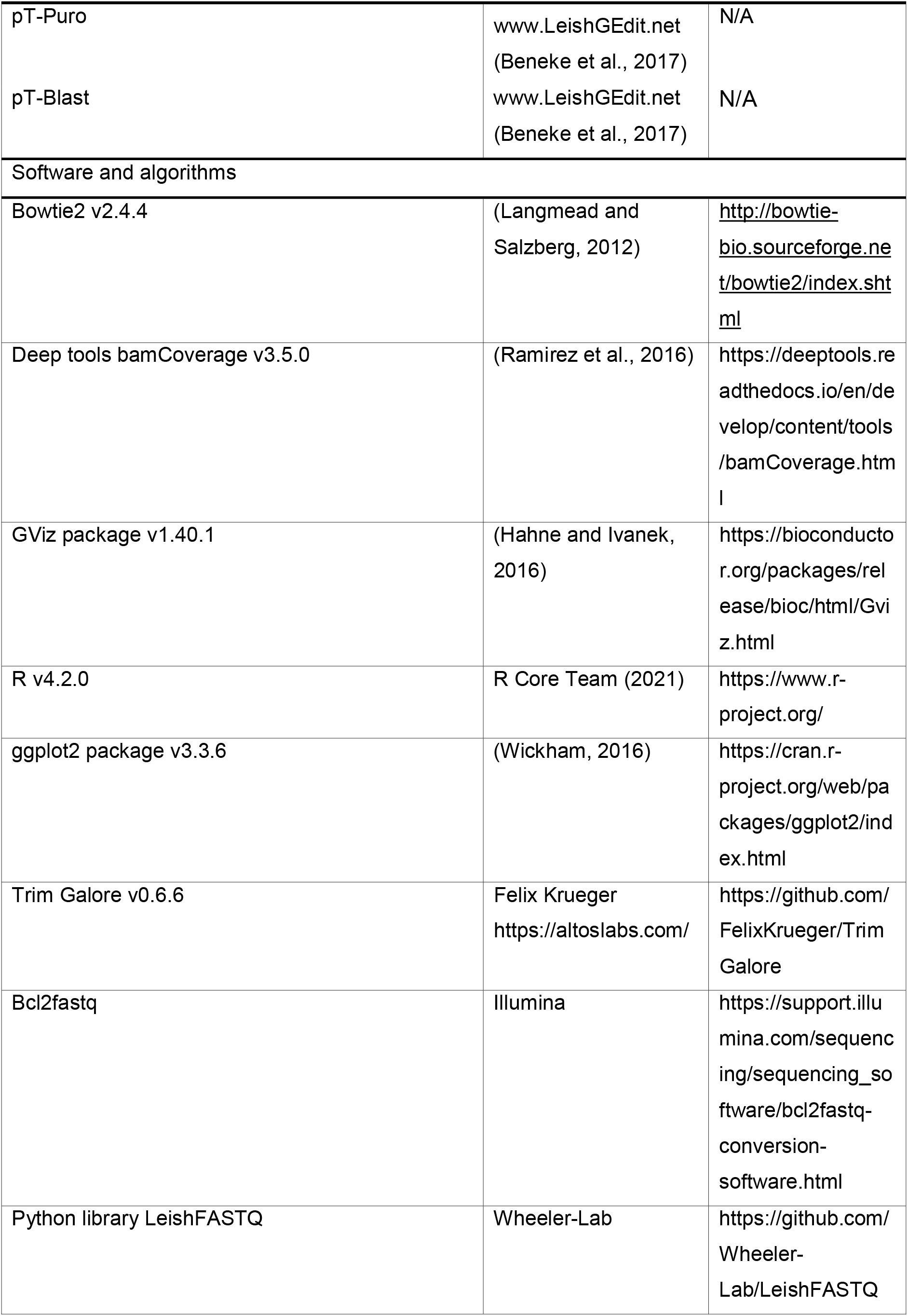

### Contact for Reagent and Resource Sharing

Further information can be directed to Eva Gluenz and requests for resources and reagents should be directed to and will be fulfilled by Eva Gluenz.

### Experimental Animals

Eight week old female BALB/c mice were purchased from Charles River Laboratories and maintained in the pathogen-free facility at the University of York. All mice used in experiments were socially housed under a 12h light/dark cycle and were only used in this experiment. All experiments were conducted according to the Animals (Scientific Procedures) Act of 1986, United Kingdom, and had approval from the University of York, Animal Welfare and Ethical Review Body (AWERB) committee.

### Pluripotent Stem Cells

Three human induced pluripotent stem cell (hiPSC) lines (SFC840-03-03, SFC841-03-01, and SFC856-03-04) used in this study were previously derived from in-house grown dermal fibroblasts from three healthy human donors reprogrammed with Cytotune Sendai viruses (ThermoFisher) (Ethics Committee: National Health Service, Health Research Authority, NRES Committee South Central, Berkshire, UK, REC 10/H0505/71). They have been published previously (Dafinca et al., 2016; Fernandes et al., 2016; Haenseler et al., 2017) and are deposited in EBiSC (STBCi026-A, STBCi044-A, STBCi063-A). They were maintained in the Stem Cell Facility at the James & Lillian Martin Centre for Stem Cell Research (Sir William Dunn School of Pathology, University of Oxford (United Kingdom), as QCed frozen banked stocks and cultured in mTeSR (StemCell Technologies) on Geltrex (Invitrogen) with minimal subsequent passaging (using 0.5mM EDTA (Invitrogen)) to ensure karyotypic integrity, as previously described (van Wilgenburg et al., 2013).

### Leishmania Parasites

Promastigote forms of the *L. mexicana* cell line *L. mex* Cas9 T7 (Beneke et al., 2017) were grown in T25 cm^2^ flasks at 28 °C or flat bottom well plates at 28 °C + 5 % CO_2_ in filter- sterilised M199 medium (Life Technologies) supplemented with 2.2 g/L NaHCO_3_, 0.005% hemin, 40 mM 4-(2-Hydroxyethyl)piperazine-1-ethanesulfonic acid (HEPES) pH 7.4 and 10 % FCS. 50 μg/ml Nourseothricin Sulphate and 32 μg/ml Hygromycin B were added to the medium for the maintenance of the *sp*Cas9 and T7 RNA polymerase transgenes (Beneke et al., 2017).

### Transportome protein identification

Membrane transporter proteins included in the TransLeish library were identified by searching the genome of *L. mexicana* MHOM/GT/2001/U1103 (TritrypDB (Shanmugasundram et al., 2023)) with the keywords “*transporter, transport, exchanger, permease, carrier, channel, porin, pump, symporter, antiporter, uniporter, porter, facilitator, efflux, ABC, ATP-binding cassette, P-glycoprotein, ATPase”*. Criteria for inclusion were the presence of protein domains indicative of transporter function (Supplementary Table 1), published experimental evidence of its function, as well as protein sequence identity to homologues present in TCD. From resulting candidate proteins, we removed components of transport systems involved in protein translocation, intraflagellar transport, nuclear pores, proteins of the electron transport chain and components of intraorganellar membrane contact sites. To assign proteins to a TCDB family, each sequence was used as a query for a TCDB BLAST search (https://www.tcdb.org/progs/blast.php) using the default parameters.

### Parasite growth curves

For assessment of cell growth under different pH conditions the pH of supplemented M199 medium was adjusted to either 7.4 or 5.5, using HCl 10N solution. Cells were seeded at a starting density of 1x10^6^ cells/ml and left to grow for up to 96h; each condition was assessed in three replicate cultures. Cell counts and cell volumes were determined with the CASY^®^ cell counter (Cambridge Bioscience) using a 60 µm capillary and measurement range set between 2 and 10 µm.

### Macrophage cell culture

Human induced pluripotent stem cell (iPSC) derived macrophages were generated as previously described (van Wilgenburg et al., 2013) (Vaughan-Jackson et al., 2021). Briefly, iPSC from three different healthy human donors were expanded for one week, lifted with tryplE (Invitrogen), formed into uniform 10,000-cell embryoid bodies using Aggrewells (StemCell Technologies) in mTeSR with 1 mM Rock-inhibitor (Y27632; Abcam) and differentiated towards hemogenic endothelium with growth factors 50 ng/mL BMP4 (Invitrogen), 50 ng/mL VEGF (Invitrogen), and 20 ng/mL SCF (Miltenyi) with daily medium change for one week. The embryoid bodies were then transferred to T175 flasks in XVIVO- 15 (Lonza) with 100 ng/mL M-CSF (Invitrogen) and 25 ng/mL IL-3 (Invitrogen) to induce myelopoeisis. Macrophage precursors were harvested from the supernatant, passed through a 40 µm cell strainer to exclude cell aggregates or embryoid bodies, centrifuged at 400 G, resuspended in macrophage medium and cultured in T25 flasks (Advanced DMEM/F-12 (Gibco), GlutaMAX 2mM (Gibco), HEPES 15mM (Gibco), human recombinant Insulin solution 5µg/mL (Sigma), M-CSF 100ng/mL (Invitrogen) Penicillin-Streptomycin 1% (Gibco)) for 7 days at 37 °C and 5 % CO_2_, with medium changes at day 3-4, to induce differentiation to ready-to-use adherent macrophages.

24h prior to infections, a total of 10^6^ macrophages were seeded in the wells of flat-bottomed 6-well plates, medium was switched to fresh pre-warmed DMEM + 10 % FBS + 1 % pen-strep and cells were further incubated at 34 °C, 5 % CO_2_. Pools of barcoded *Leishmania* cell lines were generated as described in the section “Pooling of cells for bar-seq experiments”. Stationary phase cultures of promastigote forms were used to infect the 34 °C-adapted macrophages with a multiplicity of infection of 20 parasites to 1 macrophage. After 3 h cells were washed three times with medium to remove free parasites. Fresh medium was added to the washed infected macrophages and cells were left to incubate at 34 °C, 5 % CO_2_ until DNA extraction.

### Animal work

To assess virulence and fitness of the null mutants in the pooled knock-out library, three groups of six female BALB/c mice were infected subcutaneously at the left footpad with 2x10^6^ promastigotes per animal. Presence or absence of lesion was observed, and footpads were measured weekly using a calliper. Animals were culled after 72h, 3- and 6-weeks post inoculum for collection of the footpad.

### CRISPR-Cas9 gene knockouts

Gene deletions were done using the CRISPR-Cas9 method described in (Beneke et al., 2017). In brief, the *L. mex* Cas9 T7 cell line was transfected with a mix of four PCR amplicons: Two serving as transcription templates for sgRNAs, targeting sites upstream (5’) and downstream (3’) of the target gene ORF, the other two amplicons serving as donor DNA for DNA break repair, containing a blasticidin and puromycin resistance gene, respectively, flanked by 30 nt of sequence identical to the target locus and a unique 17-nt barcode together with the barcode flanking sequences GTGTATCGGATGTCAGTTGC and GTATAATGCAGACCTGCTGC (Beneke and Gluenz, 2020). The primer sequences for generating the PCR amplicons were taken from (McCoy et al., 2023)Beneke, 2022} and pTBlast and pTPuro served as template DNA for the amplification of donor DNA. For array deletions, an identical strategy was used, by replacing the gene array, with donor DNA cassettes that were amplified using the upstream forward primer for the first gene in the array and the downstream reverse primer for the last gene in the array (all primer sequences available in Supplementary Table 5). Parasite transfections were done on 96-well plates as described in (Beneke and Gluenz, 2019). Puromycin and blasticidin were added between 6 and 12h after transfection and drug resistant populations selected as described in (Beneke and Gluenz, 2019), with at least four passages before extraction of genomic DNA. Fitness screens were done using populations for which gene deletions were confirmed by diagnostic PCR, without further subcloning.

### Diagnostic PCR for knockout validation

Genomic DNA was extracted using the protocol from (Rotureau et al., 2005). A diagnostic PCR was done to test for the loss of the target gene sequence in putative knockout cell lines, with L. mex Cas9 T7 gDNA serving as a control. Primer sequences were taken from (Beneke et al., 2019) (sequences available in Supplementary Table 5). A second PCR was performed on the mutant gDNA, amplifying a fragment of the blasticidin resistance gene with primers 518F: 5’-CACCCTCATTGAAAGAGCAAC-3’ and 518R: 5’-CACTATCGCTTTGATCCCAGGA-3’. Finally, to test for presence of gDNA in samples extracted from mutants independently from the introduction of the drug resistance cassette, primers targeting genes *LmxM.34.5260* (*FAP174*), *LmxM.18.0610* or *LmxM.18.1620*, respectively, were used for PCR amplification.

FAF: 5’-AGAGGAACAGAAACCCGAGGAC-3’, FAR: 5’-CTTTAGCGATTCCACCTCTGCG-3’ 94F: 5’-ACCCTTTCACGAGCATGATGGA-3’, 94R: 5’-GAGCTCAATAAAGTCCGCACCG-3’ 96F: 5’-AATGACTCGACCAAGATCGGGG-3’, 96R: 5’-CCCAGAAAGAAGCACGCAGATG-3’

### Pooling of cells for bar-seq experiments

The barcoded mutant and parental cell lines were combined such that the mixed pool contained similar numbers of each individual cell line: For the *in vitro* screen, a total of 252 transporter mutants, five barcoded parental lines (SBL1-5; barcodes introduced into the SSU locus (Beneke et al., 2019)), and seven non-transporter knock-out mutants, of which three acted as controls; Δ*IFT88* (*LmxM.27.1130*), Δ*dihydroorotase* (*LmxM.16.0580*) and Δ*MBO2* (*LmxM.33.2480*), were combined into a pool of 1x10^6^ cells/ml. This pool was split into three equal aliquots, which were left to grow in separate flasks for 48 hours at 28 °C + 5 % CO_2_ in M199 medium. For iPSC-derived macrophage and mouse infections, a total of 254 transporter mutants (184 confirmed null mutants and 70 incomplete knockouts, see Supplementary Table 1), 57 knockout mutants of flagellar proteins, five barcoded parental lines (SBL1-5) and five control knock-out mutants Δ*Kharon1* (*LmxM.36.5850*), Δ*BBS2* (*LmxM.29.0590*), Δ*IFT88* (LmxM.27.1130), Δ*PMM* (*LmxM.36.1960*) and Δ*GDP-MP* (*LmxM.23.0110*) were pooled in similar proportions. Fifteen additional barcoded parental lines (SBL6-20) were added to the pool at dilutions of 1:2 (SBL 6-8), 1:4 (SBL 9-11), 1:8 (SBL 12-14), 1:16 (SBL 15-17) and 1:32 (SBL 18-20), relative to the other mutants. To account for the slower growth rates of some mutants, the *Leishmania* were first combined in three sub-pools, seeded at different initial cell densities in M199 medium: 325 cell lines with “normal” growth, 1x10^6^ cells/ml; six “slow” growing cell lines, 3x10^6^ cells/ml; four “very slow” cell lines, 6x10^6^ cells/ml. The three sub-pools were then left to grow for four days to stationary phase before combining the cells into a masterpool, from which replicates were prepared for DNA isolation (T0) and infection of macrophages or mice in 6 replicates.

### Sampling of DNA for sequencing

*Leishmania* promastigote genomic DNA was extracted from approximately 1x10^7^ cells using the Qiagen DNeasy Blood & Tissue Kit according to the manufacturer’s instructions, eluting in 40 μl of bi-distilled water (Ambion). For extraction of genomic DNA from infected macrophages, cells were scraped from the bottom of the wells, pelleted by centrifugation and total DNA was extracted, using the Qiagen DNeasy Blood & Tissue Kit as described above. DNA extraction from the mouse footpad lesions was performed using the Qiagen DNeasy Blood & Tissue Kit according to the manufacturer’s instructions, except for an extended incubation in proteinase K solution and a final elution in 75 μl of TE buffer.

### Bar-seq library preparation and sequencing

For each DNA sample, the barcode region was amplified with custom designed p5 and p7 primers containing indexes for multiplexing and adapters for Illumina sequencing (The primers were synthesized at 0.2 µmol scale and PAGE purified by Life Technologies or by Microsynth.ch) (Beneke et al., 2019).

For promastigote DNA samples, the quantity, purity and length of the total genomic DNA was assessed using a Thermo Fisher Scientific Qubit 4.0 fluorometer with the Qubit dsDNA HS Assay Kit (Thermo Fisher Scientific, Q32854), a DeNovix DS-11 FX spectrophotometer and an Agilent FEMTO Pulse System with a Genomic DNA 165 kb Kit (Agilent, FP-1002-0275), respectively. Bar-seq amplicon libraries were made via PCR as follows: 100 ng *Leishmania* gDNA, 1 µL of 10 mM dNTP Mix (R1091, Thermo Fisher Scientific), 1.25 µL of 4 µM P5 primer, 1.25 uL of 4 µM P7 primer, 1 uL of Platinum SuperFi II DNA Polymerase (12361010, Thermo Fisher Scientific), 10 uL of the 5X SuperFi II Buffer and nuclease free water in a total volume of 50 µL. A negative template control (NTC) was included at each PCR set-up. The following three-step protocol, was employed: an initial denaturation at 94 °C for 5 min, then 31 cycles of denaturation at 94 °C for 30 s, annealing at 60 °C for 30 s and extension at 72 °C 15 min. A final extension at 72 °C for 7 min was employed before a 4 °C hold. The CleanNGS kit (CNGS-0050, Clean NA) based on paramagnetic particle technology was used according to the user manual to purify each amplicon library. Thereafter, each library was evaluated using a Thermo Fisher Scientific Qubit 4.0 fluorometer with the Qubit dsDNA HS Assay Kit (Thermo Fisher Scientific, Q32854) and an Agilent Fragment Analyzer (Agilent) with a HS NGS Fragment Kit (Agilent, DNF-474), respectively. The bar-seq library shows a clear peak at 218 bp. Libraries were equimolar pooled into one pool. These library pools were further qPCR-based quantified using the JetSeq Library Quantification Lo-ROX kit (BIO-68029, Bioline) according to the manufacturer’s guidelines. The amplicon library pools were diluted to 4 nM and spiked with 30 % PhiX Control v3 (illumina, FC-110-3001) and loaded at an on-plate concentration of 8 pM to reach a cluster density of ∼1’000 K/mm2. The libraries were-paired end sequenced with a read set-up of 76:6:8:76 using a MiSeq Reagent Kit v3 150 cycles (illumina, MS-102-3001) on an illumina MiSeq instrument. The quality of each sequencing run was assessed using illumina Sequencing Analysis Viewer (illumina version 2.4.7) and all base call files were demultiplexed and converted into FASTQ files using illumina bcl2fastq conversion software v2.20 but the default setting were changed to allow for 0 mismatches. All steps post gDNA extraction to sequencing data generation and data utility were performed at the Next Generation Sequencing Platform, University of Bern, Switzerland.

For samples from hiPSC-Mac, barcodes were amplified from 100 ng DNA using 31 PCR cycles. For mouse footpad tissue samples, 2000 ng DNA was used as input, with 35 PCR cycles. Amplicons were purified using SPRI magnetic beads and multiplexed by pooling them in equal proportions. The pooled library was once again bead purified and quantified by qPCR using NEBNext Library Quant Kit (NEB, E7630). The library size was verified using a High Sensitivity DNA Kit on a 2100AB Bioanalyzer instrument (Agilent). For whole genome sequencing, a total of 600 ng of genomic DNA from the relevant cell lines was fragmented using DNA fragmentase (NEB, M0348) for 25 minutes at 37 °C. The reaction was stopped using 5 µl of 0.5 M Ethylenediaminetetraacetic Acid (EDTA, Sigma). Fragments of about 350 bp were purified and single-indexed DNA libraries were prepared using standard Illumina TruSeq Nano DNA Library Prep (Reference Guide 15041110 D) and Indexed Sequencing (Overview Guide 15057455). Library fragment size and concentrations were quality verified by qPCR using the NEBNext Library Quant Kit (NEB, E7630) and loading 1:1, 1:3, 1:10 and 1:50 dilutions of the library on a Bioanalyzer High sensitivity DNA Chip (Agilent) and running it on Bioanalyzer 2100AB (Agilent). The amplicon pool was diluted to 4 nM and mixed with 20-40 % single-indexed *Leishmania* genomic DNA. The final library was spiked with 1 % PhiX DNA and the Illumina sequencer was loaded with 8 pM to allow low cluster density (∼800 K/mm^2^). Sequencing was performed at the Glasgow Polyomics facility using NextSeq Mid Output v2 150 bp kits, with paired-end sequencing (2x75 bp), following the manufactures instructions. NextSeq raw files were de-multiplexed using bcl2fastq (Illumina).

Sequencing reads containing barcode sequences were grouped and counted by scanning sequencing reads for the correct left (GTGTATCGGATGTCAGTTGC) and right (GTATAATGCAGACCTGCTGC) flanking sequences (allowing up to two mismatches per flank) and extracting the 17nt barcode using the python library LeishFASTQ (https://github.com/Wheeler-Lab/LeishFASTQ). Previously designed barcode sequences for all genes have been reported in (Beneke et al., 2022), therefore reads were grouped into these four categories:

i. Invalid reads: sequencing reads that do not contain a valid barcode-like sequence (incorrect or non-existent flanking sequence while allowing for two mismatched nts, barcode sequence of the wrong length).
ii. Unknown barcode sequences: sequencing reads that contain a valid barcode-like construct, but the barcode sequence does not exist in the LeishGEdit database.
iii. Foreign barcode sequences: sequencing reads that contain a valid barcode sequence that exists in the LeishGEdit database (allowing a single nt mismatch in the barcode sequence) but the corresponding cell line had not been included in the pool of cell lines.
iv. Pool member barcode sequences: sequencing reads that contain a valid barcode sequence of a cell line that had been included in the pool (allowing a single nt mismatch in the barcode sequence).

The sequence read counts for each of these categories are shown in Supplementary Table 4 for all sequencing samples.

### Quantification of barcoded cell line fitness

Barcode proportions were calculated by dividing the detected abundance of a given barcode in the sample by the total number of barcoded reads generated from that sample to control for variation in sequencing sample read depths. The barcode change over time was assessed by taking the barcode proportion at a given time point and dividing by the barcode proportion at the 0 h time point for each replicate separately to control for variation in barcoded cell line prevalence in the population before the start of the assay. Cell line fitness scores were assigned by dividing the median of the corresponding barcode change over all replicates with the median change of all parental cell line barcodes in all replicates. A fitness score above one indicates that proportion of barcodes from a particular cell line have increased relative to the parental cell line reference, corresponding to faster growth and/or better survival than the parental cell line from the start of the assay up to that time point. A fitness score below one indicates the inverse. P-values were calculated using a Mann-Whitney U test against the null hypothesis that the barcode changes from all replicates of a particular cell line in a given time point cannot be distinguished from the barcode changes of all parental cell lines in all replicates of the same time point. Cell lines were labelled as having a strong fitness phenotype in a given time point if their p-value was below 0.05 and their fitness score was either below 0.5 (deleterious phenotype) or above 2 (beneficial phenotype).

### Genomic Data Processing

Genomic sequencing data was trimmed using Trim Galore version 0.6.6 to remove adaptor sequence and bases where the Phred-scored quality was below 20 from the 3’ end of each read. Reads that were less than 20bp after trimming were discarded. Trimmed reads were aligned to the *Leishmania mexicana* MHOMGT2001U1103 reference genome obtained from build 55 of TriTrypDB. Alignment was carried out using bowtie2 version 2.4.4 using default parameters.

### Confirming Knockouts from Genomic Data

Quantitative coverage data was extracted from the alignments in bigwig format using Deeptools bamCoverage version 3.5.0, using a 50 bp bin size and normalised using the Counts Per Million (CPM) method, where the number of counts per bin is divided by the total number of mapped reads in millions. For each knockout, normalised data from the relevant line and from the Cas9 T7 parental line was plotted over the genomic region of the gene that was expected to be knocked out +/- 1.5 kb using the GViz package version 1.40.1 in R version 4.2.0.

### Estimating Ploidy

Quantitative coverage data was extracted from the alignments in bedgraph format using Deeptools bamCoverage using a 1kb bin size. Assuming that the majority of chromosomes are diploid, scaling to the ploidy was carried out by dividing the coverage for each bin by the median coverage over all bins divided by 2 (coverage/(medianCoverage/2)). The distribution of scaled coverage values was plotted for each chromosome in each line with the Cas9 T7 parental line for comparison, using ggplot2 version 3.3.6 in R. Additionally, an estimate of the ploidy of each chromosome in each line was obtained by dividing the median coverage over all the bins in the chromosome by the median coverage over all the bins in the genome divided by 2 (medianCoverageChromosome/(medianCoverageGenome/2)).

## Author contributions

Conceptualisation, AAW, CMC, EG Methodology, TB, UD, RJW, EG Software, UD, RJW

Formal analysis, AAW, UD, EG Investigation, AAW, CMC, RN, TB, KC Resources, EG, JCM, SAC

Data curation, AAW, EG, UD, RJW Writing original draft, AAW, EG

Writing review and editing, AAW, RJW, TB, RN, JCM, EG Visualisation, AAW

Supervision, EG

Project administration, EG

Funding Acquisition, EG, AAW, RJW, JCM

## Supporting information

Legends and References to Supplementary Files.pdf

Supplementary Table 1 - TransLeishDB

Supplementary Table_2 - Diagnostic PCR Results

Supplementary Table 3 - Tandem Arrays

Supplementary Table 4 - Fitness Scores and Read Counts

Supplementary Table 5 - Barcodes and Primers

## Acknowledgements

We like to thank Amanda Williams (University of Oxford), Julie Galbraith and Csilla Balazs (University of Glasgow, Polyomics facility), Pamela Nicholson and Daniela Steiner (NGS Facility, University of Bern) for help with Illumina sequencing, William James (James & Lillian Martin Centre, Sir William Dunn School of Pathology, University of Oxford) for supporting the work with iPSC-derived macrophages, Caroline Ricce Espada for generating the Ros3 knockout cell line, Keith Gull (Sir William Dunn School of Pathology, University of Oxford) for access to equipment and all past and current members of EG lab for helpful discussions.

## References

1. Alsford, S., Turner, D.J., Obado, S.O., Sanchez-Flores, A., Glover, L., Berriman, M., Hertz-Fowler, C., and Horn, D. (2011). High-throughput phenotyping using parallel sequencing of RNA interference targets in the African trypanosome. Genome research 21, 915–924.

2. Alzahrani, K.J.H., Ali, J.A.M., Eze, A.A., Looi, W.L., Tagoe, D.N.A., Creek, D.J., Barrett, M.P., and de Koning, H.P. (2017). Functional and genetic evidence that nucleoside transport is highly conserved in *Leishmania* species: Implications for pyrimidine-based chemotherapy. International journal for parasitology Drugs and drug resistance 7, 206–226.

3. Annang, F., Perez-Moreno, G., Garcia-Hernandez, R., Cordon-Obras, C., Martin, J., Tormo, J.R., Rodriguez, L., de Pedro, N., Gomez-Perez, V., Valente, M., et al. (2015). High-throughput screening platform for natural product-based drug discovery against 3 neglected tropical diseases: human African trypanosomiasis, leishmaniasis, and Chagas disease. J Biomol Screen 20, 82–91.

4. Antoine, J.C., Prina, E., Jouanne, C., and Bongrand, P. (1990). Parasitophorous vacuoles of Leishmania amazonensis-infected macrophages maintain an acidic pH. Infect Immun 58, 779–787.

5. Arcari, T., Manzano, J.I., and Gamarro, F. (2017). ABCI3 Is a New Mitochondrial ABC Transporter from Leishmania major Involved in Susceptibility to Antimonials and Infectivity. Antimicrobial agents and chemotherapy 61.

6. Baker, N., Hamilton, G., Wilkes, J.M., Hutchinson, S., Barrett, M.P., and Horn, D. (2015). Vacuolar ATPase depletion affects mitochondrial ATPase function, kinetoplast dependency, and drug sensitivity in trypanosomes. Proceedings of the National Academy of Sciences of the United States of America 112, 9112–9117.

7. Bakker-Grunwald, T. (1992). Ion transport in parasitic protozoa. J Exp Biol 172, 311–322.

8. Bates, P.A., Robertson, C.D., Tetley, L., and Coombs, G.H. (1992). Axenic cultivation and characterization of Leishmania mexicana amastigote-like forms. Parasitology 105 ( *Pt 2*), 193–202.

9. Beneke, T., Demay, F., Hookway, E., Ashman, N., Jeffery, H., Smith, J., Valli, J., Becvar, T., Myskova, J., Lestinova, T., et al. (2019). Genetic dissection of a *Leishmania* flagellar proteome demonstrates requirement for directional motility in sand fly infections. PLoS pathogens 15, e1007828.

10. Beneke, T., Dobramysl, U., Catta-Preta, C.M.C., Mottram, J.C., Gluenz, E., and Wheeler, R.J. (2022). Genome sequence of Leishmania mexicana MNYC/BZ/62/M379 expressing Cas9 and T7 RNA polymerase. Wellcome Open Res 7, 294.

11. Beneke, T., and Gluenz, E. (2019). LeishGEdit: A Method for Rapid Gene Knockout and Tagging Using CRISPR-Cas9. Methods Mol Biol 1971, 189–210.

12. Beneke, T., and Gluenz, E. (2020). Bar-seq strategies for the LeishGEdit toolbox. Molecular and biochemical parasitology 239, 111295.

13. Beneke, T., Madden, R., Makin, L., Valli, J., Sunter, J., and Gluenz, E. (2017). A CRISPR Cas9 high-throughput genome editing toolkit for kinetoplastids. Royal Society open science 4, 170095.

14. Bowman, E.J., and Bowman, B.J. (2005). V-ATPases as drug targets. J Bioenerg Biomembr 37, 431–435.

15. Burchmore, R.J., and Barrett, M.P. (2001). Life in vacuoles--nutrient acquisition by Leishmania amastigotes. International journal for parasitology 31, 1311–1320.

16. Burchmore, R.J.S., Rodriguez-Contreras, D., McBride, K., Barrett, M.P., Modi, G., Sacks, D., and Landfear, S.M. (2003). Genetic characterization of glucose transporter function in Leishmania mexicana. Proceedings of the National Academy of Sciences of the United States of America 100, 3901–3906.

17. Cabello-Donayre, M., Orrego, L.M., Herraez, E., Vargas, P., Martinez-Garcia, M., Campos-Salinas, J., Perez-Victoria, I., Vicente, B., Marin, J.J.G., and Perez- Victoria, J.M. (2020). Leishmania heme uptake involves LmFLVCRb, a novel porphyrin transporter essential for the parasite. Cell Mol Life Sci 77, 1827–1845.

18. Campos-Salinas, J., Cabello-Donayre, M., Garcia-Hernandez, R., Perez-Victoria, I., Castanys, S., Gamarro, F., and Perez-Victoria, J.M. (2011). A new ATP-binding cassette protein is involved in intracellular haem trafficking in Leishmania. Molecular microbiology 79, 1430–1444.

19. Collins, M.P., and Forgac, M. (2020). Regulation and function of V-ATPases in physiology and disease. Biochim Biophys Acta Biomembr 1862, 183341.

20. Dafinca, R., Scaber, J., Ababneh, N., Lalic, T., Weir, G., Christian, H., Vowles, J., Douglas, A.G., Fletcher-Jones, A., Browne, C., et al. (2016). C9orf72 Hexanucleotide Expansions Are Associated with Altered Endoplasmic Reticulum Calcium Homeostasis and Stress Granule Formation in Induced Pluripotent Stem Cell-Derived Neurons from Patients with Amyotrophic Lateral Sclerosis and Frontotemporal Dementia. Stem Cells 34, 2063–2078.

21. De Muylder, G., Vanhollebeke, B., Caljon, G., Wolfe, A.R., McKerrow, J., and Dujardin, J.C. (2016). Naloxonazine, an Amastigote-Specific Compound, Affects Leishmania Parasites through Modulation of Host-Encoded Functions. PLoS Negl Trop Dis 10, e0005234.

22. Eaton, A.F., Merkulova, M., and Brown, D. (2021). The H(+)-ATPase (V-ATPase): from proton pump to signaling complex in health and disease. Am J Physiol Cell Physiol 320, C392–C414.

23. Elbourne, L.D., Tetu, S.G., Hassan, K.A., and Paulsen, I.T. (2017). TransportDB 2.0: a database for exploring membrane transporters in sequenced genomes from all domains of life. Nucleic acids research 45, D320–D324.

24. Fernandes, H.J., Hartfield, E.M., Christian, H.C., Emmanoulidou, E., Zheng, Y., Booth, H., Bogetofte, H., Lang, C., Ryan, B.J., Sardi, S.P., et al. (2016). ER Stress and Autophagic Perturbations Lead to Elevated Extracellular alpha-Synuclein in GBA-N370S Parkinson’s iPSC-Derived Dopamine Neurons. Stem Cell Reports 6, 342–356.

25. Garami, A., and Ilg, T. (2001). The role of phosphomannose isomerase in Leishmania mexicana glycoconjugate synthesis and virulence. The Journal of biological chemistry 276, 6566–6575.

26. Glaser, T.A., Baatz, J.E., Kreishman, G.P., and Mukkada, A.J. (1988). pH homeostasis in Leishmania donovani amastigotes and promastigotes. Proceedings of the National Academy of Sciences of the United States of America 85, 7602–7606.

27. Goldman-Pinkovich, A., Balno, C., Strasser, R., Zeituni-Molad, M., Bendelak, K., Rentsch, D., Ephros, M., Wiese, M., Jardim, A., Myler, P.J., et al. (2016). An Arginine Deprivation Response Pathway Is Induced in Leishmania during Macrophage Invasion. PLoS pathogens 12, e1005494.

28. Haenseler, W., Zambon, F., Lee, H., Vowles, J., Rinaldi, F., Duggal, G., Houlden, H., Gwinn, K., Wray, S., Luk, K.C., et al. (2017). Excess alpha-synuclein compromises phagocytosis in iPSC-derived macrophages. Scientific reports 7, 9003.

29. Hahne, F., and Ivanek, R. (2016). Visualizing Genomic Data Using Gviz and Bioconductor. Methods Mol Biol 1418, 335–351.

30. Huynh, C., Sacks, D.L., and Andrews, N.W. (2006). A Leishmania amazonensis ZIP family iron transporter is essential for parasite replication within macrophage phagolysosomes. The Journal of experimental medicine 203, 2363–2375.

31. Huynh, C., Yuan, X., Miguel, D.C., Renberg, R.L., Protchenko, O., Philpott, C.C., Hamza, I., and Andrews, N.W. (2012). Heme uptake by Leishmania amazonensis is mediated by the transmembrane protein LHR1. PLoS pathogens 8, e1002795.

32. Inbar, E., Schlisselberg, D., Suter Grotemeyer, M., Rentsch, D., and Zilberstein, D. (2013). A versatile proline/alanine transporter in the unicellular pathogen Leishmania donovani regulates amino acid homoeostasis and osmotic stress responses. The Biochemical journal 449, 555–566.

33. Johnston, N.R., Nallur, S., Gordon, P.B., Smith, K.D., and Strobel, S.A. (2020). Genome-Wide Identification of Genes Involved in General Acid Stress and Fluoride Toxicity in Saccharomyces cerevisiae. Front Microbiol 11, 1410.

34. Jones, N.G., Catta-Preta, C.M.C., Lima, A., and Mottram, J.C. (2018). Genetically Validated Drug Targets in Leishmania: Current Knowledge and Future Prospects. ACS infectious diseases 4, 467–477.

35. Kloehn, J., Saunders, E.C., O’Callaghan, S., Dagley, M.J., and McConville, M.J. (2015). Characterization of metabolically quiescent Leishmania parasites in murine lesions using heavy water labeling. PLoS pathogens 11, e1004683.

36. Koreny, L., Obornik, M., and Lukes, J. (2013). Make it, take it, or leave it: heme metabolism of parasites. PLoS pathogens 9, e1003088.

37. Langmead, B., and Salzberg, S.L. (2012). Fast gapped-read alignment with Bowtie 2. Nature methods 9, 357–359.

38. Ma, D., Russell, D.G., Beverley, S.M., and Turco, S.J. (1997). Golgi GDP-mannose uptake requires Leishmania LPG2. A member of a eukaryotic family of putative nucleotide-sugar transporters. The Journal of biological chemistry 272, 3799–3805.

39. Manzano, J.I., Perea, A., Leon-Guerrero, D., Campos-Salinas, J., Piacenza, L., Castanys, S., and Gamarro, F. (2017). Leishmania LABCG1 and LABCG2 transporters are involved in virulence and oxidative stress: functional linkage with autophagy. Parasites & vectors 10, 267.

40. Marchesini, N., and Docampo, R. (2002). A plasma membrane P-type H(+)-ATPase regulates intracellular pH in Leishmania mexicana amazonensis. Molecular and biochemical parasitology 119, 225–236.

41. Marquis, N., Gourbal, B., Rosen, B.P., Mukhopadhyay, R., and Ouellette, M. (2005). Modulation in aquaglyceroporin AQP1 gene transcript levels in drug-resistant Leishmania. Molecular microbiology 57, 1690–1699.

42. Martin, R.E. (2020). The transportome of the malaria parasite. Biol Rev Camb Philos Soc 95, 305–332.

43. Martinez-Garcia, M., Campos-Salinas, J., Cabello-Donayre, M., Pineda-Molina, E., Galvez, F.J., Orrego, L.M., Sanchez-Canete, M.P., Malagarie-Cazenave, S., Koeller, D.M., and Perez-Victoria, J.M. (2016). LmABCB3, an atypical mitochondrial ABC transporter essential for Leishmania major virulence, acts in heme and cytosolic iron/sulfur clusters biogenesis. Parasites & vectors 9, 7.

44. McConville, M.J., Saunders, E.C., Kloehn, J., and Dagley, M.J. (2015). Leishmania carbon metabolism in the macrophage phagolysosome- feast or famine? F1000Res 4, 938.

45. McCoy, C.J., Paupelin-Vaucelle, H., Gorilak, P., Beneke, T., Varga, V., and Gluenz, E. (2023). ULK4 and Fused/STK36 interact to mediate assembly of a motile flagellum. Molecular biology of the cell, mbcE22060222.

46. Meade, J.C., Hudson, K.M., Stringer, S.L., and Stringer, J.R. (1989). A tandem pair of Leishmania donovani cation transporting ATPase genes encode isoforms that are differentially expressed. Molecular and biochemical parasitology 33, 81–91.

47. Mira, N.P., Palma, M., Guerreiro, J.F., and Sa-Correia, I. (2010). Genome-wide identification of Saccharomyces cerevisiae genes required for tolerance to acetic acid. Microb Cell Fact 9, 79.

48. Mittra, B., Laranjeira-Silva, M.F., Perrone Bezerra de Menezes, J., Jensen, J., Michailowsky, V., and Andrews, N.W. (2016). A Trypanosomatid Iron Transporter that Regulates Mitochondrial Function Is Required for Leishmania amazonensis Virulence. PLoS pathogens 12, e1005340.

49. Nayak, A., Akpunarlieva, S., Barrett, M., and Burchmore, R. (2018). A defined medium for Leishmania culture allows definition of essential amino acids. Experimental parasitology 185, 39–52.

50. Perez-Victoria, F.J., Gamarro, F., Ouellette, M., and Castanys, S. (2003). Functional cloning of the miltefosine transporter. A novel P-type phospholipid translocase from Leishmania involved in drug resistance. The Journal of biological chemistry 278, 49965–49971.

51. Perzov, N., Padler-Karavani, V., Nelson, H., and Nelson, N. (2002). Characterization of yeast V-ATPase mutants lacking Vph1p or Stv1p and the effect on endocytosis. J Exp Biol 205, 1209–1219.

52. Ramirez, F., Ryan, D.P., Gruning, B., Bhardwaj, V., Kilpert, F., Richter, A.S., Heyne, S., Dundar, F., and Manke, T. (2016). deepTools2: a next generation web server for deep-sequencing data analysis. Nucleic acids research 44, W160–165.

53. Rodriguez-Contreras, D., Feng, X., Keeney, K.M., Bouwer, H.G., and Landfear, S.M. (2007). Phenotypic characterization of a glucose transporter null mutant in Leishmania mexicana. Molecular and biochemical parasitology 153, 9–18.

54. Rotureau, B., Gego, A., and Carme, B. (2005). Trypanosomatid protozoa: a simplified DNA isolation procedure. Experimental parasitology 111, 207–209.

55. Saier, M.H., Jr. (2000). A functional-phylogenetic classification system for transmembrane solute transporters. Microbiol Mol Biol Rev 64, 354–411.

56. Saier, M.H., Jr., Reddy, V.S., Tamang, D.G., and Vastermark, A. (2014). The transporter classification database. Nucleic acids research 42, D251–258.

57. Saier, M.H., Reddy, V.S., Moreno-Hagelsieb, G., Hendargo, K.J., Zhang, Y., Iddamsetty, V., Lam, K.J.K., Tian, N., Russum, S., Wang, J., et al. (2021). The Transporter Classification Database (TCDB): 2021 update. Nucleic acids research 49, D461–D467.

58. Saunders, E.C., Ng, W.W., Kloehn, J., Chambers, J.M., Ng, M., and McConville, M.J. (2014). Induction of a stringent metabolic response in intracellular stages of Leishmania mexicana leads to increased dependence on mitochondrial metabolism. PLoS pathogens 10, e1003888.

59. Shanmugasundram, A., Starns, D., Bohme, U., Amos, B., Wilkinson, P.A., Harb, O.S., Warrenfeltz, S., Kissinger, J.C., McDowell, M.A., Roos, D.S., et al. (2023). TriTrypDB: An integrated functional genomics resource for kinetoplastida. PLoS Negl Trop Dis 17, e0011058.

60. Sidik, S.M., Huet, D., Ganesan, S.M., Huynh, M.H., Wang, T., Nasamu, A.S., Thiru, P., Saeij, J.P., Carruthers, V.B., Niles, J.C., et al. (2016). A Genome-wide CRISPR Screen in Toxoplasma Identifies Essential Apicomplexan Genes. Cell 166, 1423–1435 e1412.

61. Stasic, A.J., Chasen, N.M., Dykes, E.J., Vella, S.A., Asady, B., Starai, V.J., and Moreno, S.N.J. (2019). The Toxoplasma Vacuolar H(+)-ATPase Regulates Intracellular pH and Impacts the Maturation of Essential Secretory Proteins. Cell Rep 27, 2132–2146 e2137.

62. Tran, K.D., Rodriguez-Contreras, D., Vieira, D.P., Yates, P.A., David, L., Beatty, W., Elferich, J., and Landfear, S.M. (2013). KHARON1 mediates flagellar targeting of a glucose transporter in Leishmania mexicana and is critical for viability of infectious intracellular amastigotes. The Journal of biological chemistry 288, 22721–22733.

63. UniProt, C. (2021). UniProt: the universal protein knowledgebase in 2021. Nucleic acids research 49, D480–D489.

64. van Wilgenburg, B., Browne, C., Vowles, J., and Cowley, S.A. (2013). Efficient, long term production of monocyte-derived macrophages from human pluripotent stem cells under partly-defined and fully-defined conditions. PloS one 8, e71098.

65. Vaughan-Jackson, A., Stodolak, S., Ebrahimi, K.H., Browne, C., Reardon, P.K., Pires, E., Gilbert-Jaramillo, J., Cowley, S.A., and James, W.S. (2021). Differentiation of human induced pluripotent stem cells to authentic macrophages using a defined, serum-free, open-source medium. Stem Cell Reports 16, 1735–1748.

66. Vickers, T.J., and Beverley, S.M. (2011). Folate metabolic pathways in Leishmania. Essays in biochemistry 51, 63–80.

67. Vince, J.E., Tull, D., Landfear, S., and McConville, M.J. (2011). Lysosomal degradation of Leishmania hexose and inositol transporters is regulated in a stage-, nutrient- and ubiquitin-dependent manner. International journal for parasitology 41, 791–800.

68. Wickham, H. (2016). Ggplot2: Elegant graphics for data analysis (2nd ed.) (Springer Cham).

69. Williams, R.A., Tetley, L., Mottram, J.C., and Coombs, G.H. (2006). Cysteine peptidases CPA and CPB are vital for autophagy and differentiation in Leishmania mexicana. Molecular microbiology 61, 655–674.

70. Yernaux, C., Fransen, M., Brees, C., Lorenzen, S., and Michels, P.A. (2006). Trypanosoma brucei glycosomal ABC transporters: identification and membrane targeting. Mol Membr Biol 23, 157–172.

71. Zhu, Y., Davis, A., Smith, B.J., Curtis, J., and Handman, E. (2009). Leishmania major CorA-like magnesium transporters play a critical role in parasite development and virulence. International journal for parasitology 39, 713–723.

72. Zilberstein, D., Philosoph, H., and Gepstein, A. (1989). Maintenance of cytoplasmic pH and proton motive force in promastigotes of Leishmania donovani. Molecular and biochemical parasitology 36, 109–117.

