## Supplementary material for "TransLeish: Identification of membrane transporters essential for survival of intracellular *Leishmania* parasites in a systematic gene deletion screen": Legends and References to Supplementary Files.pdf

### Supplementary Figure legends

***Supplementary Figure 1. Diagnostic PCR analysis to determine gene deletion status.***
(A) Diagnostic PCR amplification of target gene for knockout validation. The open reading frame of the gene of interest (GOI) was amplified from gDNA with a forward (OF) and reverse (OR) primer pair and products assessed on an agarose gel. Two samples were assessed for each GOI: amplification from L. mex Cas9 T7 parental gDNA and amplification from gDNA of mutants that survived transfection and drug selection. Numbers above gel lanes refer to the mutant numbers, as listed in Supplementary Table 2. (B) Control PCR amplification of a non-target gene from mutant gDNA. From each mutant gDNA sample, a fragment of the Blasticidin resistance cassette was amplified with primers BF1 and BR1 (lanes marked with a purple star symbol) and of a second gene as depicted in the schematic above the gel image. Gel lanes are numbered as in A. (C) Repeats of diagnostic PCR for selected mutants as in A and B, but using a higher annealing temperature of 60ºC. (D) Repeats of diagnostic PCR for selected mutants where results from the first PCR were unclear, using the same conditions as in (A) and (B). See Supplementary Table 1 for a summary of diagnostic PCR results.

***Supplementary Figure 2. Assessment of gene deletion by whole genome sequencing of selected mutants.***
Whole genome sequencing was carried out for puromycin and blasticidin resistant mutant lines Δ*LmxM.03.500*, Δ*LmxM.08_29.2780*, Δ*LmxM.08_29.1640*, Δ*LmxM.22.1420*, Δ*LmxM.32.3040*, Δ*LmxM.31.2660* and Δ*LmxM.31.1110* and the *L. mex* Cas9T7 parental cell line. The plots show the relative read counts across the target gene ORF (marked by grey bar) plus 1.5kb of flanking sequences upstream and downstream of the target gene (CPM-normalised coverage in 50bp bins). Reads from the mutant line are coloured green and the reads from the parental line are coloured pink.

***Supplementary Figure 3. Analysis of chromosome ploidy in selected mutants.***
Whole genome sequencing read coverage for each chromosome was scaled to ploidy by dividing the coverage value for each bin by median coverage/2. The distribution of scaled coverage is plotted for each chromosome for each mutant line (green) and the parental line (pink). Mutant D*LmxM.31.2660*, showed an increase of chromosome 32 from 2n to 3n, mutant D*LmxM.08_29.2780*, showed an increase of chromosome 6 from 2n to 3n and in mutant D*LmxM.03.0500*, chromosome 3 decreased from 3n to 2n. A shift towards lower copy numbers for chromosome 3 was also seen in the knockout lines D*LmxM.08_29.1640*,

D*LmxM.31.1110*, D*LmxM.08_29.2780*, D*LmxM.22.1420* and D*LmxM.32.3040*, when compared to the parental.

***Supplementary Figure 4. Strategy for tandem array knockouts.***
(A) Criteria for deletion of genes in tandem arrays. Genes fitting into the category "Unique" were targeted for array deletion. (B) CRISPR-Cas9 strategy for array replacement using drug-selectable donor DNA cassettes, with homology arms targeting 30 nt upstream of the first gene in the array and 30 nt downstream of the end of the last gene in the array. (C) Loci amplified in diagnostic PCRs to determine whether drug resistance genes are linked to the target gene locus in heterozygous knockouts. (D) Schematic showing types of modified targeted locus in mutants generated utilising 1 or 2 drug selection markers.

***Supplementary Figure 5. Results of tandem array knockouts.***
(A-D) Diagnostic PCRs of heterozygous (Puromycin or Blasticidin resistant) array knockouts, using the strategy shown in Supplementary Figure 4C. Numbers above gel lanes refer to the mutant numbers, as listed in Supplementary Table 1. (A) Diagnostic PCR amplification of each gene from the targeted arrays for knockout validation. Two DNA samples were tested for each GOI: the gDNA of mutants that survived transfection and drug selection (left lane) and *L. mex* Cas9 T7 parental gDNA (right lane). (B) Control PCR amplification of non-target gene from mutant gDNA. From each mutant gDNA sample, a fragment of the Blasticidin resistance cassette was amplified with primers BF1 and BR1 and of a second control gene as depicted in the schematic above the gel image. Gel lanes are numbered as in A. (C) Control PCR amplification of 5' UTR and drug resistance gene from mutant gDNA. This diagnostic PCR amplifies a fragment that links the 5' UTR of the GOI to the integrated drug selection marker thus testing for integration of the donor DNA cassette into the target locus. Gel lanes are numbered as in A. (D) Control PCR amplification of 5' UTR and target gene from mutant gDNA. For single drug resistant array mutants, this diagnostic PCR tests for the retention of the first GOI in the array in the targeted locus. Gel lanes are numbered as in A. (E-F) Diagnostic PCRs of homozygous Puromycin and Blasticidin resistant array knockouts, using the strategy shown in Supplementary Figure 4C. (E) Diagnostic PCR amplification of target genes for knockout validation. Gel lanes are numbered as in A. (F) Control PCR amplification of the Blasticidin resistance gene from mutant gDNA.

***Supplementary Figure 6. Barcode proportions over time for all experiments.***
(A) Barcode proportions in promastigote cultures at 24 h (normalised to 0 h). (B) Trajectories of barcode proportions at 24, 48, 72, 96 and 120 h (normalised to 0 h) in samples from infected macrophages. (C) Trajectories of barcode proportions at 72 h, 3 and 6 weeks

(normalised to 0 h) in samples taken from infected mice. Green, parental controls (SBL1-5); red, control knockout mutants; grey, all other transporter mutants.

***Supplementary Figure 7. Fitness scores of SBL titration.***

TransLeish cell lines ranked in order of their fitness scores in macrophages (A-D) or in mice (E-G). Mutant cell lines are depicted as small grey dots, the parental control cell lines are shown as larger dots, coloured according to the dilution at which they were added to the pool.

### Supplementary Tables Legends

***Supplementary Table 1. TransLeish DB.***

Information about genes targeted in the knockout screen and the resulting mutant cell lines.

***Supplementary Table 2. Diagnostic PCR Results.***

Results of diagnostic PCRs for knockout validations for all mutant cell lines.

***Supplementary Table 3. Tandem Arrays.***

Tab "Table 3A": ID of genes in tandem arrays. Tab "Table 3B": Arrays considered for deletion. Tab "Table 3C": Diagnostic PCR for Blasticidin resistant mutants. Tab "Table 3D": Diagnostic PCR of Puromycin resistant mutants. Tab "Table 3E": Diagnostic PCR of Blasticidin and Puromycin resistant mutants. Tab "Table 3F" - BlastP % protein sequence ID.

***Supplementary Table 4. Fitness Scores and raw read counts.***

Fitness scores, raw read counts and summary of reads for each timepoint from promastigotes growth *in vitro*, hiPSC-Mac infections (Mac 3h, Mac 24h, Mac 48h, Mac 120h) and infection of mice (FP 72h, FP 3w and FP 6w).

***Supplementary Table 5. Barcodes and Primers.***

Barcode and primer sequences used for the generation of mutant cell lines and diagnostic PCRs.

### References to Supplementary Files

Al-Salabi, M.I., Wallace, L.J., and De Koning, H.P. (2003). A *Leishmania major* nucleobase transporter responsible for allopurinol uptake is a functional homolog of the *Trypanosoma brucei* H2 transporter. Mol Pharmacol 63, 814-820.

Aldfer, M.M., AlSiari, T.A., Elati, H.A.A., Natto, M.J., Alfayez, I.A., Campagnaro, G.D., Sani, B., Burchmore, R.J.S., Diallinas, G., and De Koning, H.P. (2022). Nucleoside Transport and Nucleobase Uptake Null Mutants in *Leishmania mexicana* for the Routine Expression and Characterization of Purine and Pyrimidine Transporters. Int J Mol Sci 23.

Araujo-Santos, J.M., Parodi-Talice, A., Castanys, S., and Gamarro, F. (2005). The overexpression of an intracellular ABCA-like transporter alters phospholipid trafficking in *Leishmania*. Biochem Biophys Res Commun 330, 349-355.

Arcari, T., Manzano, J.I., and Gamarro, F. (2017). ABCI3 Is a New Mitochondrial ABC Transporter from *Leishmania major* Involved in Susceptibility to Antimonials and Infectivity. Antimicrob Agents Chemother 61.

Baker, N., Hamilton, G., Wilkes, J.M., Hutchinson, S., Barrett, M.P., and Horn, D. (2015). Vacuolar ATPase depletion affects mitochondrial ATPase function, kinetoplast dependency, and drug sensitivity in trypanosomes. Proc Natl Acad Sci U S A 112, 9112-9117.

Balcazar, D.E., Vanrell, M.C., Romano, P.S., Pereira, C.A., Goldbaum, F.A., Bonomi, H.R., and Carrillo, C. (2017). The superfamily keeps growing: Identification in trypanosomatids of RibJ, the first riboflavin transporter family in protists. PLoS Negl Trop Dis 11, e0005513.

Benaim, G., Garcia-Marchan, Y., Reyes, C., Uzcanga, G., and Figarella, K. (2013). Identification of a sphingosine-sensitive $Ca^{2+}$ channel in the plasma membrane of *Leishmania mexicana*. Biochem Biophys Res Commun 430, 1091-1096.

Bertolini, M.S., Chiurillo, M.A., Lander, N., Vercesi, A.E., and Docampo, R. (2019). MICU1 and MICU2 Play an Essential Role in Mitochondrial Ca(2+) Uptake, Growth, and Infectivity of the Human Pathogen *Trypanosoma cruzi*. mBio 10.

Burchmore, R.J., Rodriguez-Contreras, D., McBride, K., Merkel, P., Barrett, M.P., Modi, G., Sacks, D., and Landfear, S.M. (2003). Genetic characterization of glucose transporter function in *Leishmania mexicana*. Proc Natl Acad Sci U S A 100, 3901-3906.

Cabello-Donayre, M., Orrego, L.M., Herraez, E., Vargas, P., Martinez-Garcia, M., Campos-Salinas, J., Perez-Victoria, I., Vicente, B., Marin, J.J.G., and Perez-Victoria, J.M. (2020).

*Leishmania* heme uptake involves LmFLVCRb, a novel porphyrin transporter essential for the parasite. Cell Mol Life Sci 77, 1827-1845.

Campos-Salinas, J., Cabello-Donayre, M., Garcia-Hernandez, R., Perez-Victoria, I., Castanys, S., Gamarro, F., and Perez-Victoria, J.M. (2011). A new ATP-binding cassette protein is involved in intracellular haem trafficking in *Leishmania*. Mol Microbiol 79, 1430-1444.

Campos-Salinas, J., Leon-Guerrero, D., Gonzalez-Rey, E., Delgado, M., Castanys, S., Perez-Victoria, J.M., and Gamarro, F. (2013). LABCG2, a new ABC transporter implicated in phosphatidylserine exposure, is involved in the infectivity and pathogenicity of *Leishmania*. PLoS Negl Trop Dis 7, e2179.

Capul, A.A., Barron, T., Dobson, D.E., Turco, S.J., and Beverley, S.M. (2007). Two functionally divergent UDP-Gal nucleotide sugar transporters participate in phosphoglycan synthesis in *Leishmania major*. J Biol Chem 282, 14006-14017.

Carter, N.S., Drew, M.E., Sanchez, M., Vasudevan, G., Landfear, S.M., and Ullman, B. (2000). Cloning of a novel inosine-guanosine transporter gene from *Leishmania donovani* by functional rescue of a transport-deficient mutant. J Biol Chem 275, 20935-20941.

Carvalho, S., Barreira da Silva, R., Shawki, A., Castro, H., Lamy, M., Eide, D., Costa, V., Mackenzie, B., and Tomas, A.M. (2015). LiZIP3 is a cellular zinc transporter that mediates the tightly regulated import of zinc in *Leishmania infantum* parasites. Mol Microbiol 96, 581-595.

Castanys-Munoz, E., Alder-Baerens, N., Pomorski, T., Gamarro, F., and Castanys, S. (2007). A novel ATP-binding cassette transporter from *Leishmania* is involved in transport of phosphatidylcholine analogues and resistance to alkyl-phospholipids. Mol Microbiol 64, 1141-1153.

Castanys-Munoz, E., Perez-Victoria, J.M., Gamarro, F., and Castanys, S. (2008). Characterization of an ABCG-like transporter from the protozoan parasite *Leishmania* with a role in drug resistance and transbilayer lipid movement. Antimicrob Agents Chemother 52, 3573-3579.

Chiurillo, M.A., Lander, N., Bertolini, M.S., Vercesi, A.E., and Docampo, R. (2019). Functional analysis and importance for host cell infection of the Ca(2+)-conducting subunits of the mitochondrial calcium uniporter of *Trypanosoma cruzi*. Mol Biol Cell 30, 1676-1690.

Coelho, A.C., Beverley, S.M., and Cotrim, P.C. (2003). Functional genetic identification of PRP1, an ABC transporter superfamily member conferring pentamidine resistance in *Leishmania major*. Mol Biochem Parasitol 130, 83-90.

Colasante, C., Pena Diaz, P., Clayton, C., and Voncken, F. (2009). Mitochondrial carrier family inventory of *Trypanosoma brucei brucei*: Identification, expression and subcellular localisation. Mol Biochem Parasitol 167, 104-117.

Dave, N., Cetiner, U., Arroyo, D., Fonbuena, J., Tiwari, M., Barrera, P., Lander, N., Anishkin, A., Sukharev, S., and Jimenez, V. (2021). A novel mechanosensitive channel controls osmoregulation, differentiation, and infectivity in *Trypanosoma cruzi*. Elife 10.

Dean, S., Marchetti, R., Kirk, K., and Matthews, K.R. (2009). A surface transporter family conveys the trypanosome differentiation signal. Nature 459, 213-217.

do Monte-Neto, R.L., Coelho, A.C., Raymond, F., Legare, D., Corbeil, J., Melo, M.N., Frezard, F., and Ouellette, M. (2011). Gene expression profiling and molecular characterization of antimony resistance in *Leishmania amazonensis*. PLoS Negl Trop Dis 5, e1167.

Docampo, R., and Huang, G. (2015). Calcium signaling in trypanosomatid parasites. Cell Calcium 57, 194-202.

Drew, M.E., Langford, C.K., Klamo, E.M., Russell, D.G., Kavanaugh, M.P., and Landfear, S.M. (1995). Functional expression of a myo-inositol/$H^+$ symporter from *Leishmania* donovani. Mol Cell Biol 15, 5508-5515.

Feng, X., Rodriguez-Contreras, D., Buffalo, C., Bouwer, H.G., Kruvand, E., Beverley, S.M., and Landfear, S.M. (2009). Amplification of an alternate transporter gene suppresses the avirulent phenotype of glucose transporter null mutants in *Leishmania mexicana*. Mol Microbiol 71, 369-381.

Hasne, M.P., and Ullman, B. (2005). Identification and characterization of a polyamine permease from the protozoan parasite *Leishmania major*. J Biol Chem 280, 15188-15194.

Hendrickson, N., Sifri, C.D., Henderson, D.M., Allen, T., Wirth, D.F., and Ullman, B. (1993). Molecular characterization of the ldmdr1 multidrug resistance gene from *Leishmania donovani*. Mol Biochem Parasitol 60, 53-64.

Huang, G., Bartlett, P.J., Thomas, A.P., Moreno, S.N., and Docampo, R. (2013). Acidocalcisomes of *Trypanosoma brucei* have an inositol 1,4,5-trisphosphate receptor that is required for growth and infectivity. Proc Natl Acad Sci U S A 110, 1887-1892.

Huang, G., and Docampo, R. (2018). The Mitochondrial Ca(2+) Uniporter Complex (MCUC) of *Trypanosoma brucei* Is a Hetero-oligomer That Contains Novel Subunits Essential for Ca(2+) Uptake. mBio 9.

Huynh, C., Sacks, D.L., and Andrews, N.W. (2006). A *Leishmania amazonensis* ZIP family iron transporter is essential for parasite replication within macrophage phagolysosomes. J Exp Med 203, 2363-2375.

Huynh, C., Yuan, X., Miguel, D.C., Renberg, R.L., Protchenko, O., Philpott, C.C., Hamza, I., and Andrews, N.W. (2012). Heme uptake by *Leishmania amazonensis* is mediated by the transmembrane protein LHR1. PLoS Pathog 8, e1002795.

Ilg, T., Demar, M., and Harbecke, D. (2001). Phosphoglycan repeat-deficient *Leishmania mexicana* parasites remain infectious to macrophages and mice. J Biol Chem 276, 4988-4997.

Inbar, E., Canepa, G.E., Carrillo, C., Glaser, F., Suter Grotemeyer, M., Rentsch, D., Zilberstein, D., and Pereira, C.A. (2012). Lysine transporters in human trypanosomatid pathogens. Amino Acids 42, 347-360.

Inbar, E., Schlisselberg, D., Suter Grotemeyer, M., Rentsch, D., and Zilberstein, D. (2013). A versatile proline/alanine transporter in the unicellular pathogen *Leishmania donovani* regulates amino acid homoeostasis and osmotic stress responses. Biochem J 449, 555-566.

Ishemgulova, A., Hlavacova, J., Majerova, K., Butenko, A., Lukes, J., Votypka, J., Volf, P., and Yurchenko, V. (2018). CRISPR/Cas9 in *Leishmania mexicana*: A case study of LmxBTN1. PLoS One 13, e0192723.

Jackson, A.P. (2007). Origins of amino acid transporter loci in trypanosomatid parasites. BMC Evol Biol 7, 26.

Jimenez, V., and Docampo, R. (2012). Molecular and electrophysiological characterization of a novel cation channel of *Trypanosoma cruzi*. PLoS Pathog 8, e1002750.

Laranjeira-Silva, M.F., Wang, W., Samuel, T.K., Maeda, F.Y., Michailowsky, V., Hamza, I., Liu, Z., and Andrews, N.W. (2018). A MFS-like plasma membrane transporter required for *Leishmania* virulence protects the parasites from iron toxicity. PLoS Pathog 14, e1007140.

Leprohon, P., Legare, D., Girard, I., Papadopoulou, B., and Ouellette, M. (2006). Modulation of Leishmania ABC protein gene expression through life stages and among drug-resistant parasites. Eukaryot Cell 5, 1713-1725.

Ma, D., Russell, D.G., Beverley, S.M., and Turco, S.J. (1997). Golgi GDP-mannose uptake requires *Leishmania* LPG2. A member of a eukaryotic family of putative nucleotide-sugar transporters. J Biol Chem 272, 3799-3805.

Manzano, J.I., Perea, A., Leon-Guerrero, D., Campos-Salinas, J., Piacenza, L., Castanys, S., and Gamarro, F. (2017). *Leishmania* LABCG1 and LABCG2 transporters are involved in virulence and oxidative stress: functional linkage with autophagy. Parasit Vectors 10, 267.

Martinez-Garcia, M., Campos-Salinas, J., Cabello-Donayre, M., Pineda-Molina, E., Galvez, F.J., Orrego, L.M., Sanchez-Canete, M.P., Malagarie-Cazenave, S., Koeller, D.M., and Perez-Victoria, J.M. (2016). LmABCB3, an atypical mitochondrial ABC transporter essential for *Leishmania major* virulence, acts in heme and cytosolic iron/sulfur clusters biogenesis. Parasit Vectors 9, 7.

Meade, J.C. (2019). P-type transport ATPases in *Leishmania* and *Trypanosoma*. Parasite 26, 69.

Mittra, B., Laranjeira-Silva, M.F., Perrone Bezerra de Menezes, J., Jensen, J., Michailowsky, V., and Andrews, N.W. (2016). A Trypanosomatid Iron Transporter that Regulates Mitochondrial Function Is Required for *Leishmania amazonensis* Virulence. PLoS Pathog 12, e1005340.

Morales Herrera, D.S., Contreras Rodriguez, L.E., Rubiano Castellanos, C.C., and Ramirez Hernandez, M.H. (2020). Identification and sub-cellular localization of a NAD transporter in *Leishmania braziliensis* (LbNDT1). Heliyon 6, e04331.

Mosimann, M., Goshima, S., Wenzler, T., Luscher, A., Uozumi, N., and Maser, P. (2010). A Trk/HKT-type $K^+$ transporter from *Trypanosoma brucei*. Eukaryot Cell 9, 539-546.

Ortiz, D., Sanchez, M.A., Pierce, S., Herrmann, T., Kimblin, N., Archie Bouwer, H.G., and Landfear, S.M. (2007). Molecular genetic analysis of purine nucleobase transport in *Leishmania major*. Mol Microbiol 64, 1228-1243.

Osorio-Mendez, J.F., Tellez, G.A., Zapata-Lopez, D., Echeverry, S., and Castano, J.C. (2023). Sequence analysis of SWEET transporters from trypanosomatids and evaluation of its expression in *Trypanosoma cruzi*. Exp Parasitol 248, 108496.

Ouameur, A.A., Girard, I., Legare, D., and Ouellette, M. (2008). Functional analysis and complex gene rearrangements of the folate/biopterin transporter (FBT) gene family in the protozoan parasite *Leishmania*. Mol Biochem Parasitol 162, 155-164.

Papadopoulou, B., Roy, G., Breton, M., Kundig, C., Dumas, C., Fillion, I., Singh, A.K., Olivier, M., and Ouellette, M. (2002). Reduced infectivity of a *Leishmania donovani* biopterin transporter genetic mutant and its use as an attenuated strain for vaccination. Infect Immun 70, 62-68.

Parodi-Talice, A., Araujo, J.M., Torres, C., Perez-Victoria, J.M., Gamarro, F., and Castanys, S. (2003). The overexpression of a new ABC transporter in *Leishmania* is related to phospholipid trafficking and reduced infectivity. Biochim Biophys Acta 1612, 195-207.

Perez-Victoria, F.J., Gamarro, F., Ouellette, M., and Castanys, S. (2003). Functional cloning of the miltefosine transporter. A novel P-type phospholipid translocase from *Leishmania* involved in drug resistance. J Biol Chem 278, 49965-49971.

Plourde, M., Ubeda, J.M., Mandal, G., Monte-Neto, R.L., Mukhopadhyay, R., and Ouellette, M. (2015). Generation of an aquaglyceroporin AQP1 null mutant in *Leishmania major*. Mol Biochem Parasitol 201, 108-111.

Prole, D.L., and Taylor, C.W. (2013). Identification and analysis of putative homologues of mechanosensitive channels in pathogenic protozoa. PLoS One 8, e66068.

Ramakrishnan, S., Unger, L.M., Baptista, R.P., Cruz-Bustos, T., and Docampo, R. (2021). Deletion of a Golgi protein in *Trypanosoma cruzi* reveals a critical role for $Mn^{2+}$ in protein glycosylation needed for host cell invasion and intracellular replication. PLoS Pathog 17, e1009399.

Richard, D., Kundig, C., and Ouellette, M. (2002). A new type of high affinity folic acid transporter in the protozoan parasite *Leishmania* and deletion of its gene in methotrexate-resistant cells. J Biol Chem 277, 29460-29467.

Rodrigues, C.O., Scott, D.A., and Docampo, R. (1999). Characterization of a vacuolar pyrophosphatase in *Trypanosoma brucei* and its localization to acidocalcisomes. Mol Cell Biol 19, 7712-7723.

Rodriguez-Duran, J., Pinto-Martinez, A., Castillo, C., and Benaim, G. (2019). Identification and electrophysiological properties of a sphingosine-dependent plasma membrane Ca(2+) channel in *Trypanosoma cruzi*. FEBS J 286, 3909-3925.

Rojas, F., Silvester, E., Young, J., Milne, R., Tettey, M., Houston, D.R., Walkinshaw, M.D., Perez-Pi, I., Auer, M., Denton, H., et al. (2019). Oligopeptide Signaling through TbGPR89 Drives Trypanosome Quorum Sensing. Cell 176, 306-317 e316.

Roy, G., Bhattacharya, A., Leprohon, P., and Ouellette, M. (2021). Decreased glutamate transport in acivicin resistant *Leishmania tarentolae*. PLoS Negl Trop Dis 15, e0010046.

Russo-Abrahao, T., Alves-Bezerra, M., Majerowicz, D., Freitas-Mesquita, A.L., Dick, C.F., Gondim, K.C., and Meyer-Fernandes, J.R. (2013). Transport of inorganic phosphate in *Leishmania infantum* and compensatory regulation at low inorganic phosphate concentration. Biochim Biophys Acta 1830, 2683-2689.

Sanchez, M.A. (2013). Molecular identification and characterization of an essential pyruvate transporter from *Trypanosoma brucei*. J Biol Chem 288, 14428-14437.

Sanchez, M.A., Tryon, R., Pierce, S., Vasudevan, G., and Landfear, S.M. (2004). Functional expression and characterization of a purine nucleobase transporter gene from *Leishmania major*. Mol Membr Biol 21, 11-18.

Shaked-Mishan, P., Suter-Grotemeyer, M., Yoel-Almagor, T., Holland, N., Zilberstein, D., and Rentsch, D. (2006). A novel high-affinity arginine transporter from the human parasitic protozoan *Leishmania donovani*. Mol Microbiol 60, 30-38.

Singha, U.K., Sharma, S., and Chaudhuri, M. (2009). Downregulation of mitochondrial porin inhibits cell growth and alters respiratory phenotype in *Trypanosoma brucei*. Eukaryot Cell 8, 1418-1428.

Spath, G.F., Lye, L.F., Segawa, H., Sacks, D.L., Turco, S.J., and Beverley, S.M. (2003). Persistence without pathology in phosphoglycan-deficient *Leishmania major*. Science 301, 1241-1243.

Stafkova, J., Mach, J., Biran, M., Verner, Z., Bringaud, F., and Tachezy, J. (2016). Mitochondrial pyruvate carrier in *Trypanosoma brucei*. Mol Microbiol 100, 442-456.

Steinmann, M.E., Gonzalez-Salgado, A., Butikofer, P., Maser, P., and Sigel, E. (2015). A heteromeric potassium channel involved in the modulation of the plasma membrane potential is essential for the survival of African trypanosomes. FASEB J 29, 3228-3237.

Steinmann, M.E., Schmidt, R.S., Macedo, J.P., Kunz Renggli, C., Butikofer, P., Rentsch, D., Maser, P., and Sigel, E. (2017). Identification and characterization of the three members of the CLC family of anion transport proteins in *Trypanosoma brucei*. PLoS One 12, e0188219.

Yernaux, C., Fransen, M., Brees, C., Lorenzen, S., and Michels, P.A. (2006). *Trypanosoma brucei* glycosomal ABC transporters: identification and membrane targeting. Mol Membr Biol 23, 157-172.

Zhang, W.W., and Matlashewski, G. (2012). Deletion of an ATP-binding cassette protein subfamily C transporter in *Leishmania donovani* results in increased virulence. Mol Biochem Parasitol 185, 165-169.

Zhang, W.W., and Matlashewski, G. (2015). CRISPR-Cas9-Mediated Genome Editing in *Leishmania donovani*. mBio 6, e00861.

Zhu, Y., Davis, A., Smith, B.J., Curtis, J., and Handman, E. (2009). *Leishmania* major CorA-like magnesium transporters play a critical role in parasite development and virulence. Int J Parasitol 39, 713-723.
