## Supplementary Figures 1-7 for "TransLeish: Identification of membrane transporters essential for survival of intracellular *Leishmania* parasites in a systematic gene deletion screen"

### Supplementary Figure 1

#### (A) Diagnostic PCR amplification of target gene for KO validation

**Left lane**, Mutant gDNA
**Right lane**, Parental gDNA (control)

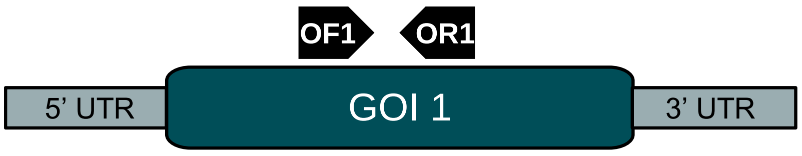


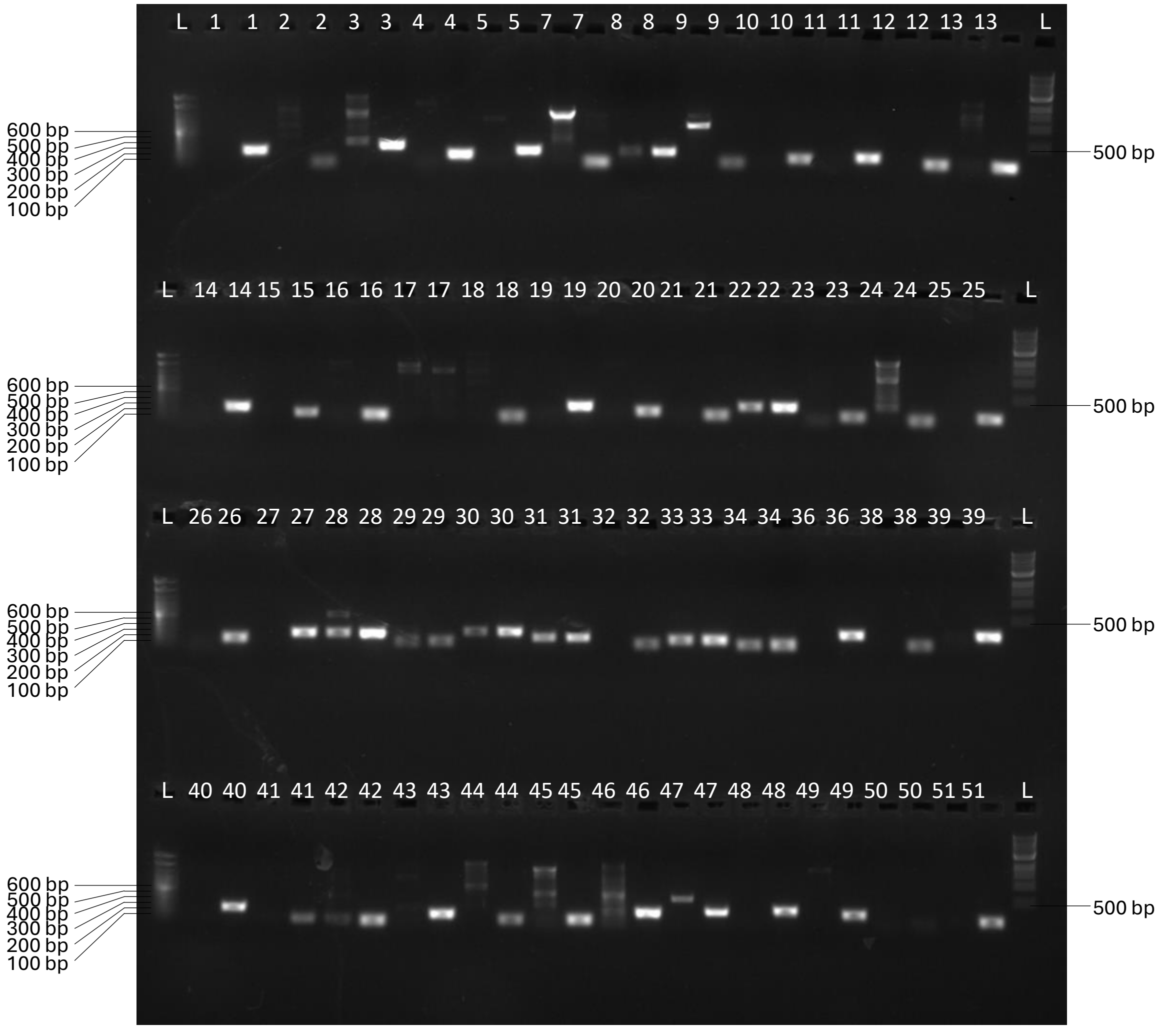


**DNA ladders**
**First** lane, 100 bp DNA Ladder, Ref. 15628050, Invitrogen
**Last** lane, 1 Kb DNA Ladder, Ref. N3232S, NEB

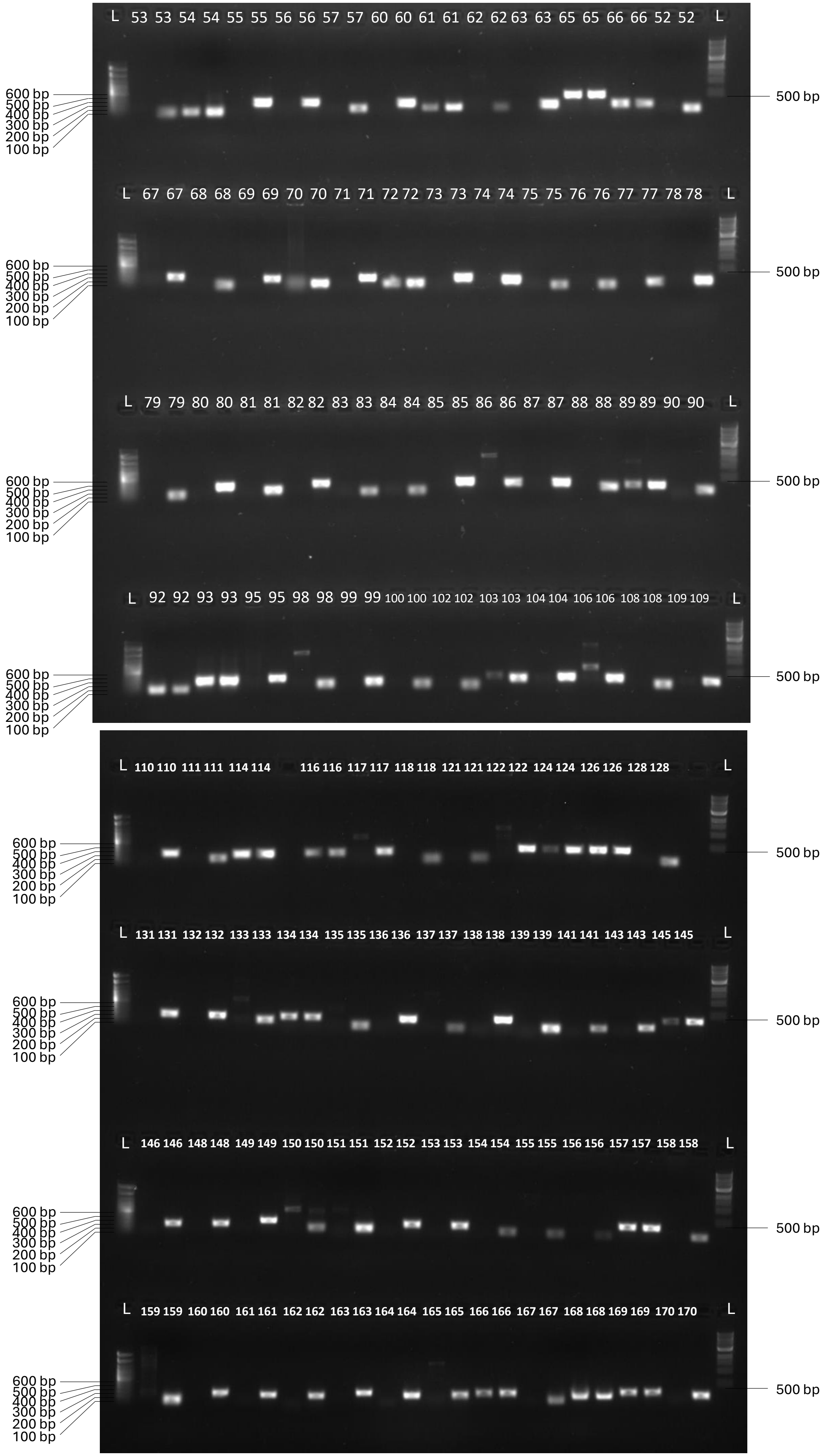
L 53 53 54 54 55 55 56 56 57 57 60 60 61 61 62 62 63 63 65 65 66 66 52 52 L
600 bp
500 bp
400 bp
300 bp
200 bp
100 bp
500 bp
L 67 67 68 68 69 69 70 70 71 71 72 72 73 73 74 74 75 75 76 76 77 77 78 78 L
600 bp
500 bp
400 bp
300 bp
200 bp
100 bp
500 bp
L 79 79 80 80 81 81 82 82 83 83 84 84 85 85 86 86 87 87 88 88 89 89 90 90 L
600 bp
500 bp
400 bp
300 bp
200 bp
100 bp
500 bp
L 92 92 93 93 95 95 98 98 99 99 100 100 102 102 103 103 104 104 106 106 108 108 109 109 L
600 bp
500 bp
400 bp
300 bp
200 bp
100 bp
500 bp
L 110 110 111 111 114 114 116 116 117 117 118 118 121 121 122 122 124 124 126 126 128 128 L
600 bp
500 bp
400 bp
300 bp
200 bp
100 bp
500 bp
L 131 131 132 132 133 133 134 134 135 135 136 136 137 137 138 138 139 139 141 141 143 143 145 145 L
600 bp
500 bp
400 bp
300 bp
200 bp
100 bp
500 bp
L 146 146 148 148 149 149 150 150 151 151 152 152 153 153 154 154 155 155 156 156 157 157 158 158 L
600 bp
500 bp
400 bp
300 bp
200 bp
100 bp
500 bp
L 159 159 160 160 161 161 162 162 163 163 164 164 165 165 166 166 167 167 168 168 169 169 170 170 L
600 bp
500 bp
400 bp
300 bp
200 bp
100 bp
500 bp

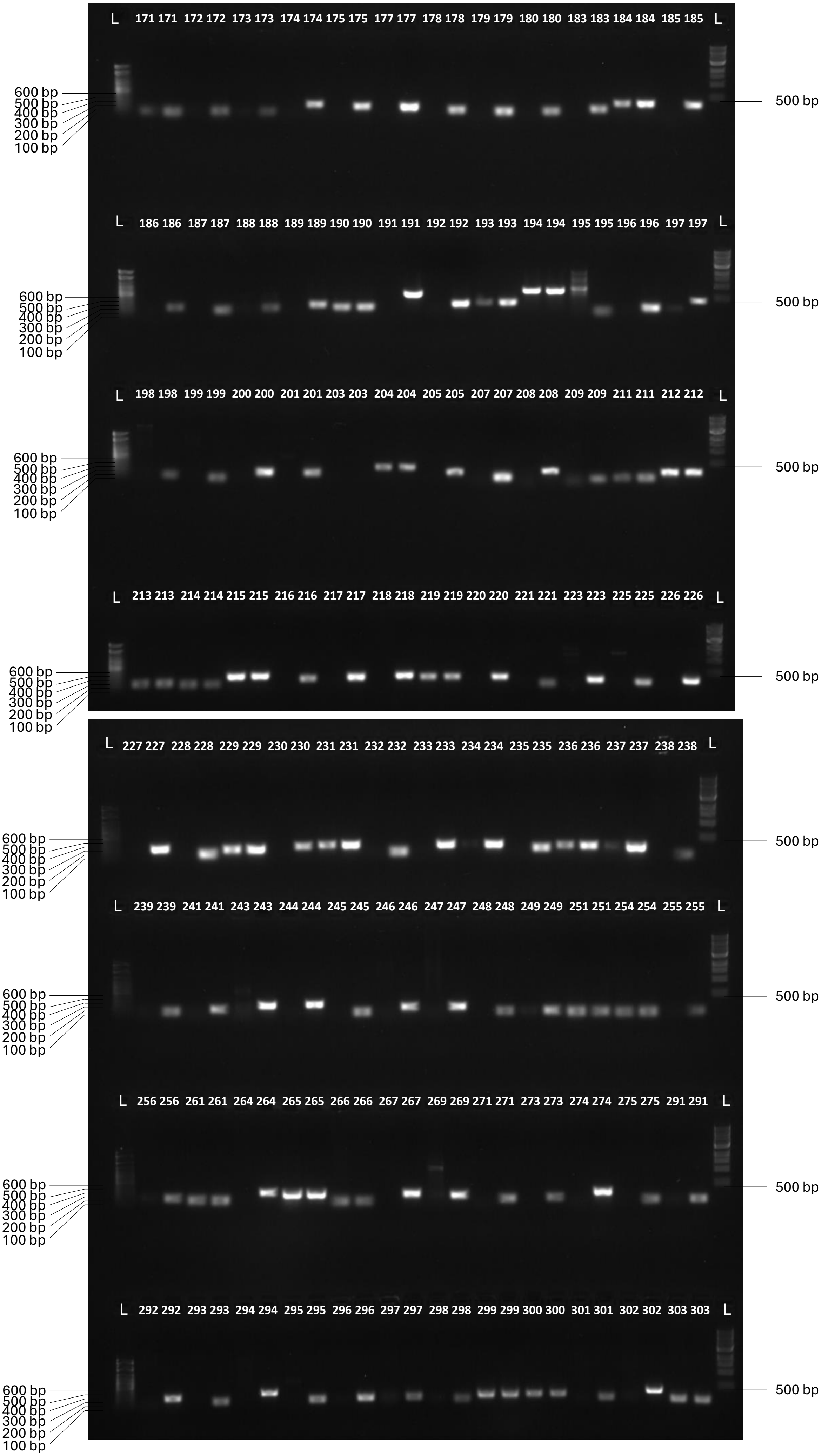
L 171 171 172 172 173 173 174 174 175 175 177 177 178 178 179 179 180 180 183 183 184 184 185 185 L
600 bp
500 bp
400 bp
300 bp
200 bp
100 bp
500 bp
L 186 186 187 187 188 188 189 189 190 190 191 191 192 192 193 193 194 194 195 195 196 196 197 197 L
600 bp
500 bp
400 bp
300 bp
200 bp
100 bp
500 bp
L 198 198 199 199 200 200 201 201 203 203 204 204 205 205 207 207 208 208 209 209 211 211 212 212 L
600 bp
500 bp
400 bp
300 bp
200 bp
100 bp
500 bp
L 213 213 214 214 215 215 216 216 217 217 218 218 219 219 220 220 221 221 223 223 225 225 226 226 L
600 bp
500 bp
400 bp
300 bp
200 bp
100 bp
500 bp
L 227 227 228 228 229 229 230 230 231 231 232 232 233 233 234 234 235 235 236 236 237 237 238 238 L
600 bp
500 bp
400 bp
300 bp
200 bp
100 bp
500 bp
L 239 239 241 241 243 243 244 244 245 245 246 246 247 247 248 248 249 249 251 251 254 254 255 255 L
600 bp
500 bp
400 bp
300 bp
200 bp
100 bp
500 bp
L 256 256 261 261 264 264 265 265 266 266 267 267 269 269 271 271 273 273 274 274 275 275 291 291 L
600 bp
500 bp
400 bp
300 bp
200 bp
100 bp
500 bp
L 292 292 293 293 294 294 295 295 296 296 297 297 298 298 299 299 300 300 301 301 302 302 303 303 L
600 bp
500 bp
400 bp
300 bp
200 bp
100 bp
500 bp

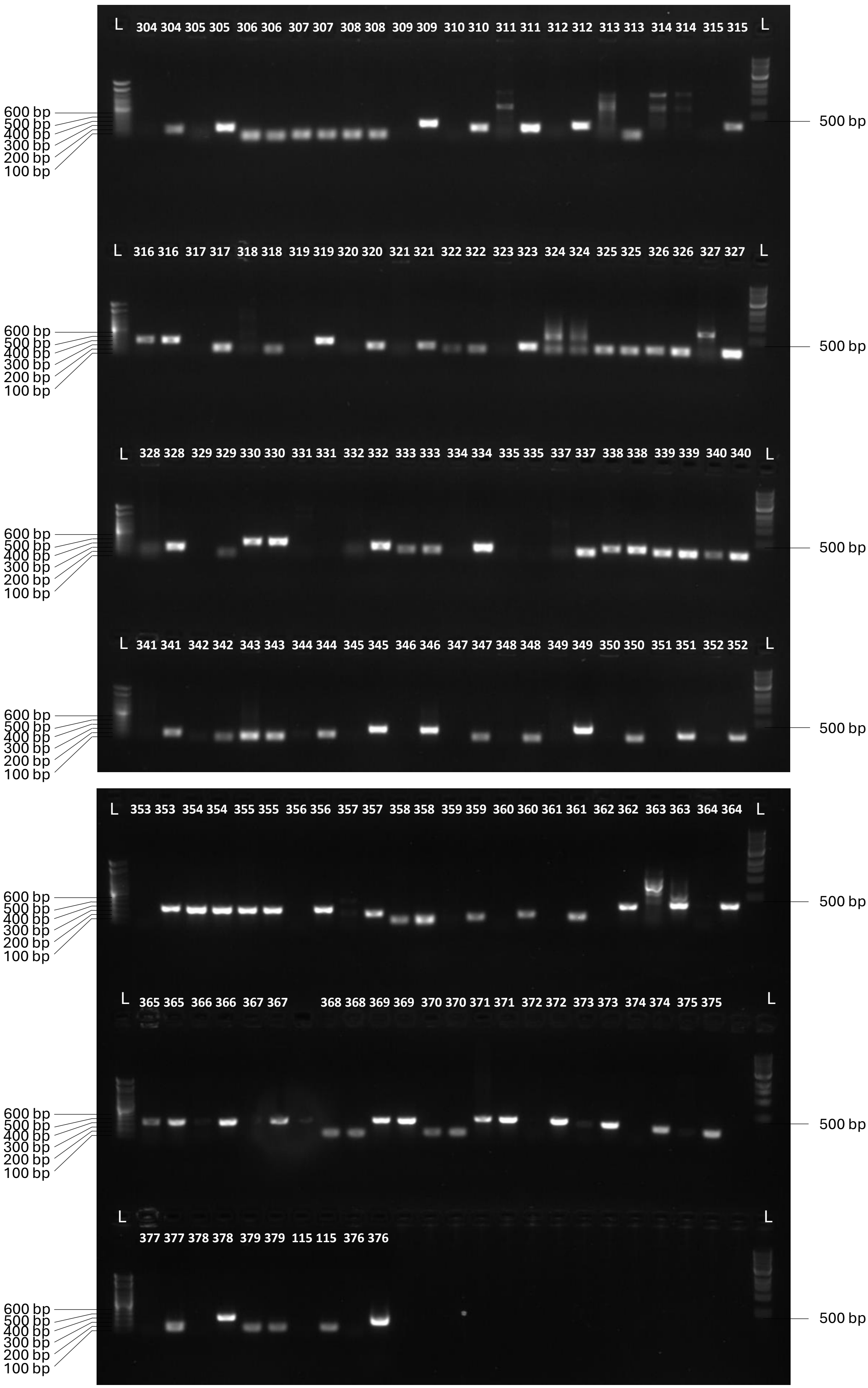
L 304 304 305 305 306 306 307 307 308 308 309 309 310 310 311 311 312 312 313 313 314 314 315 315 L
L 316 316 317 317 318 318 319 319 320 320 321 321 322 322 323 323 324 324 325 325 326 326 327 327 L
L 328 328 329 329 330 330 331 331 332 332 333 333 334 334 335 335 337 337 338 338 339 339 340 340 L
L 341 341 342 342 343 343 344 344 345 345 346 346 347 347 348 348 349 349 350 350 351 351 352 352 L
L 353 353 354 354 355 355 356 356 357 357 358 358 359 359 360 360 361 361 362 362 363 363 364 364 L
L 365 365 366 366 367 367 368 368 369 369 370 370 371 371 372 372 373 373 374 374 375 375 L
L 377 377 378 378 379 379 115 115 376 376 L
600 bp
500 bp
400 bp
300 bp
200 bp
100 bp
500 bp

**(B) Control PCR amplification of a non-target gene from mutant gDNA**

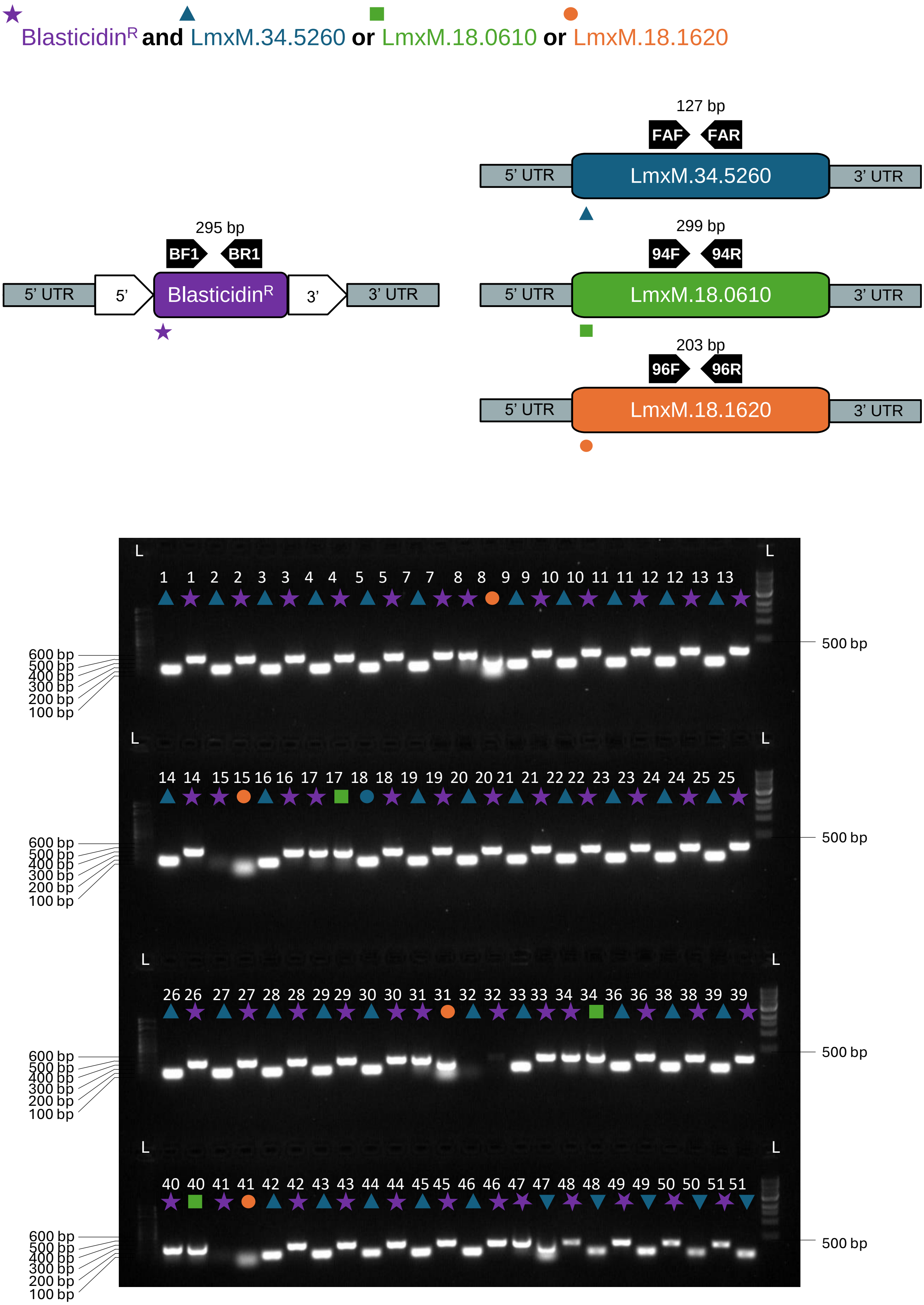

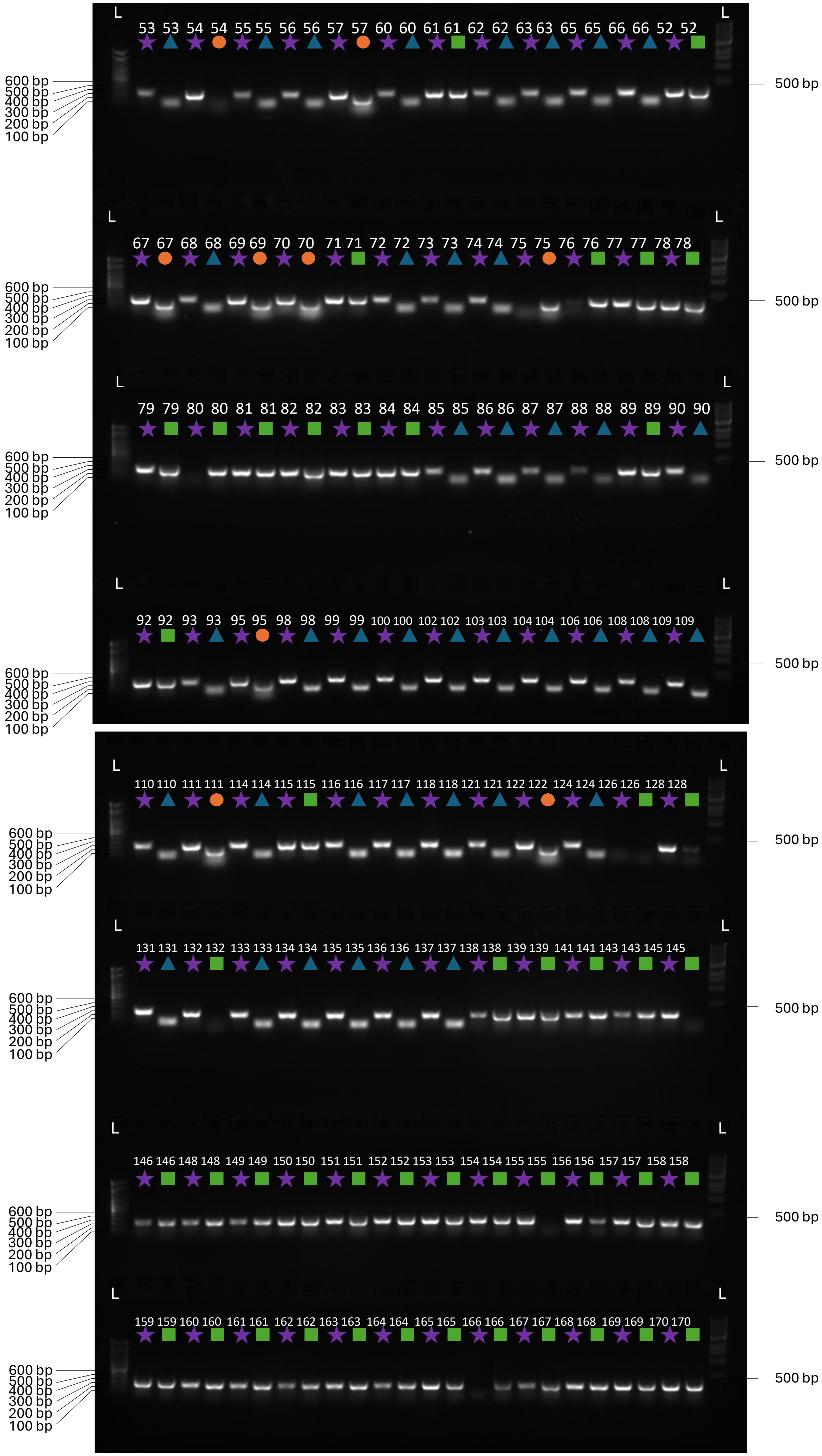
L
53 53 54 54 55 55 56 56 57 57 60 60 61 61 62 62 63 63 65 65 66 66 52 52
L
600 bp
500 bp
400 bp
300 bp
200 bp
100 bp
500 bp
67 67 68 68 69 69 70 70 71 71 72 72 73 73 74 74 75 75 76 76 77 77 78 78
79 79 80 80 81 81 82 82 83 83 84 84 85 85 86 86 87 87 88 88 89 89 90 90
92 92 93 93 95 95 98 98 99 99 100 100 102 102 103 103 104 104 106 106 108 108 109 109
110 110 111 111 114 114 115 115 116 116 117 117 118 118 121 121 122 122 124 124 126 126 128 128
131 131 132 132 133 133 134 134 135 135 136 136 137 137 138 138 139 139 141 141 143 143 145 145
146 146 148 148 149 149 150 150 151 151 152 152 153 153 154 154 155 155 156 156 157 157 158 158
159 159 160 160 161 161 162 162 163 163 164 164 165 165 166 166 167 167 168 168 169 169 170 170

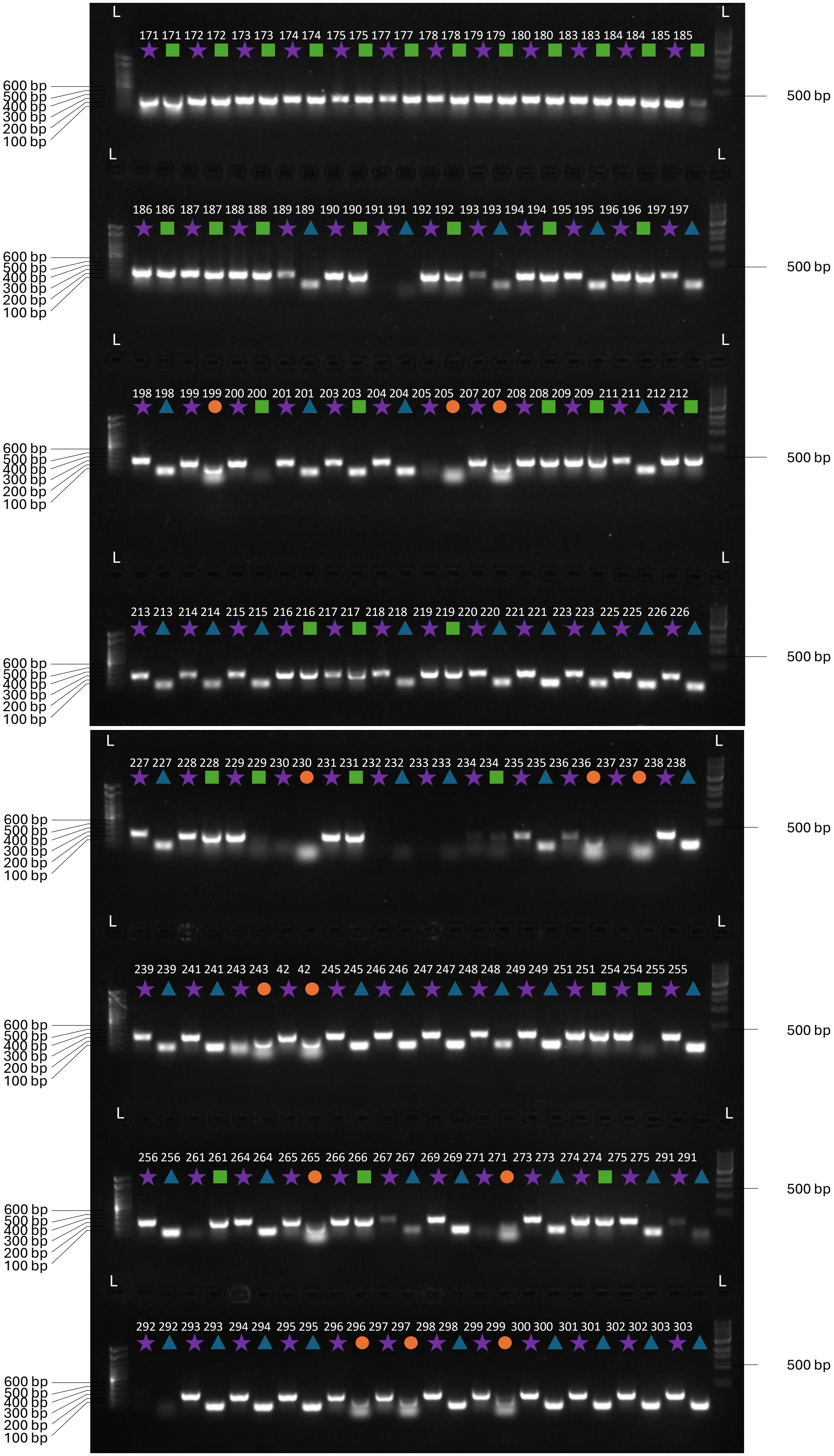

L
171 171 172 172 173 173 174 174 175 175 177 177 178 178 179 179 180 180 183 183 184 184 185 185
600 bp
500 bp
400 bp
300 bp
200 bp
100 bp
500 bp
186 186 187 187 188 188 189 189 190 190 191 191 192 192 193 193 194 194 195 195 196 196 197 197
198 198 199 199 200 200 201 201 203 203 204 204 205 205 207 207 208 208 209 209 211 211 212 212
213 213 214 214 215 215 216 216 217 217 218 218 219 219 220 220 221 221 223 223 225 225 226 226
227 227 228 228 229 229 230 230 231 231 232 232 233 233 234 234 235 235 236 236 237 237 238 238
239 239 241 241 243 243 42 42 245 245 246 246 247 247 248 248 249 249 251 251 254 254 255 255
256 256 261 261 264 264 265 265 266 266 267 267 269 269 271 271 273 273 274 274 275 275 291 291
292 292 293 293 294 294 295 295 296 296 297 297 298 298 299 299 300 300 301 301 302 302 303 303

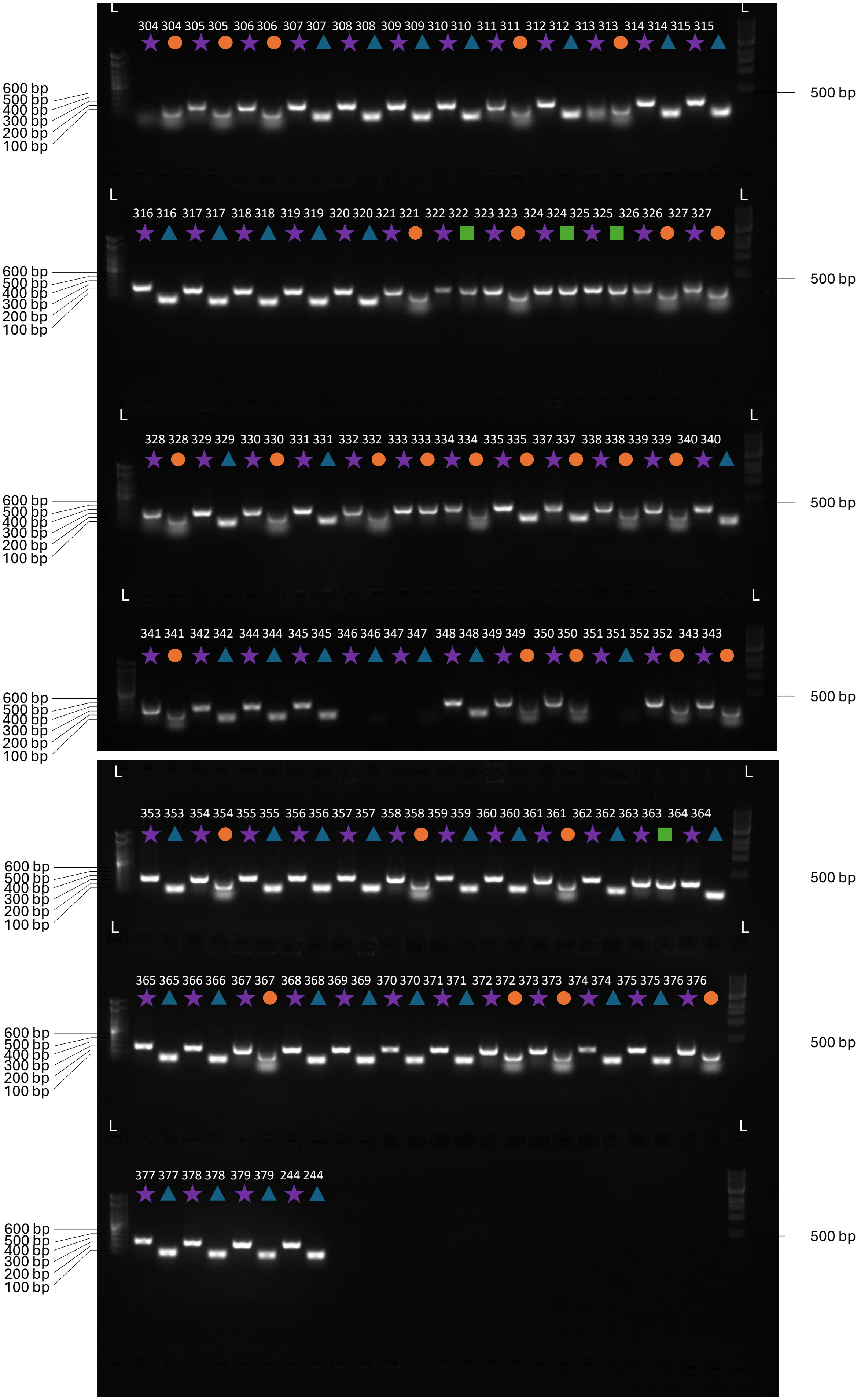
L
304 304 305 305 306 306 307 307 308 308 309 309 310 310 311 311 312 312 313 313 314 314 315 315
600 bp
500 bp
400 bp
300 bp
200 bp
100 bp
500 bp
316 316 317 317 318 318 319 319 320 320 321 321 322 322 323 323 324 324 325 325 326 326 327 327
328 328 329 329 330 330 331 331 332 332 333 333 334 334 335 335 337 337 338 338 339 339 340 340
341 341 342 342 344 344 345 345 346 346 347 347 348 348 349 349 350 350 351 351 352 352 343 343
353 353 354 354 355 355 356 356 357 357 358 358 359 359 360 360 361 361 362 362 363 363 364 364
365 365 366 366 367 367 368 368 369 369 370 370 371 371 372 372 373 373 374 374 375 375 376 376
377 377 378 378 379 379 244 244

**(C) Repeats of diagnostic PCR for selected mutants as in A and B, but using a higher annealing temperature of 60ºC.**

**Left lane**, mutant gDNA
**Right lane**, Parental gDNA (control)

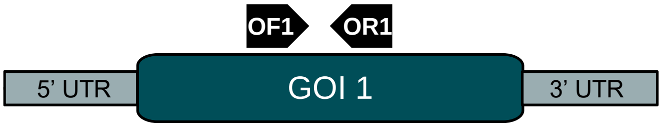


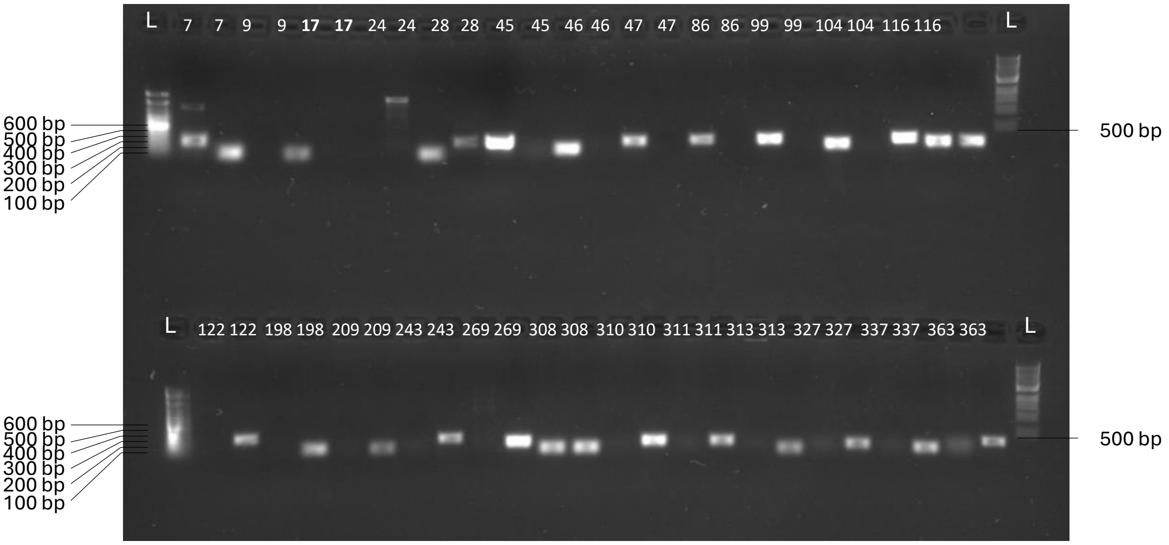


**Control PCR amplification of non-target gene from mutant gDNA**
**Left lane**, Blasticidin$^{R}$
**Right lane**, LmxM.18.0610

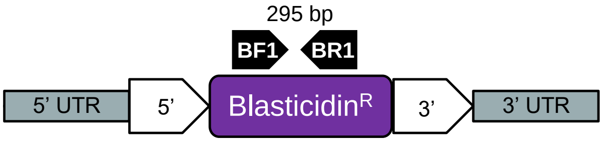


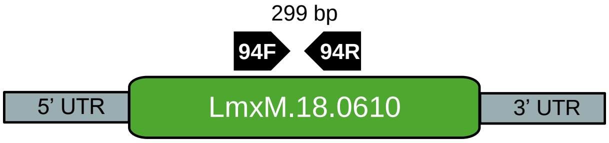


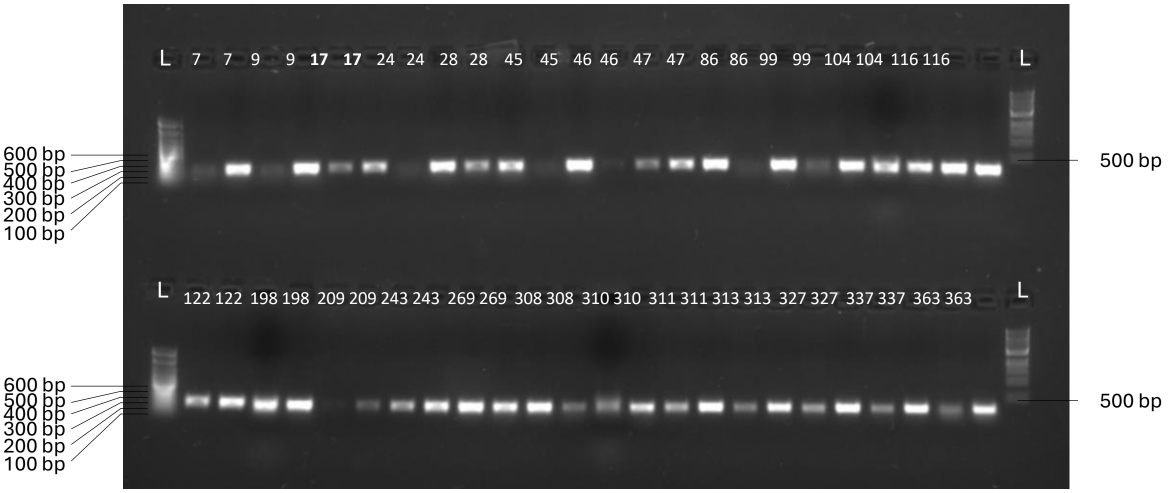

**(D) Repeats of diagnostic PCR for selected mutants where results from the first PCR were unclear, using the same conditions as in (A) and (B).**

**Left lane**, KO gDNA
**Right lane**, Parental gDNA

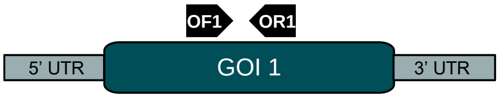


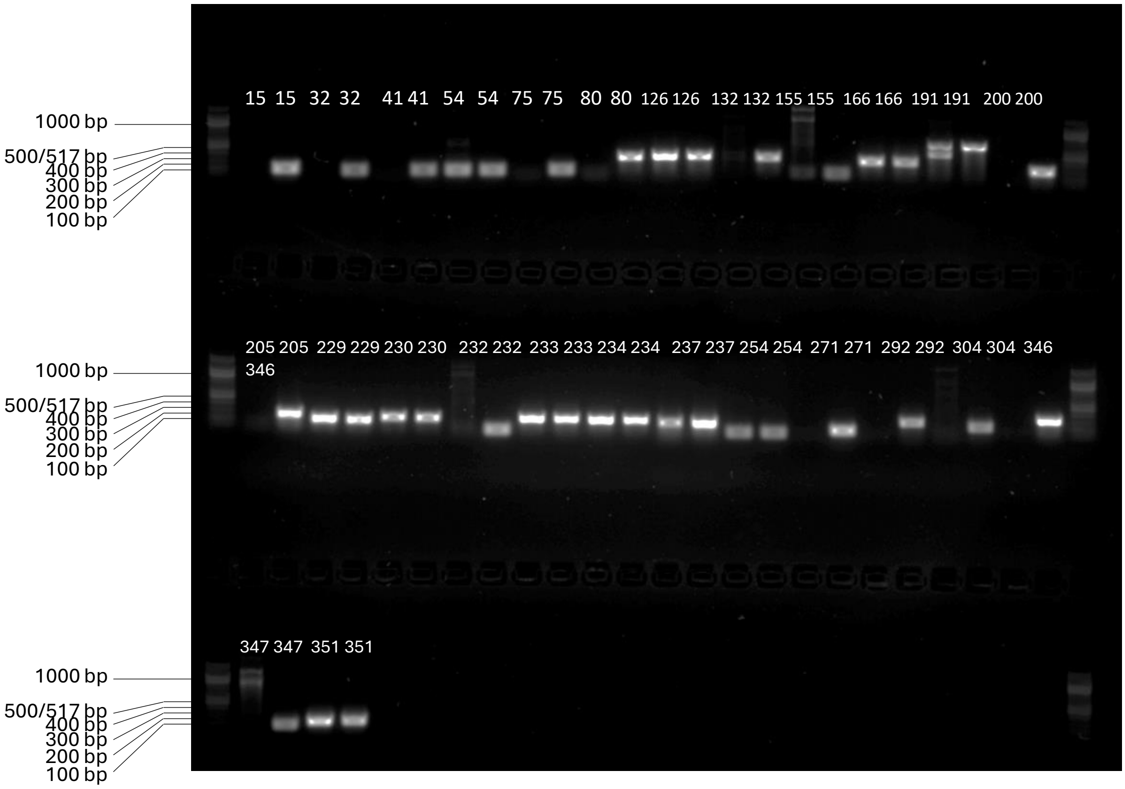


**Control PCR amplification of non-target gene from mutant gDNA**
**Left lane**, Blasticidin$^{R}$
**Right lane**, LmxM.34.5260

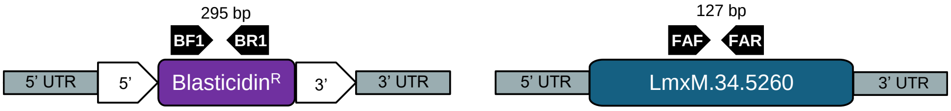


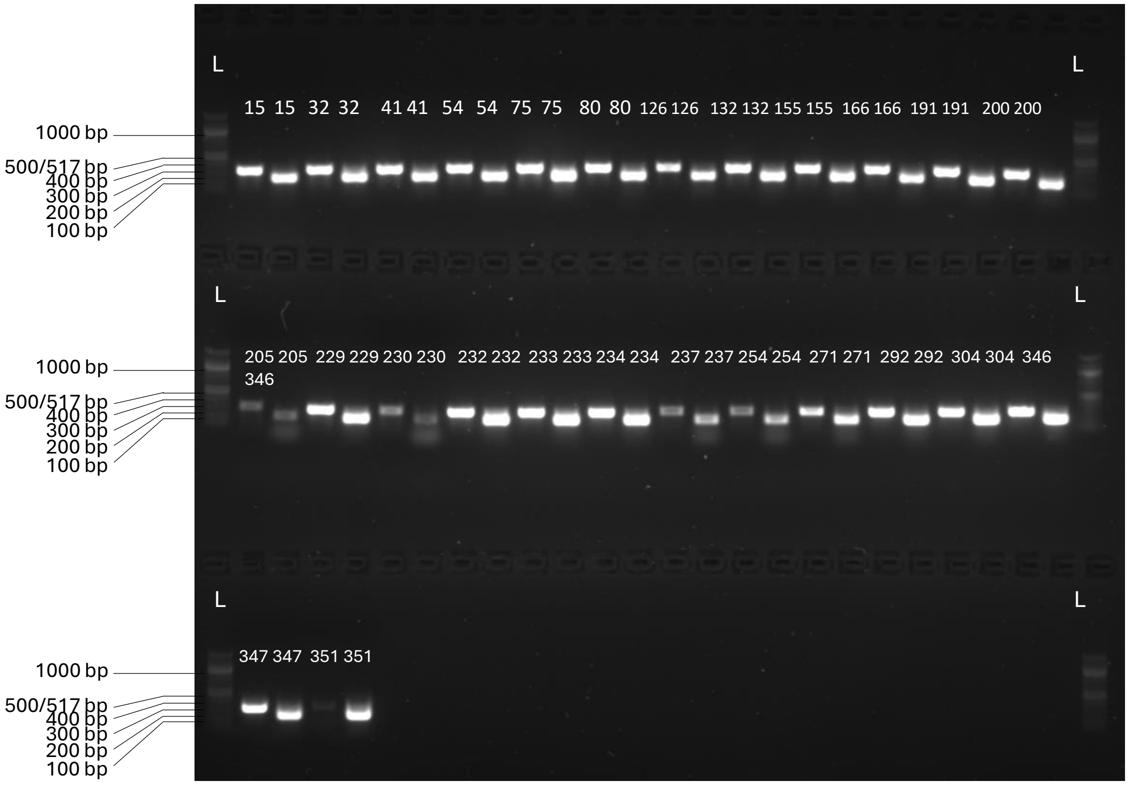


**DNA ladders**
**First** and **last** lanes, Quick-Load Purple 100 bp DNA Ladder, Ref. N0551S, NEB

### Supplementary Figure 2

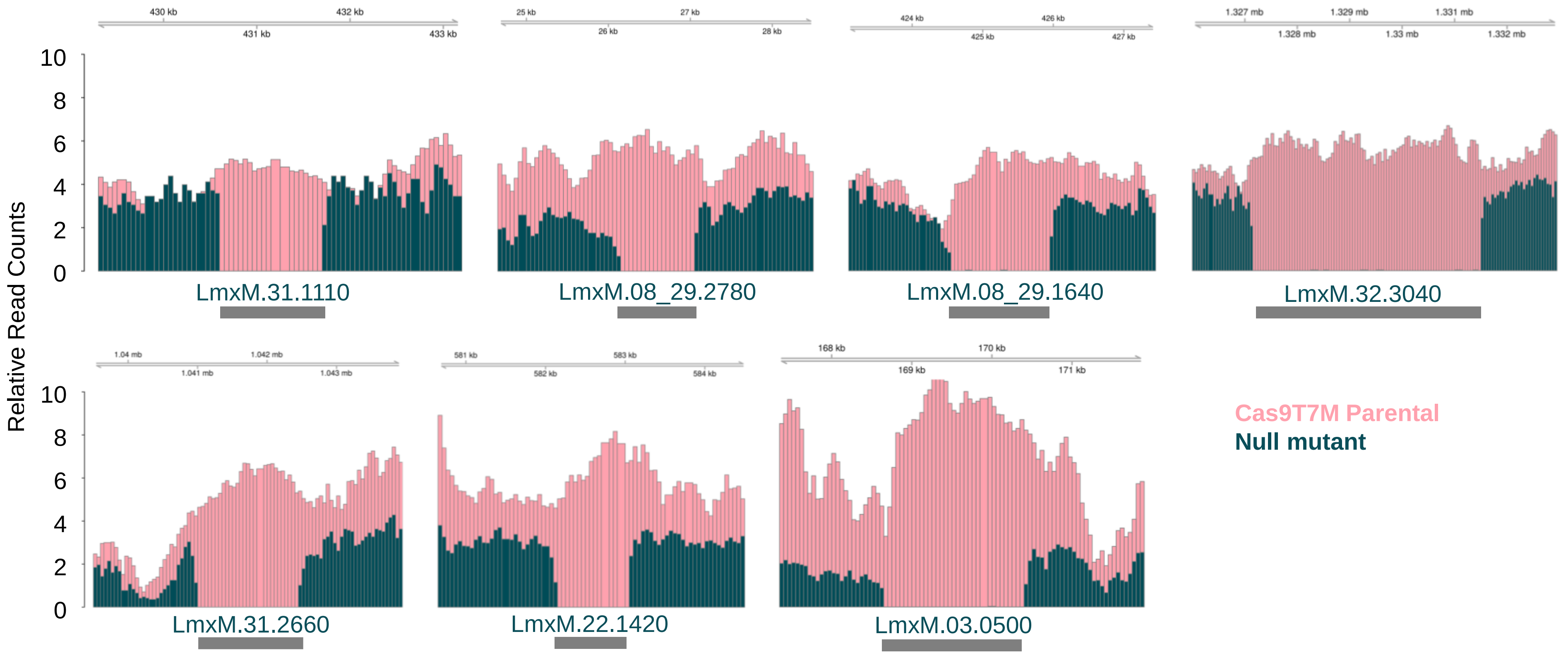

### Supplementary Figure 3

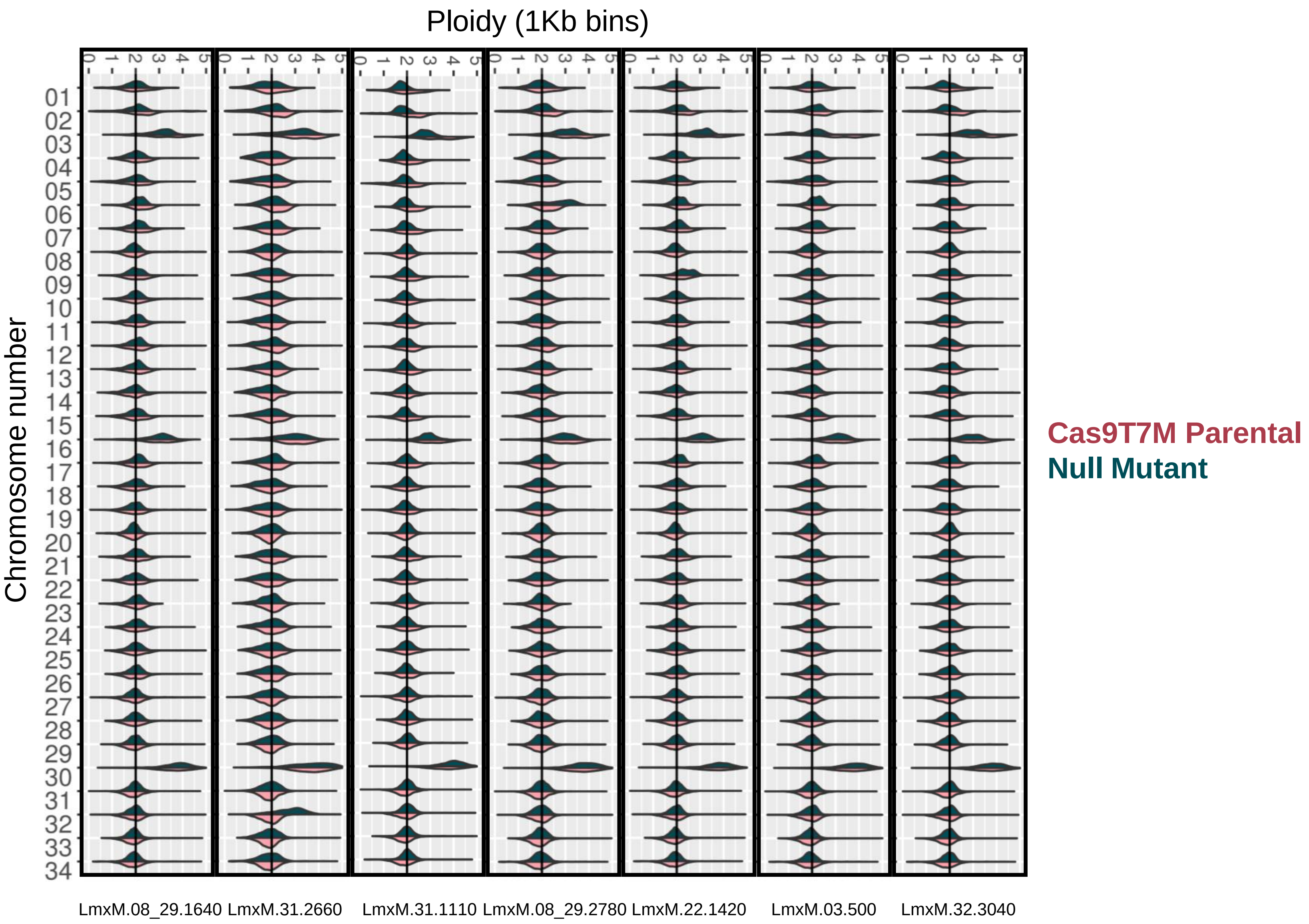

### Supplementary Figure 4

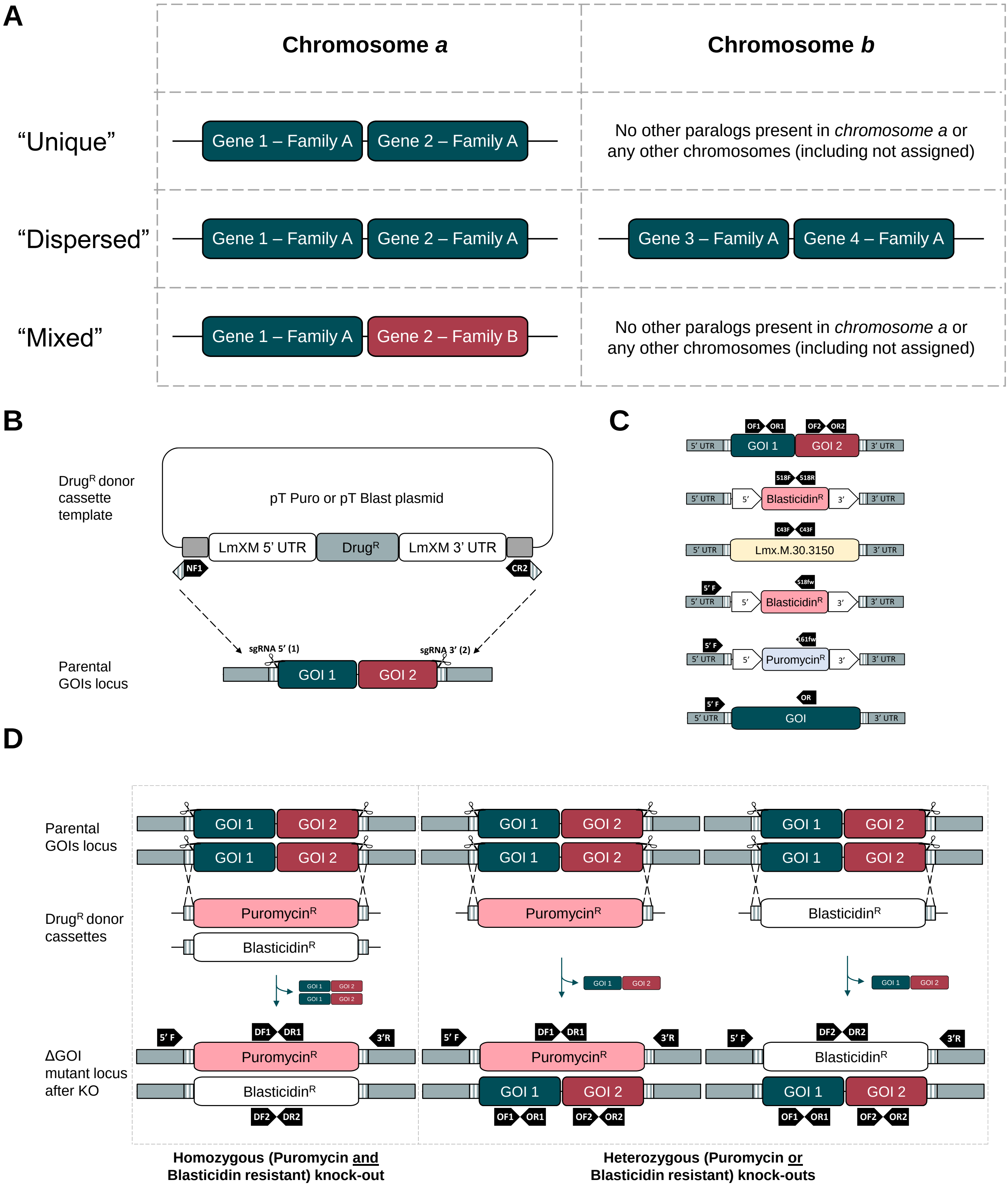

### Supplementary Figure 5

Mutants selected only with Blasticidin or Puromycin (Heterozygous)

**(A) Diagnostic PCR amplification of target genes for KO validation**

**Left lane**, Mutant gDNA

**Right lane**, Parental gDNA (control)

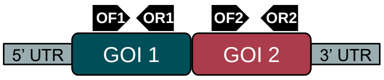


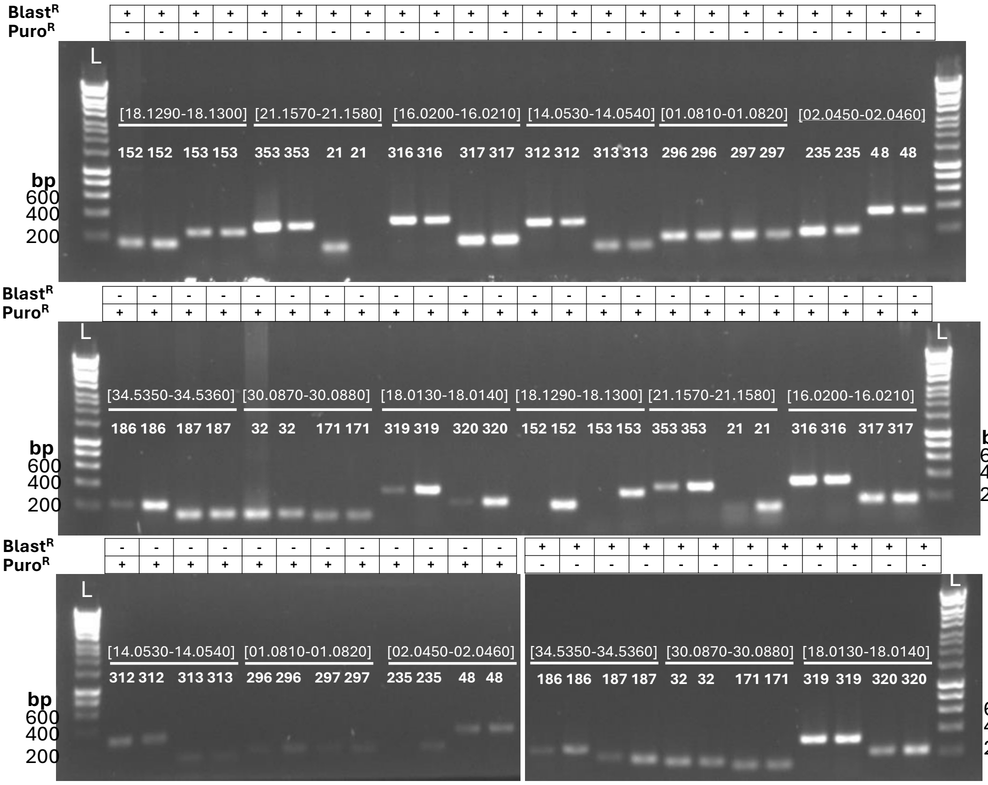


**(B) Control PCR amplification of non-target gene from mutant gDNA**

**Left lane**, Blasticidin$^R$ gene

**Right lane**, CFAB43 (LmxM.30.3150)

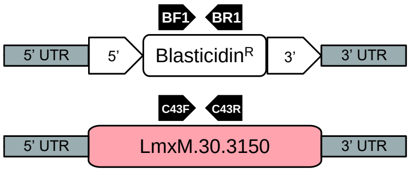


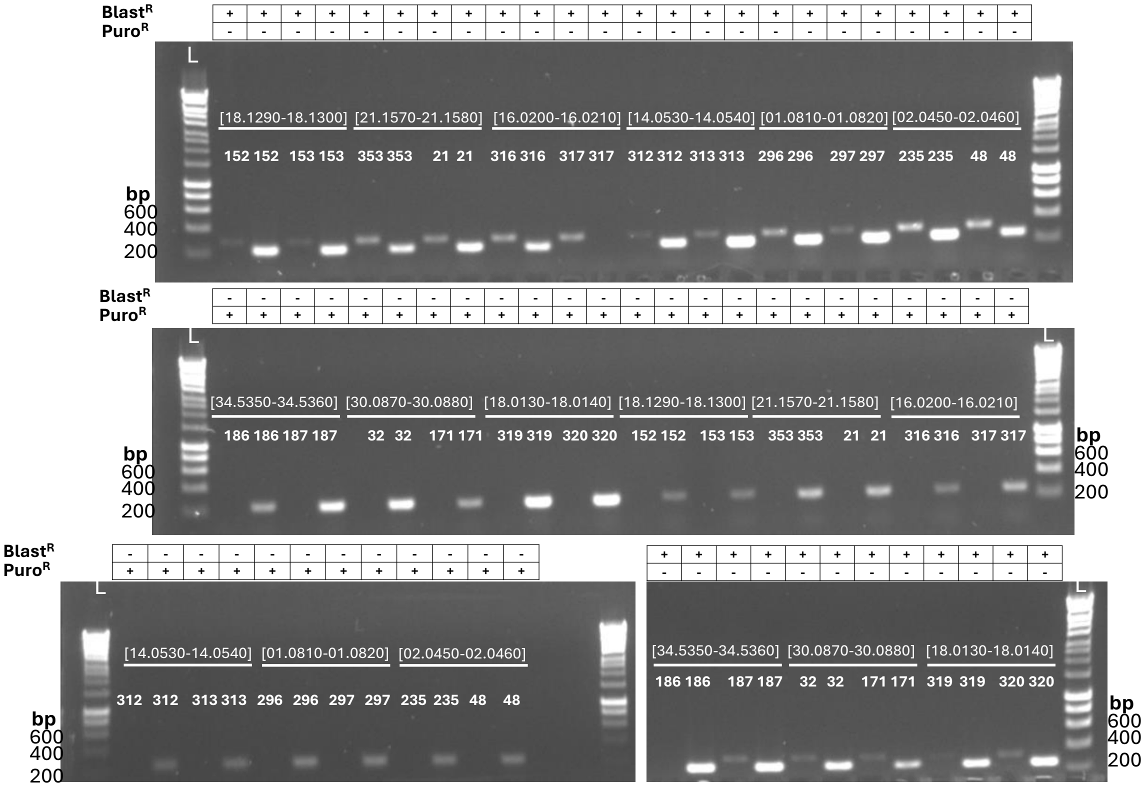

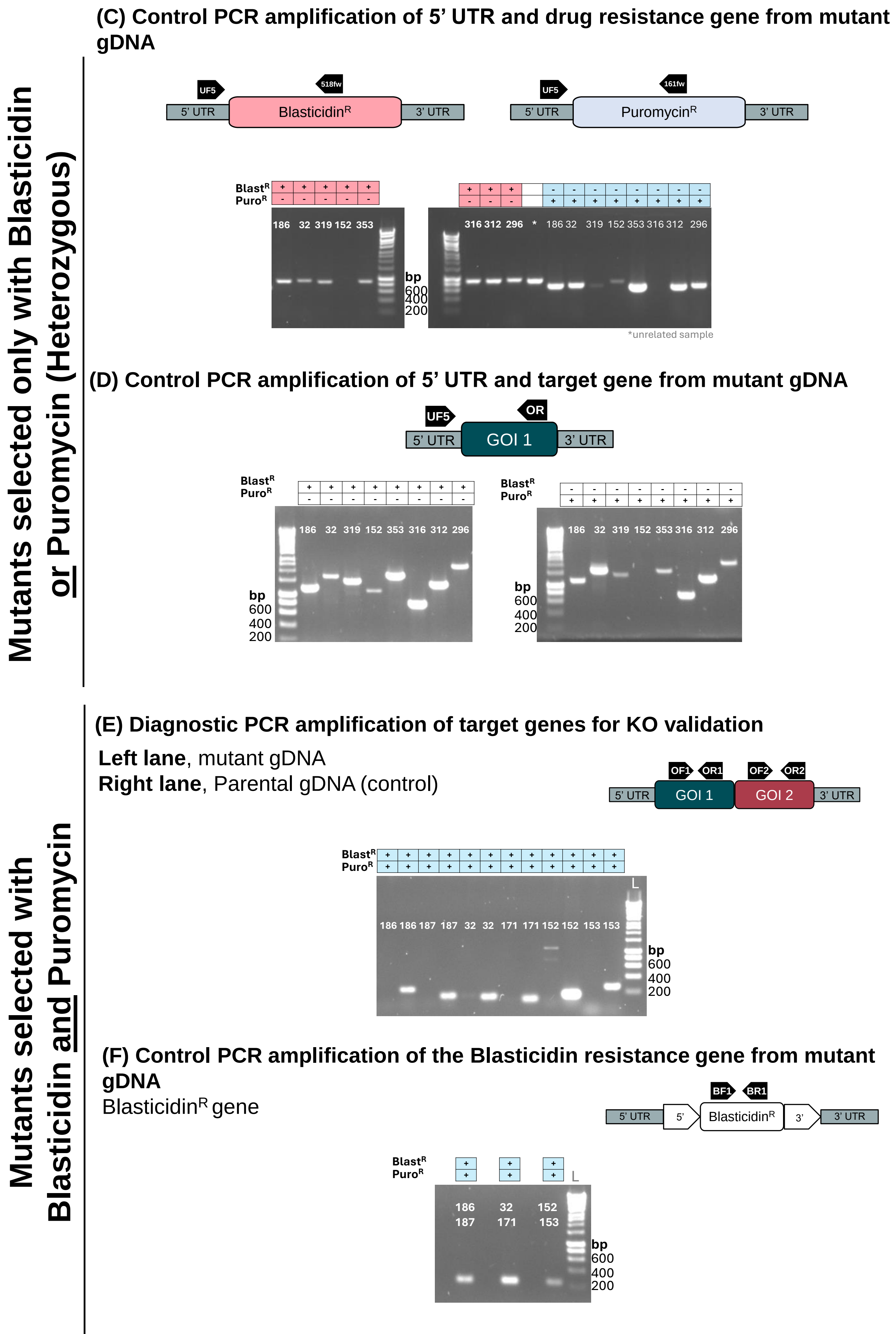
Mutants selected only with Blasticidin or Puromycin (Heterozygous)
(C) Control PCR amplification of 5' UTR and drug resistance gene from mutant gDNA
UF5
518fw
5' UTR
BlasticidinR
3' UTR
161fw
PuromycinR
BlastR
PuroR
186 32 319 152 353
316 312 296 * 186 32 319 152 353 316 312 296
bp
600
400
200
*unrelated sample
(D) Control PCR amplification of 5' UTR and target gene from mutant gDNA
OR
GOI 1
186 32 319 152 353 316 312 296
Mutants selected with Blasticidin and Puromycin
(E) Diagnostic PCR amplification of target genes for KO validation
Left lane, mutant gDNA
Right lane, Parental gDNA (control)
OF1
OR1
OF2
OR2
GOI 2
186 186 187 187 32 32 171 171 152 152 153 153
L
(F) Control PCR amplification of the Blasticidin resistance gene from mutant gDNA
BlasticidinR gene
BF1
BR1
5'
3'
186 187
32 171
152 153

### Supplementary Figure 6

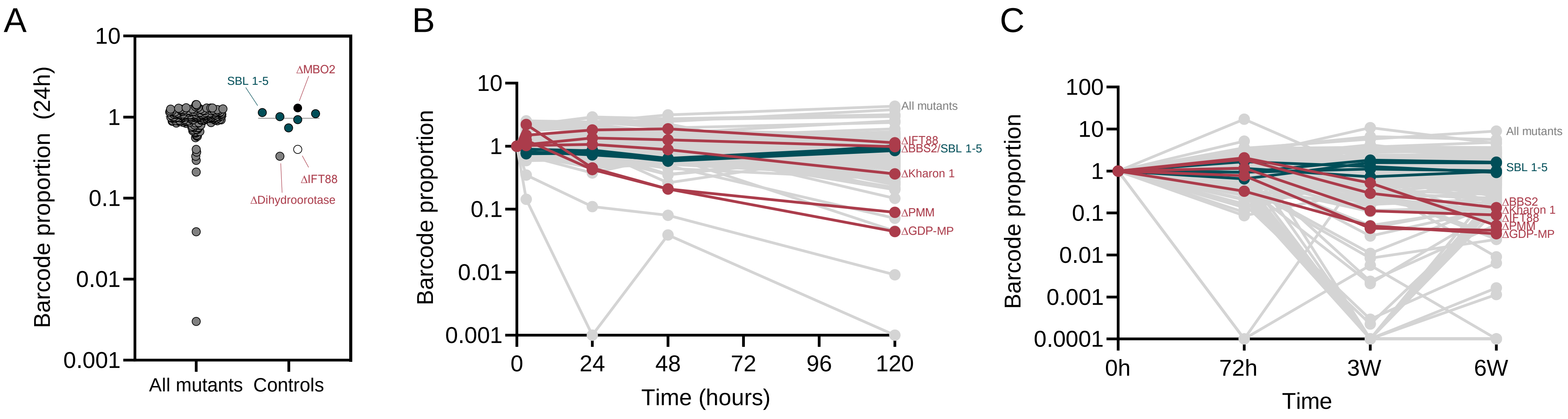

### Supplementary Figure 7

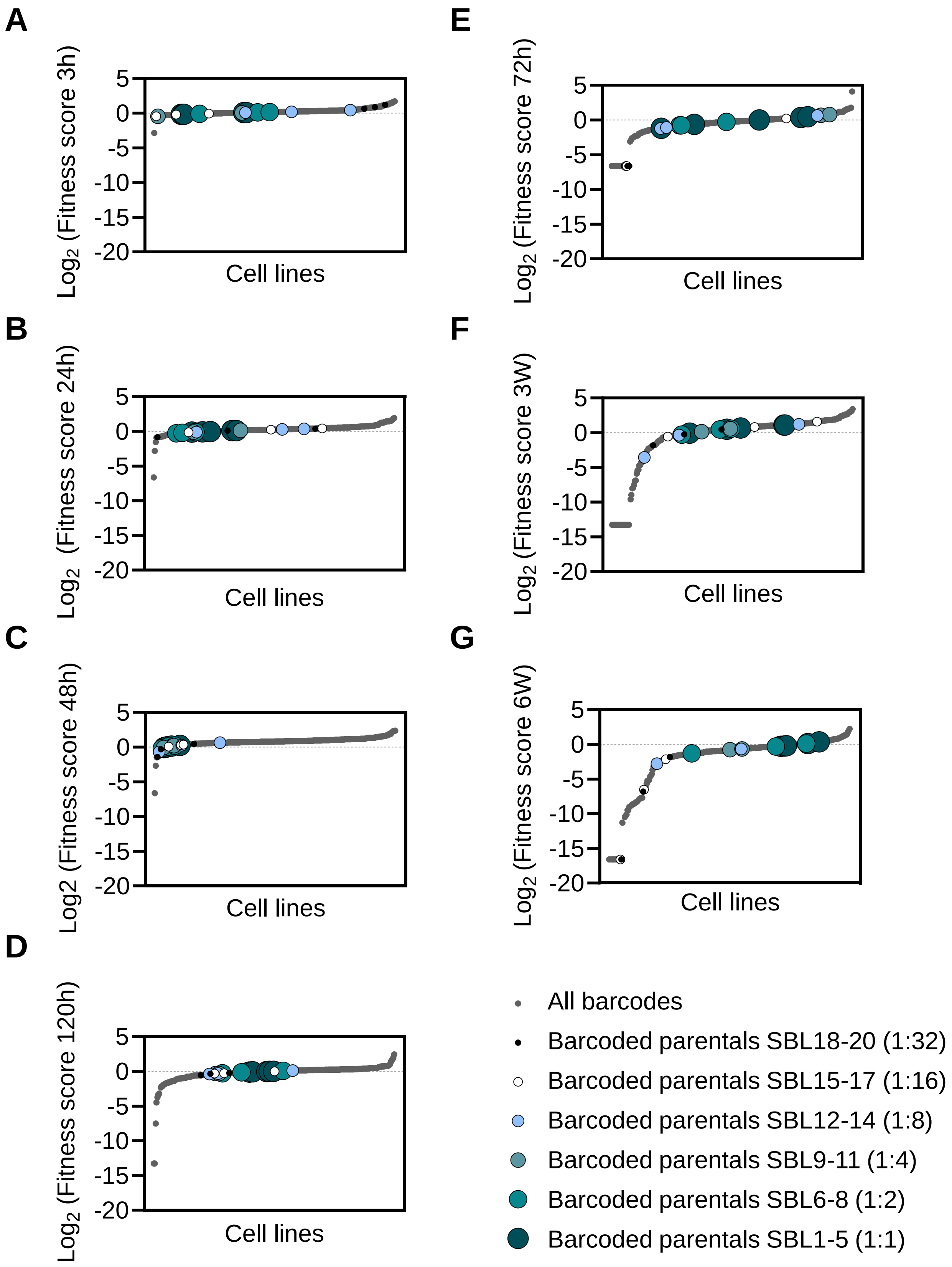
